# MicroRNA-directed control of complex mRNA stability patterns across cell types

**DOI:** 10.1101/2024.10.28.620728

**Authors:** Lukas Oesinghaus, Sebastian Castillo-Hair, Morgan Bean, Nicole Ludwig, Shusruto Rishik, Hao Yuan Kueh, Andreas Keller, Georg Seelig

**Author notes:** These authors contributed equally (Sebastian Castillo-Hair, Morgan Bean).

## Abstract

Limiting expression to target cell types or states is a longstanding goal in gene therapy, which could be met by sensing endogenous microRNA. However, an unclear association between microRNA expression and activity currently hampers such an approach. Here, we probe this relationship by measuring the stability of synthetic microRNA-responsive 3’UTRs across 10 cell lines in a library format. By systematically addressing biases in microRNA expression data and confounding factors such as microRNA crosstalk, we demonstrate that a straightforward model can quantitatively predict the stability of reporters containing full microRNA target sites purely from expression data. We use this model to design constructs with previously unattainable response patterns across our cell lines. We then generalize this approach to primary tissues and cells and show that mRNA can be engineered to be differentially active between rested and exhausted T cell states. The rules we derive for microRNA expression data selection and processing should apply to microRNA-responsive devices for any environment with available expression data.

## Main text

The ability to predictably sense and respond to specific cell states promises to increase the specificity of mRNA, gene, and cell therapies (*1–4*). In principle, cell state-sensing can be achieved by engineering cis-regulatory elements that respond to transcription factors, microRNA, or RNA binding proteins that are differentially expressed in the target cell state. Practically, we usually lack the ability to quantitatively predict sensor function from available prior information, which tends to be limited to expression data for trans-acting RNAs and proteins that serve as inputs to the sensor, making it difficult to generalize beyond cellular environments compatible with iterative experimental testing.

So far, maybe the most widespread approach to transgene targeting relies on microRNA (miRNA) (*5, 6*). miRNA are short (∼22nt) regulatory RNA bound by the RNA-induced silencing complex (RISC), which actively search for and degrade complementary targets, usually in the 3’UTR of mRNAs (*7*). miRNA are attractive for engineering applications because regulation can be achieved simply by inserting target sites complementary to a miRNA into the transgene. Although natural miRNA regulation uses partial target sites, fully complementary target sites confer a stronger regulatory effect. Due to their often cell type-specific expression, miRNA are well-suited to classify cell state via synthetic gene circuits (*8–11*). Most commonly, multiple full target sites for a highly and differentially expressed miRNA are used to exclude transgene expression in a specific tissue (*12–14*).

Still, it remains difficult to achieve complex transgene expression patterns beyond exclusion from a single tissue. Such objectives require multiple target sites for miRNA that are expressed in several tissues but there is no method to quantitatively predict the degree of repression, even for the simple case of full target sites generally used in engineering applications. Previous high-throughput studies have found that a minimum miRNA expression threshold is necessary for effective regulation but found otherwise limited correlation between miRNA expression and target stability (*15, 16*). Smaller studies find low, intermediate, and high correlations (*5, 17–19*). miRNA expression and activity are thus generally believed to be relatively weakly associated (*6, 10, 11*). How multiple miRNA targets combine to produce an overall regulatory effect is similarly contested (*9, 16, 19*). This necessitates individual validation of miRNAs and makes it difficult to anticipate behavior in novel environments (*10*).

Many miRNA sequence-specific features have been discussed to potentially contribute to variability between the activity of individual miRNAs, such as differential AGO sorting (*17*), cleavage activities (*20*), titration by targets on other transcripts (*21, 22*), subcellular localization (*15, 23*), or post-transcriptional modification (*15, 24*). Similarly, target sites can be occluded by secondary structure (*25–27*) or have differing efficiencies based on their proximity to other target sites (*28*). On the other hand, miRNA expression data is well known to be potentially highly biased depending on measurement method, and much effort has been invested into developing less biased measurement protocols (*29–32*). The choice and impact of the used miRNA data is therefore another potentially relevant factor in prediction accuracy.

To address these questions and limitations, we measure the activity of all annotated high-confidence human miRNAs (*33, 34*) across ten different cell lines using reporter libraries. We find that a very simple function quantitatively predicts the stability of reporter genes containing arbitrary combinations of fully complementary miRNA targets in their 3’UTR or 5’UTR purely from readily collectible expression data, but only after identifying high-quality miRNA expression datasets and correcting them for systematic bias. The choice and processing of miRNA expression data is thus found to be by far the most central aspect of predicting the function of full miRNA target sites, while commonly discussed biological differences between individual miRNAs are, with a few exceptions such as crosstalk between highly similar miRNA sequences, almost irrelevant in comparison, at least in the context of fully complementary target sites. Using this model, it is possible to quantitatively design mRNA with defined stability patterns across cell types without the need for individual verification or high-throughput activity measurements, which we demonstrate by designing complex expression patterns across our ten cell lines. Going beyond cell lines, we measure a reporter library in primary mouse CD8+ T cells and show that our model correctly predicts miRNAs capable of distinguishing rested and exhausted T cell states.

## A universal transfer function predicts full miRNA target sites

To measure the regulatory activity of human miRNA, we created reporters for 1,382 miRNAs selected from two major miRNA databases (miRBase (*33*), MirGeneDB (*34*)) (**Fig. 1A**). Single, fully complementary miRNA target sites were inserted into the 3’UTR of a plasmid-based reporter gene (**Fig. 1A, Fig. S1**). The reporter library (10,021 total constructs, most of which will be discussed later) was transiently transfected into 10 cell lines, each of which is characterized by a unique miRNA expression profile. Stability of reporter transcripts in each cell line was inferred from the ratio of mRNA (48h after transfection) to input plasmid counts as determined by sequencing (*35*). The resulting relative stability values are normalized such that the stability for constructs with inactive target sites in the 3’UTR context used (the ‘main context’) is 1.

**Fig. 1.**
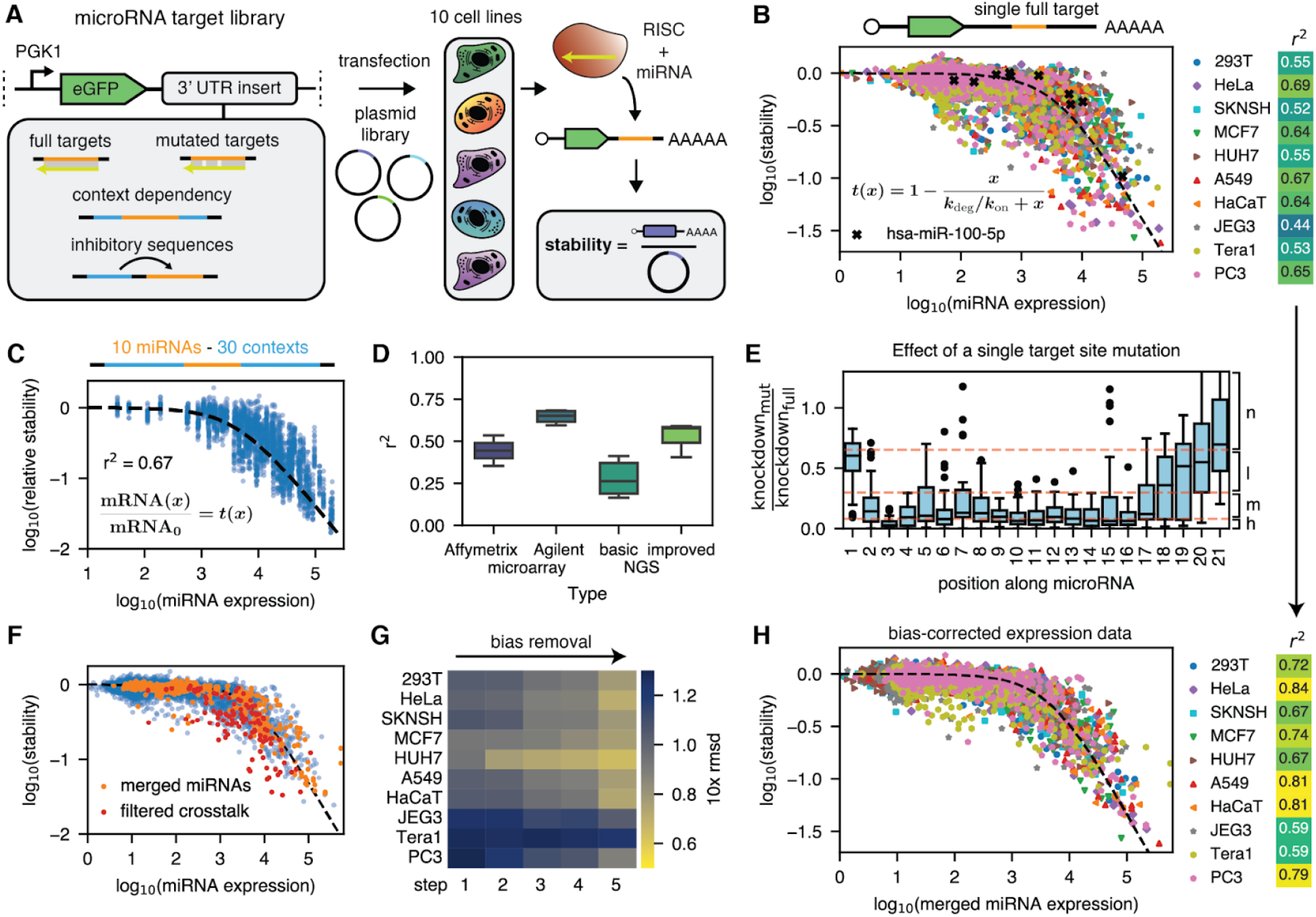
A universal transfer function predicts miRNA activity from expression data. (**A**) Overview of the performed high-throughput stability assay. MicroRNA target sites are ordered as an oligo pool and cloned into a reporter plasmid. The plasmid library is transiently transfected into cell lines, where constructs are degraded by the RNA-induced silencing complex (RISC). Purified mRNA is sequenced. Sequencing counts for mRNA and the plasmid library are used to calculate stability values. (**B**) Stabilities for single full target sites across the different cell lines. The dashed line shows the best fit for the function shown in the inset. The x-axis uses unscaled microarray expression data (*37*). The heatmap shows Pearson r^2^ values between the fit and the measured data. (**C**) The transfer function predicts the relative stability of 3’UTRs containing miRNA targets in different context sequences. (**D**) Pearson r^2^ derived from fitting the transfer function for different miRNA data source types. (**E**) Effect of a single mutation on the relative knockdown between mutated and full target sites for miRNAs where the stability s_full_ of the non-mutated target site is less than 1/3. The knockdown *k* is given by *k* = 1/*s* − 1. n: no l: low, m: medium, h: high impact mutations (**F**) Potential crosstalk was identified based on the mutation classification in (E). The expression of near-identical miRNAs was merged (orange) and other miRNA targets that are likely to experience crosstalk (red) are filtered. (**G**) Changes in the root mean square deviation (rmsd) between measured data and the fit due to expression data bias removal. 1: unscaled microarray data, 2: scaled microarray data, 3: combination of microarray and sequencing data, 4: bias-aware merging, 5: removal of crosstalk. (**H**) Stability for single full target sites using the merged and cross-filtered expression data. The heatmap on the right demonstrates the improvements in the model fit.

Experimental replicates have a Pearson r^2^ above 0.98, suggesting that the assay is highly reproducible (**Fig. S2-S4**). High correlation between mRNA abundance and reporter protein expression measured with flow cytometry is consistent with a model wherein full target sites in the 3’UTR primarily modulate mRNA stability rather than translation (r^2^=0.95, **Fig. S5**)(*36*). To rule out potential biases in our ratiometric stability assay, we separately determined stability from a time course experiment by electroporating our plasmid library into K562 cells, adding actinomycin D after 48h to stop transcription and then collecting multiple mRNA samples over 24h. By fitting the sequence of measured relative abundances to a simple exponential, we quantified mRNA decay rates for the library members (**Fig. S6A-C**). We find that a stability of 1 in the ratiometric stability assay corresponds to a half-life of around 8.9h (**Fig. S6D-E**). We further find a very high correlation (r^2^=0.84) between the time course and ratiometric stability values although we also observe that ratiometric stability values are more spread out than those determined from a time course (**Fig. S6F**).

Our experiments reveal a nearly monotonic, universal relationship between miRNA expression obtained via microarray (*37*) and stability (**Fig. 1B**). As an example, measurements of the miR-100-5p reporter across cell lines reveal varying stability values, and aggregating these across all miRNAs enables mapping the full regulation function. Different miRNAs have varying levels of experimental evidence in databases. The monotonic relationship only clearly holds for miRNAs that are listed as high confidence in miRBase or that are listed in the manually curated MirGeneDB, while low-confidence miRNAs in miRBase sometimes display erroneously (*37*) high measured expression with no activity (**Fig. S7**). We therefore exclude them from the analysis.

The observed relationship between mRNA and miRNA abundance is well explained by a simple transfer function with microRNA concentration as an input and mRNA stability as an output derived under the assumptions of mass action kinetics and instantaneous degradation of the target transcript by the RISC (r^2^ = 0.44-0.69, **Fig. 1B inset, Supplementary Text 1**) (*28, 38*). For low miRNA concentrations ([x]<<k_deg_/k_on_) mRNA degradation is dominated by the baseline degradation rate k_deg_ which in our model captures all miRNA-independent mechanisms for mRNA degradation. In the opposite regime, the degradation rate increases linearly with miRNA concentration. Empirically, mRNA stability starts decreasing at around 1,000 miRNA transcripts per million (tpm), in line with previous results (*15*). To confirm that our observations are not an artifact of the specific reporter context used, we embedded 10 miRNA targets in 30 additional 3’UTR sequences (**Methods**), yielding 3,000 measurements across all cell lines. The transfer function correctly predicts the relative stability due to miRNA target sites across the different contexts (**Fig. 1C**). Different context sequences could occlude target sites by secondary structure formation, thereby causing outliers. We engineered additional context sequences to test this hypothesis and found that, while strong occlusion of the target site strongly reduces miRNA activity (**Fig. S8**), secondary structure in natural 3’UTRs is usually too weak to create strong outliers (**Fig. S9**).

We observe mostly minor but systematic global deviations from the transfer function for some cell lines (**Fig. S10**), likely due to variations in total miRNA levels and the main context stability k_deg_ between cell lines (**Fig. S11**, Methods). A scaling factor of the total miRNA concentration for each cell line accounts for both of these issues and improves the model fit (**Fig. S12**).

### Combining datasets corrects data bias

The universal relationship we observe between miRNA expression and reporter activity seems partially at odds with previous high-throughput experiments (*15, 16*) and in fact, we only observe a universal relationship when using certain miRNA measurements. We hypothesized that the correlation between stability and expression is determined by the method used to collect the miRNA expression data. To test this hypothesis, we collected expression datasets of different origins, including short RNA sequencing (*39–43*) and microarrays from Agilent (*44–47*) and Affymetrix (*48, 49*) and tested their mutual correlation (**Fig. S13A-C**). We additionally include datasets from what we refer to as “improved sequencing methods,” which involve approaches to reduce ligation bias in library construction (*29–31, 50, 51*). High correlation between different collection methods is only observed between Agilent microarray data and improved sequencing methods. We hypothesize that cross-method correlation signifies correctness: We calculated r^2^ values for our transfer function (**Fig. 1D** and **S13D**) and found consistently high correlation even for older publications using Agilent microarray data (r^2^∼0.62) and for improved sequencing methods (r^2^∼0.52). In contrast, standard small RNA sequencing datasets perform poorly (r^2^ 0.11 to 0.41).

Even the highest quality microarray and sequencing data still suffer from technology-specific biases in which certain miRNAs are over- or undercounted. To identify putative technology-specific outliers we combine microarray data (*37*) with improved sequencing data. Because the sample preparation shares little in common, technical biases between these two different types of measurements should not correlate. Biased data points are defined as outliers from the transfer function in one but not the other dataset at least four cell lines (**Fig. S14**).

Because technology-specific bias depends only on miRNA sequence features, we only identify these consistent outliers across multiple cell lines as genuine biased data points. Bias-aware dataset merging is performed by using the geometric mean except in cases of technology-specific bias, where the data from the other technology is used. Dataset merging removes around a third of outliers with a more than two-fold difference to the predicted stability value. Because biases are technology-specific, this identification of false positives and negatives using reporter assays in cell lines should be transferable to more difficult systems such as tissues.

### Crosstalk between miRNA family members explains outliers from ideal behavior

MicroRNAs occur in families with highly similar sequences, leading to potential crosstalk (**Fig. S15**). Because of this high sequence similarity, a microRNA can sometimes degrade a reporter transcript carrying targets designed for other family members. Such crosstalk can result in apparent deviation from the transfer function model. To quantify crosstalk, we measured a reporter library of 1,096 partially mutated miRNA target sites (**Fig. S16A**). Results for individual mismatches qualitatively agree with previous observations (**Fig. 1E, S16B-C**): An adenine at position 1 does not act as a mismatch (*52*). The strongest effects are observed early in the seed and around the cleavage site at positions 10 and 11 (*53, 54*), while mutations beyond position 17-21 have little effect (*55*). Wobble base pairs are generally more tolerated than genuine mismatches but are still damaging (*54*).

We then used these measurements to build a model to predict the impact of multiple mutations. We classify the impact of individual mutations as high, medium, low and none based on their position only and whether the mutation causes a mismatch or a wobble base pair with the miRNA. Targets with multiple mutations are then grouped by the number and impact class of their individual mutations (**Fig. S16D-E**). Activity rapidly falls off with increasing numbers of mutations; multiple mutations can abrogate cleavage even if no mutation individually has a substantial effect (**Fig. S16E, S17A**). We used the classification of individual mutations to build a regression tree model that predicts the effect of multiple mutations (**Fig. S17B**). The model predicts the impact of mutations on a held-out miRNA (**Fig. S17C**), although the effect of mutations is highly sequence-specific (**Fig. S17D**).

Returning to the problem of bias correction, we use these insights to remove the effect of crosstalk from our data. For this, we merge miRNAs that are identical for the first 18 bases by adding their expressing levels and then use the mutation classification (Methods) to filter target sites that are likely to experience substantial crosstalk (**Fig. 1F**), removing many of the strongest outliers.

### Bias correction yields an updated model

Several additional sources of bias and noise can largely be excluded: Low counts in the stability data, miRNA GC content, non-canonical poly(A)-signals, and miRNA length have little to no effect (**Fig. S18A-D**). Some miRNA targets contain homopolymer stretches but except for miR-3613-3p, which contains ten consecutive As in its target site, these do not impact reporter stability (**Fig. S18E**). Variability in miRNA expression within a cell line depending on the source and experimental conditions is another potential source of bias. We therefore collected improved microRNA sequencing data using the same total RNA extracted during the stability assay for HeLa, JEG3, and Tera1 cells. However, there was no improvement in the stability prediction using this data over the previously collected data (**Fig. S18F**).

Weak seed-pairing stability or a high abundance of seed target sites in the transcriptome are two additional miRNA sequence-specific features that have been reported to reduce targeting efficiency (*21, 22*). We calculated the thermodynamic seed-pairing stability associated with each microRNA target site and compared this to the deviation from the fit for expressed miRNAs (**Fig. S19A-B**). Although we find a negative correlation (r=-0.29), this is entirely driven by two sets of closely related miRNA target sites (r=-0.05 if excluded), indicating that seed-pairing stability is not a strong factor for the vast majority of sites. Similarly, we find a decrease in targeting efficiency of around 30% higher relative stability for (rare) very high seed target site abundances (>4000 counts across all human 3’UTRs). Neither effect is substantial or common enough to warrant inclusion in our simplified prediction model.

Each step of bias removal improves the model fit (**Fig. 1G, Fig. S20**). The effect of scaling miRNA concentrations is minor for all cell lines except HUH7 and PC3. Merging data sources and addressing crosstalk substantially improves the fit for almost all cell lines. Tera1 and Jeg3 improve the least, potentially indicating additional sources of variation in these cell lines. Bias removal substantially improves the agreement between the transfer function and measured stabilities (**Fig. 1H**). We call the model without and with scaling factors and bias-aware merging the “baseline model” and the “updated model.”

### Multiple target sites are described by an additive model of miRNA concentrations

Next, we investigated how multiple target sites cooperate to control reporter stability. Multiple models of miRNA interactions have been suggested in the literature (*9, 16, 56*). Here, we focus on the additive model, which assumes that target sites are independent of each other, because it arises naturally from the same differential equations that yielded our transfer function (**Fig. 2A, Supplementary Text 1**). We chose 100 miRNAs spanning a range of expression values and created reporters by repeating their target sites two to six times. In the additive model, the behavior should be predicted by multiplying the miRNA expression data by the repeat number and using the transfer function to predict stability. This approach yields accurate predictions (r^2^∼0.8) with similar accuracy for all target copy numbers (**Fig. 2B, S21-S22**). An alternative model which assumes that regulation is independent of the number of repeats leads to systematic overprediction.

**Fig. 2.**
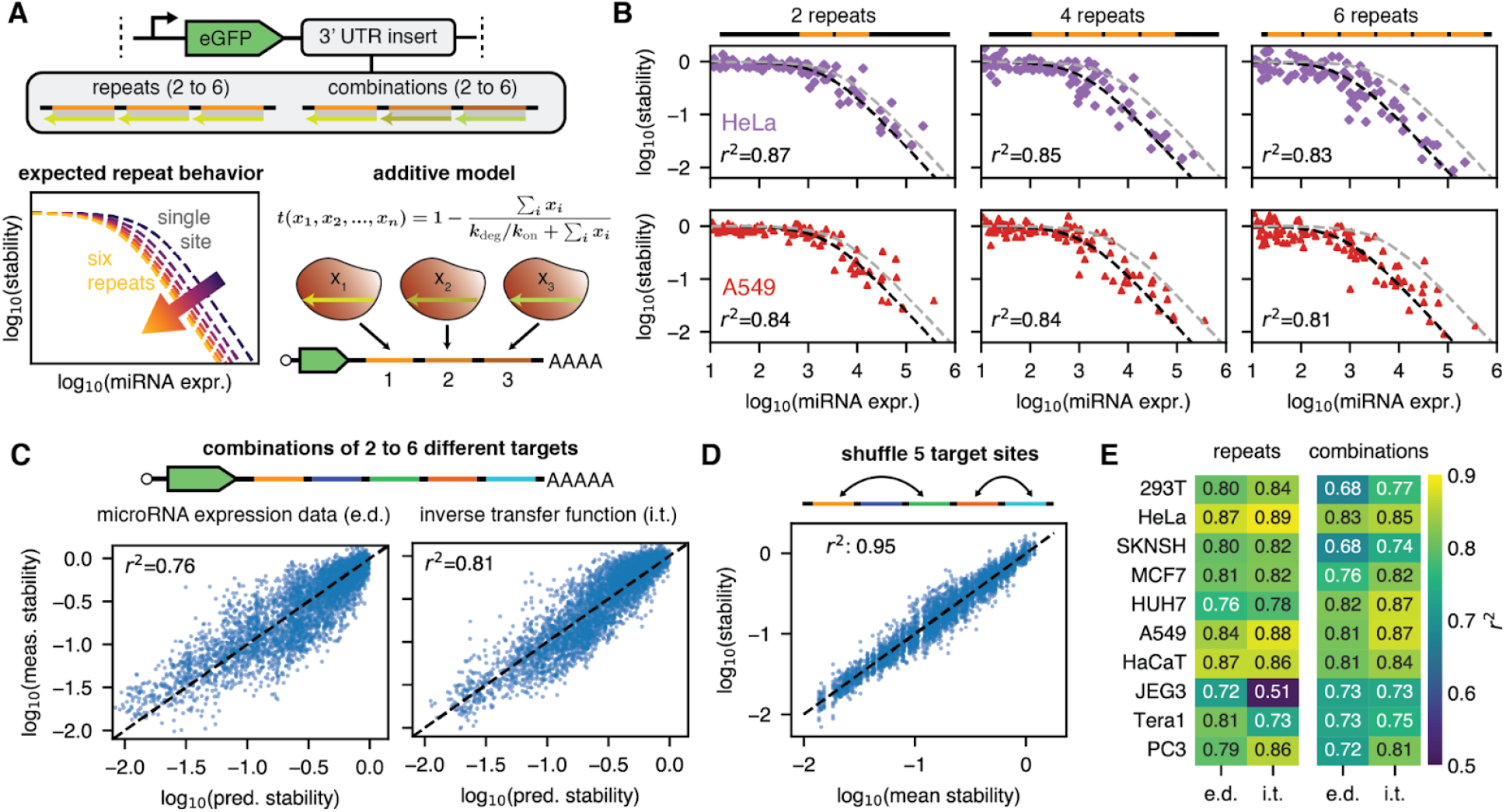
The additive model predicts the behavior of multiple target sites. (**A**) We measure the stability of repeats and combinations of target sites. The additive model predicts that the concentration of individual target sites should be summed before the transfer function is applied. (**B**) Measurements and predictions using the additive model (dashed black line) for 2 to 6 target site repeats in two cell lines. We use scaled, merged and crosstalk-filtered microRNA expression data. The gray dashed line shows the prediction for a single target site. (**C**) Measurements and predictions using the additive model for combinations of 2 to 6 target sites across all our cell lines. We use either scaled, merged and crosstalk-filtered microRNA expression data (left) or expression data inferred from the stability data for individual target sites via inversion of the transfer function (right). (**D**) Mean and individual stability data for combinations of 5 target sites with shuffled ordering on the 3’UTR. (**E**) Comparison of model fit Pearson r^2^ values for repeats and combinations of miRNA target sites for actual miRNA expression data (e.d.) or expression data derived from inverting the transfer function for individual sites (i.t).

To investigate the behavior of multiple target sites for different miRNAs, we tested 500 combinations of 2 to 6 different miRNA targets chosen to best distinguish different models (Methods). We see an overall r^2^ of 0.76 (**Fig. 2C**, left panel) for the correlation between the additive model and measured data, confirming that the additive model also works well for multiple different miRNAs. A prediction based simply on the strongest target site systematically overpredicts the stability (**Fig. S23**). Although stability prediction works well, there are still outliers. To distinguish between effects not considered in the additive model and inaccuracies in our expression data, we used the stability data for single target sites to estimate the true miRNA expression levels (inverted transfer function model, see methods). This approach reduces some of the deviation from ideal behavior in the prediction (**Fig. 2C**, right panel).

As a further test of the additive model, we chose 30 different sets of 5 targets and generated 15 randomly shuffled versions of their position in the 3’UTR. If secondary structure, cooperative effects, proximity to the coding region, or other factors matter, shuffling of target site position should change the measured stability. If the additive model is correct, the position should make no difference. We compare the mean stability of the 15 sites with the individual measured stabilities (**Fig. 2D**). With an r^2^ of 0.95, we can conclude that the position along the UTR is not relevant. Overall, the updated model prediction accuracy from expression data comes close to matching and sometimes exceeds that of the inverted transfer function model, which requires difficult-to-collect activity data, across all cell lines for both repeats and combinations (**Fig. 2E**).

Although the additive model works well for most miRNA combinations, there are notable outliers, especially in the case of repeated target sites for the same miRNA. First, some miRNAs show a pattern where an even number of repeats has less activity than an odd number (**Fig. S24A-B**). This is likely due to self-complementarity of the target, which causes strong target-target interactions and prevents miRNA binding via secondary structure formation (**Fig. S24C-D**). Unbiased discovery of all occluded targets via ΔΔG calculations is difficult due to the low accuracy of secondary structure prediction for mRNAs in cells (**Fig. S24E-F**) (*57*). Second, some miRNA target sites lead to a nearly monotonic increase in stability with increasing target numbers (**Fig. S25A**). This effect occurs more strongly in cell lines where the cognate miRNA is not expressed, indicating that this is not an effect of non-canonical miRNA regulation (**Fig. S25B-C**). We suspect RBPs as a potential cause, although identification of concrete candidates is difficult using current prediction methods.

### MicroRNA target sites in the 5’UTR follow the same transfer function but have unexpected outliers and translational effects

Although far less common for natural miRNA regulation, miRNA target sites in the 5’UTR of genes can be a viable engineering strategy and have potential advantages over target sites in the 3’UTR. Target sites in the 5’UTR have been reported to have a stronger overall effect on protein expression than those in the 3’UTR (*58*), to work synergistically with those in the 3’UTR (*16*), and to be less sensitive to pseudouridine modification (*59*). Thus, we were curious whether our model would also predict the effect of target sites in the 5’UTR.

We introduced our miRNA target library into the 5’UTR of our expression construct and measured stabilities in HEK293T and HeLa (**Fig. 3A**). The overall distribution of stabilities is similar and highly correlated (r=0.84 for HEK293T, r=0.81 for HeLa) between the 3’ and 5’UTR, although there are more unstable sequences for target sites in the 5’UTR (**Fig. 3B, Fig. S26A-C**). Looking at the performance of our improved model using the same parameters as for the 3’UTR across different numbers of target site repeats, we find that it performs well for one to three repeats (up to r^2^∼0.8 for two target sites). The reduction of stability due to miRNA targets is slightly smaller than in the 3’UTR, possibly due to occlusion of potential target sites by ribosomes (**Fig. 3C**). For an increasing number of target sites, we find that while active miRNA sites are still generally predicted well, there is an unexpected loss of stability for some target sites with low corresponding miRNA expression. This effect is most pronounced for designs containing six target repeats. Because this stability reduction predominantly happens for target sites whose associated miRNAs are not expressed, we can be nearly certain that this loss of stability is unrelated to microRNA regulation. Notably, the presence of miR-203a-3p target sites is strongly associated with reduced stability, especially when they are close to the 5’ end of the mRNA (**Fig. S26D-F**).

**Fig. 3.**
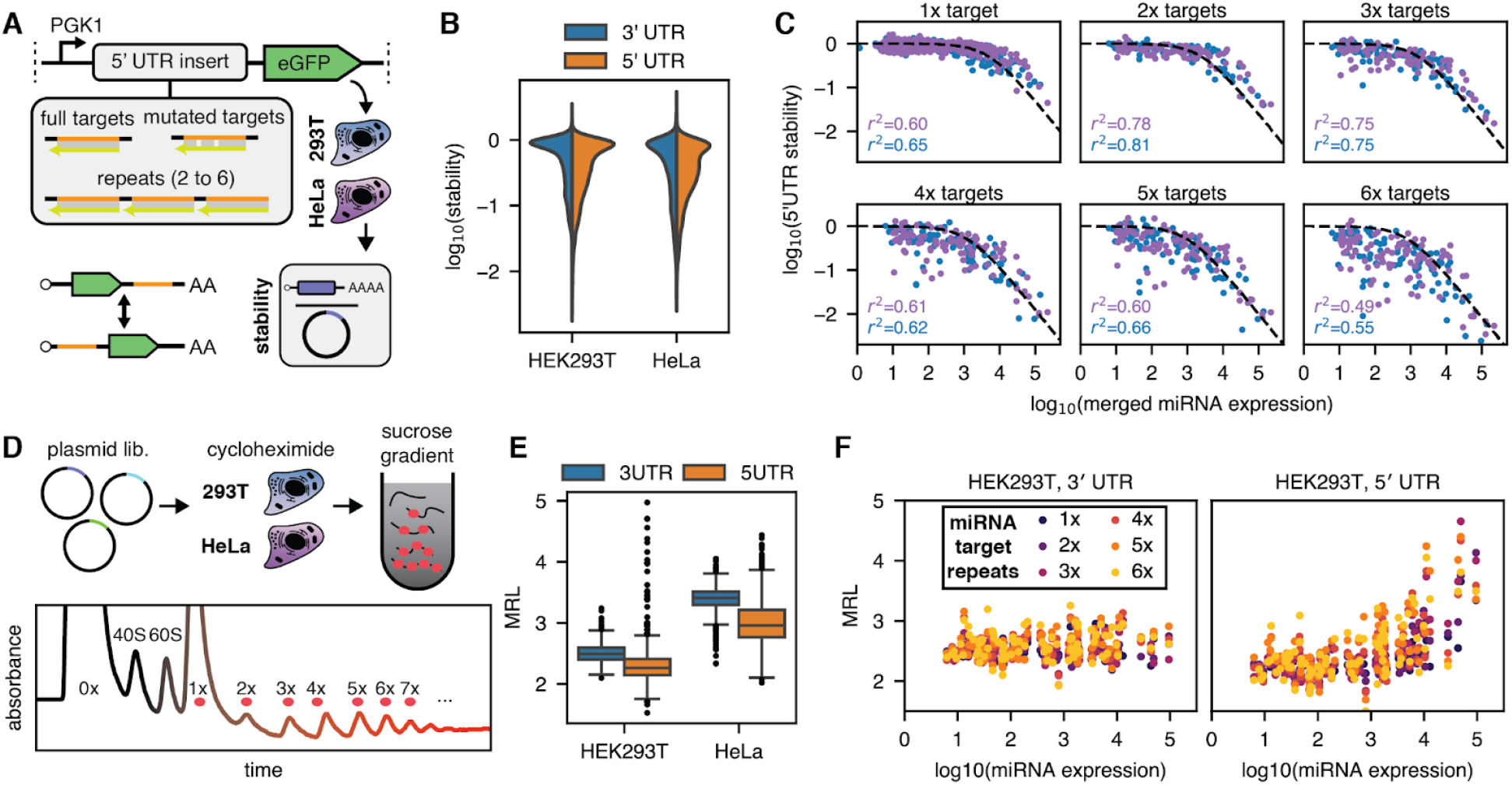
MicroRNA target sites in the 5’ UTR follow the same transfer function. (**A**) We measure the same library as for the 3’UTR inserted into the 5’ UTR in HEK293T and HeLa cells using the ratiometric stability assay. (**B**) Distribution of stabilities in the 3’ and 5’ UTR. (**C**) Stability versus microRNA expression for different numbers of microRNA target repeats in the 5’ UTR. (**D**) We use polysome profiling to determine the translational effects of microRNA target sites in the 3’UTR and 5’UTR. (**E**) Distribution of the mean ribosome load (MRL) by cell line and UTR. (**F**) Impact of different numbers of repeats of microRNA target sites versus the expression of their associated microRNA.

Beyond modulating stability, 5’UTR target sites could also impact translation efficiency. We performed polysome profiling followed by sequencing (*60, 61*) on our 3’ and 5’UTR libraries in HEK293T and HeLa, which yields a distribution across different ribosome counts for each construct in our library (**Fig 3D**), from which we calculated Mean Ribosome Load (MRL) as a measure of translation efficiency to assess potential effects of miRNA target sites, **Fig. S27A-G**). The distribution of MRLs is relatively narrow but broader for the 5’ UTR library (**Fig. 3E**). For miRNA target sites in the 3’UTR, the MRL distribution is independent of miRNA expression (**Fig. 3F, Fig. S27H-I**), underscoring our earlier flow cytometry data indicating that full target sites in the 3’UTR do not affect translation (cf. **Fig S5**). For sites in the 5’ UTR, we observe an increase in MRL both with the number of target sites and the miRNA expression. We note that the direction of the effect is unexpected and might be due to a selection effect in which actively translated mRNAs are less likely to be degraded due to ribosome-RISC interactions.

In summary, although the effect of miRNA target sites in the 5’UTRs are well predicted by the same transfer function used for 3’UTR targets, there is more variability due to stability-modifying effects unrelated to miRNA regulation and the presence of translational effects. More generally, for mRNA therapy applications, the 5’UTR sequence is strongly constrained by the need for optimization for strong expression (*61*). It is therefore usually less attractive for engineering purposes despite potential advantages and we will focus on 3’UTR design in the following.

### Model-based design of 3’UTRs with tailored stability profiles

We used our model to design 3’UTRs with defined stability patterns based purely on miRNA expression data (**Fig. 4A**). The chosen stability patterns represent tasks of varying difficulty given the repressive nature of miRNA regulation. We test “binary” designs aiming for low expression (stability 0) in one or more target cell lines and high expression (stability 1) everywhere else, binary designs resulting in high expression in the target cell lines (stability 1) and low expression in all others (stability 0), and graduated expression patterns, where the goal is to achieve randomly chosen intermediate stabilities between 0 and 1 for each cell line (**Fig. 4B**). The baseline model was used to generate the designs. This reflects the real-world challenge of creating designs for tissues and cell types for which miRNA expression data is available but for which we have no reporter activity measurements. We used an evolutionary algorithm to select four to six target sites based on the weighted mean squared error (mse) between the target and the predicted pattern. We created designs for either a subset of six or all cell lines based on the notion that the design should be easier for a smaller number of target cell lines or for all 10 cell lines. In total, we generated 1,782 designs for six cell lines and 1,986 designs for all cell lines, respectively (**Fig. S28**).

**Fig. 4.**
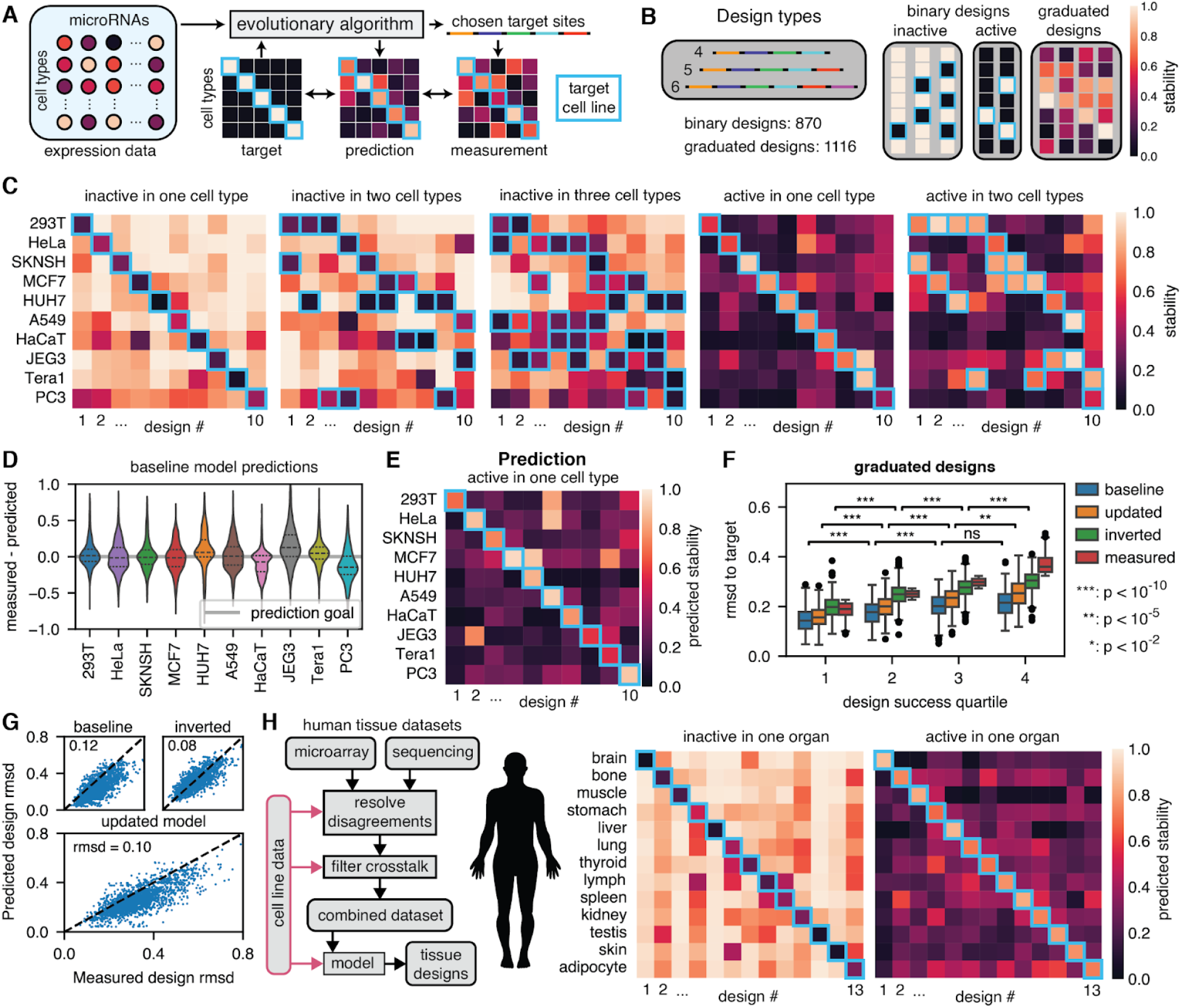
Model-based design of tailored stability profiles from expression data. (**A**) An evolutionary algorithm uses expression data and our model to generate and predict 3’UTR designs. Designs are chosen based on a weighted root mean square deviation (RMSD) between predicted and target stability patterns. (**B**) Different stability patterns across cell lines are designed using 4,5 or 6 miRNA target sites. Designs either use binary expression patterns or graduated patterns. (**C**) Best-performing designs for either high or low stability in one or a few cell lines and the opposite in the others. The blue boxes show the cell lines in which high/low stability is desired. (**D**) Deviations of the measurement for binary designs from the predictions of the baseline model used for design. (**E**) Prediction by the baseline model for the best-performing designs with high stability in a single cell line. (**F**) RMSD of model predictions from target stability values for graduated designs across different quartiles of design success (measured RMSD to the target pattern). p-values were calculated using a two-sided Mann-Whitney U test. (**G**) Predicted and measured RMSD values from the target stabilities for the different models. (**H**) Two human tissue datasets collected using different methods were merged. We used our cell line data to help resolve disagreements and filter crosstalk. We used our model to create designs with up to six target sites that are predicted to be either active or inactive in a single organ.

For the binary designs, we experimentally tested four 3’UTRs per design target type using 4, 5, or 6 miRNA target sites (**Fig. S29-31**). The best-performing designs generally come close to achieving the desired pattern (**Fig. 4C**). Easier design tasks lead to better results: Most designs that suppress stability in one or two cell lines have relatively little off-target knockdown, but constraining expression to one or two cell lines or suppressing activities in three cell lines is much more challenging. Still, we observe deviations from the target pattern even for the best designs. We next asked whether these deviations are predicted by our models or whether they are purely random. First, we note that measurements and predictions generally agree (**Fig. S29-31**). Approximately 70% of predictions have a distance of 0.2 or less to the measured value (**Fig. 4D**).

The difficulty of the design types is anticipated by the target-prediction root mean square deviation (rmsd) (**Fig. S32A**). Designs with five or more target sites lead to significantly better predicted and measured results for binary designs constraining expression to a single cell line but not for binary designs inactivating expression (**Fig. S32B**). **Fig. 4E** shows an example prediction for restraining activity to a single cell type. Many of the most prominent deviations from the desired pattern, e.g., off-target activity in JEG3 for the second design, are predicted by the model. Two other model types further increase the prediction accuracy (updated and inverted transfer function, **Fig. S32C-E**). These other two models represent how well we can predict stability given knowledge of total miRNA concentrations and background stability (updated model) and with access to a prior high-throughput assay of miRNA activity in the same system (inverted transfer function model). Notably, all models predict performance not just for well-performing designs (**Fig. S33**) but also for poorly performing ones (**Fig. S32A, S34**).

We experimentally tested 1,116 graduated designs targeting all cell lines, achieving varying levels of success in generating the desired patterns (**Fig. S35**). Comparing across quartiles of design success, we find that for the first three quartiles, the predicted rmsd values track the measured rmsd values for all models. For the fourth quartile, the baseline model loses accuracy (**Fig. 4F**). Predictions and measurements in individual cell lines have an r^2^ of 0.6 for the updated model (**Fig. S36A**) and 75% of predictions deviate less than 0.2 from the measured value (**Fig. S36B**). 84% of designs have a mean absolute prediction error smaller than 0.2 across cell lines (**Fig. S36C**). As for the binary model, we can therefore anticipate the feasibility of achieving a certain stability pattern and the success or failure of an individual design is often predicted in advance (**Fig. S36D**).

Thus, all models predict reasonably well whether a certain design goal will be fulfilled, although the two expression data-based models overpredict design success overall (**Fig. 4G**). Large unexpected failures also occur (**Fig. S37**). Strong target occlusion due to target-target interactions is again one potential cause. In the most extreme cases, sites from the 5p and 3p arm of related miRNAs are present on the same 3’UTR, leading to a systematic underprediction of the stability. Such designs can easily be excluded by prohibiting strong secondary structure or target-target interactions. A few designs exhibit overall very high or very low stabilities for unknown reasons.

### Targeting mRNA to human tissues

Next, we asked what sort of mRNA expression patterns are possible in real tissues (**Fig. 4H**). We chose a microarray (*62*) and an improved sequencing dataset (*63*) whose collection method matches our cell line expression data (**Fig. S38**). The two datasets show little correlation before consistent outliers are addressed (**Fig. S39**). We merged the two datasets similarly as for our cell line data (**Fig. S40**). We then used the merged dataset to generate designs with between 1 and 8 target sites that either exclude or constrict expression to a single tissue (**Fig. 4H, S41A-B, Table S10**). While exclusion in some tissues with highly specific miRNAs, e.g. miR-122-5p in liver, can often be achieved by a single target site that could be chosen without a model, constraining expression to a single tissue requires many target sites and a careful balancing of on-target activity with unavoidable off-target activity that cannot be achieved without a quantitative model (**Fig. S41C-E**). Even for exclusion of a single tissue, the choice of miRNA is often not obvious and the optimal number of target sites is rarely the four repeats that are often used in the literature.

### Distinguishing rested and terminally exhausted mouse CD8+ T cell states using microRNA regulation

MicroRNA expression across distantly-related tissues is often highly divergent. Leveraging miRNA expression to enable targeted gene therapy or cell engineering applications requires detection and prediction of differential responses in closely-related cell types. As an extreme case, we therefore tested prediction of reporter construct behavior between different states of a single cell type, the T cell. Upon recognition of tumors, T cells differentiate from a resting state to a terminally ‘exhausted’ state, in which they are reduced in their self-renewal and tumor-killing capability (*64, 65*). Differentiation into this terminally exhausted state limits effective immunotherapy responses against solid tumors. We therefore tested prediction of differences between rested and exhausted CD8+ T cell states.

We created a library of repeats and combinations of mouse miRNA target sites for miRNAs that are highly expressed in mouse immune cells (*66*) and inserted it into the 3’UTR of a murine stem cell virus (MSCV) fluorescent reporter construct (**Fig. S42A**). We first verified that this new reporter library is correctly predicted by our transfer function in HEK293T cells (**Fig. S42B-F**). We find excellent agreement across different numbers of targets (r^2^∼0.8), though for this reporter we observe saturation of miRNA-mediated degradation at very high concentrations and repeat numbers (see **Supplementary Text 1**). We next verified that the model works for mouse cells by measuring the reporter library in 3T3 cells (**Fig. S42G-H**). Using improved sequencing data, we find good agreement (r^2^∼0.6). We note that the use of improved sequencing data alone, as opposed to merged microarray and improved sequencing data, implies lower expected model accuracy (**Fig. 1D**).

To characterize T cell states, we isolated CD8+ T cells from mouse spleens, using mice with a knock-in YFP reporter for *Tcf7* (encoding TCF-1), a key marker for a resting or precursor, self-renewing T cell state (*67*). We then activated T cells by T cell receptor (TCR) stimulation with αCD3/CD28 antibodies and transduced them with the reporter library. Transduced T cells were either cultured in IL-15 without TCR stimulation for 2 days, to achieve a rested T cell state; alternatively, they were cultured in IL-2, IFNα, and TCR stimulation for 5 days, to drive T cells into an exhausted state (**Fig. 5A**). We observed a strong decrease in stemness marker TCF-1, and strong increases in the inhibitory receptors PD-1 and TIM-3 in the exhausted compared to the rested condition, confirming that we generated the desired cell states (**Fig. 5B**).

**Fig. 5.**
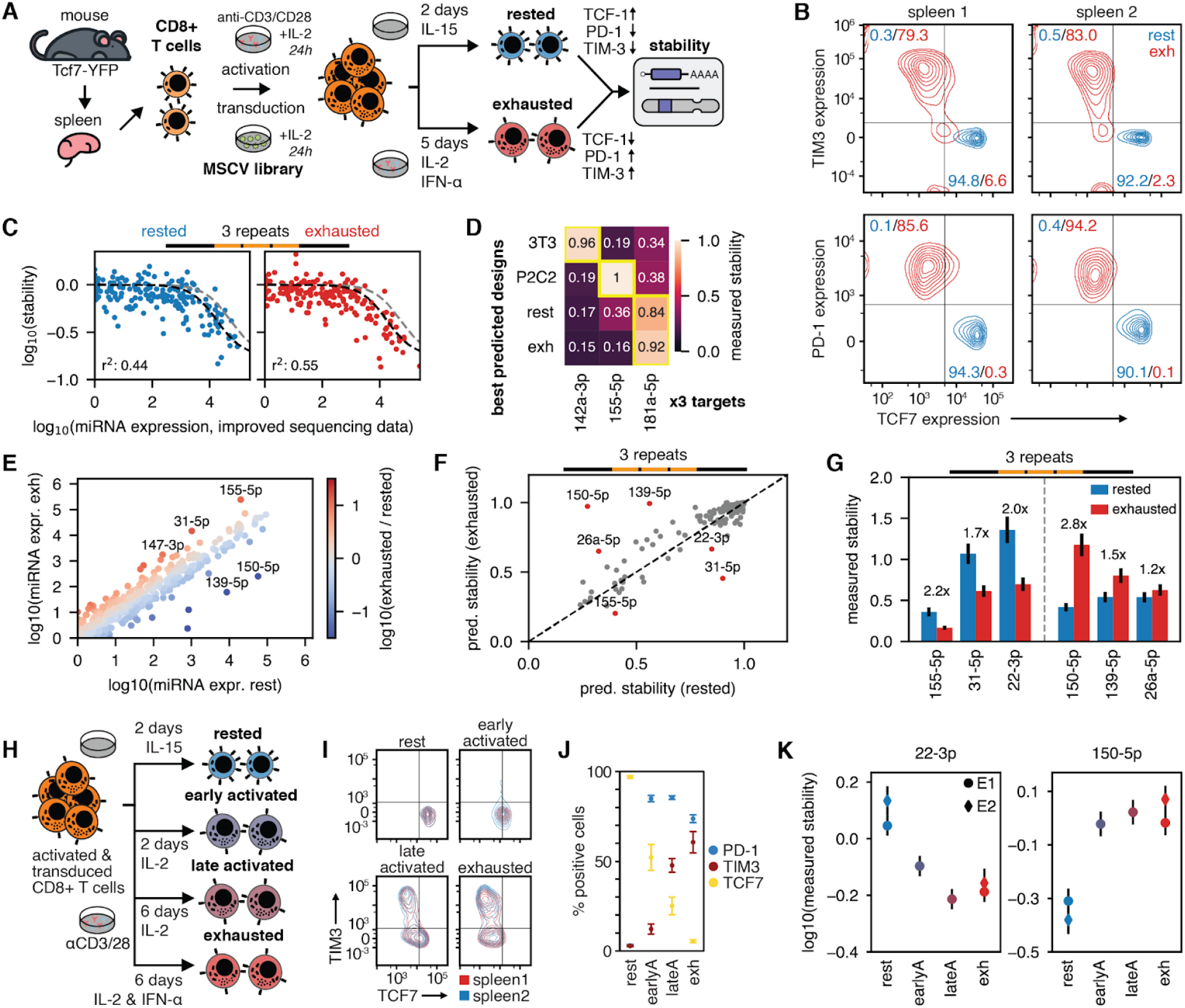
Distinguishing CD8+ T cell states using microRNA target regulation. (**A**) MSCV library transduction and T cell differentiation protocol. (**B**) Flow cytometry data for T cell state markers. (**C**) Stability versus miRNA expression for three target repeats in CD8+ T cell states. The black dashed lines show the transfer function, the gray lines the transfer function for a single target site. (**D**) Measured stabilities for three miRNA target site repeats predicted to best distinguish CD8+ T cells from 3T3 and P2C2 cells. (**E**) miRNA expression in rested and exhausted T cells. (**F**) Predicted stabilities in rested and exhausted T cells for three target repeats. (**G**) Measured stabilities of microRNA target site repeats predicted to best distinguish rested from exhausted states. (**H**) Protocol and culture conditions used to generate four levels of T cell differentiation. (**I**) Flow cytometry measurements of markers for four different T cell culture conditions. (**J**) Number of cells positive for PD-1, TIM-3, and TCF7 across the four different culture conditions. (**K**) Stability of the two microRNAs that best distinguish rested and exhausted cells in a first experiment (E1) across T cell culture conditions (E2).

We generated miRNA expression data using improved sequencing and calculated reporter stabilities in CD8+ cells by comparing mRNA abundance to integrated reporter gDNA abundance for each construct (**Fig. S43, Fig. S44A-B**). We cannot use plasmid counts as a reference because the miRNA regulation causes differential stability of viral mRNAs and therefore packing of viral constructs during production in HEK293T. This effect is much weaker than in the absence of packaging plasmid, likely because packed viral RNA is protected, and the dynamics of miRNA degradation compete with packing dynamics (**Fig. S44C**). Model prediction performance is in line with expectations for improved sequencing data (r^2^=0.44/0.56 for three target repeats in rested/exhausted CD8 T cells, **Fig. 3C**), confirming that the model works for primary mouse cells. Overall, miRNA regulation in CD8+ T cells is much weaker than in most cell lines (k_deg_/k_on_≈10^4.35^ in CD8+ T cells versus k_deg_/k_on_≈10^3.5^ in 3T3).

Compared to cell lines, in CD8+ T cells, we observe a stronger decrease in prediction performance with a larger number of target site repeats (r^2^=0.66/0.48 for one/five target sites, **Fig. S45A-D**). As with 5’UTR target sites, this variability is primarily observed for target sites whose cognate miRNAs are not expressed, making miRNA regulation an unlikely culprit. In data from mouse P2C2 cells transduced with our viral library, the increase in noise with the number of target sites is comparable to human cell lines (**Fig. S45E-F**), excluding the possibility that differences in the assay itself are responsible. The deviation from the transfer function is highly correlated between replicates and T cell states (**Fig. S46**) and can therefore be seen as representing a genuine biological effect. Indeed, three target sites that reproducibly strongly increase stability when repeated (miR-365-3p, miR-129-1-3p, miR-129-2-3p) each contain the same UUUUU(A)GGGG motif. We primarily focus on three target repeats in the following analysis to circumvent this issue.

Before addressing the question of distinguishing between T cell states, we asked whether we could reliably discern T cells from other cell types. We find that T cells, regardless of state, can be readily distinguished from 3T3 and P2C2 cell lines. Of all miRNA targets in our library, the ones predicted to perform best (margin between on- and off-target) did, in fact, best discriminate primary CD8+ from T cell lines (**Fig. 5D**).

Distinguishing T cell states is less straightforward, as miRNA expression is highly similar for the rested and exhausted T cells (**Fig. 5E**). Only a handful of miRNAs stand out: miR-155-5p and miR-31-5p are more strongly expressed in the exhausted state, miR-150-5p and miR-39-5p are more strongly expressed in the rested state. Applying our model contextualizes these miRNAs, predicting miR-155-5p as a less appealing option for discerning the exhausted T cell state due to excessive loss of stability in rested cells, and revealing miR-22-3p as a promising alternative (**Fig. 5F**). This is substantiated in stability measurements; miR-22-3p is, in fact, the top miRNA marker of T cell exhaustion, eliciting strong downregulation of the reporter in exhausted cells while maintaining high relative stability in the rested state (2.0x fold change). miR-150-5p is the top miRNA marker of T cell rest, demonstrating the reverse: strong downregulation in rested and relatively high stability in exhausted cells (2.8x fold change) (**Fig. 5G**).

Next, we investigated when miRNA changes occur as T cells differentiate from rested to exhausted states. We tested our reporter library in four culture conditions consisting of varying cytokine and exposure time combinations, each intended to elicit distinct cell states (rested, early activated, late activated, and exhausted) along T cell differentiation from naive to exhausted.

This assay evaluated the robustness of miRNA expression profiles across rested and exhausted T cell states (**Fig. 5H**). Phenotypic markers follow the expected progression along the cell state trajectory: TCF-1 is down-regulated with differentiation, PD-1 switches on early after activation, and TIM-3 increases only later (**Fig. 5I-J, Fig. S47-S48**). We find that miR-22-3p reporter stability drops from rested to late activation, roughly following TIM-3 levels (**Fig. 5K**). miR-150-5p reporter stability is low in rest and high in the other conditions, changing concomitantly with PD-1 expression during activation. Together, these results demonstrate the utility of our approach in identifying miRNA target sequences with state-specific regulation in a single cell type.

## Discussion

In summary, we find that a universal transfer function predicts the stability of miRNA-responsive UTRs from total miRNA levels as calculated by an additive model. Universality here means that the regulatory effect is a function of the miRNA concentration but is largely independent of miRNA identity and of the cellular context. The choice of miRNA expression data is key and the universal function becomes apparent only when using high-quality miRNA expression datasets. Conversely, our reporter stability data provide an easy way of benchmarking collection methods by measuring expression for our used cell lines and seeing whether the universal relationship is retained. Systematic treatment of biases increases the prediction accuracy: The expression data should be scaled to total miRNA levels and outliers removed by combining data sources. For the microRNA target sites, strong secondary structure should be eliminated and those that experience strong crosstalk excluded. Our results suggest that earlier conflicting results on the relationship between miRNA levels and activity can be resolved by careful selection and processing of the expression data. Given the ubiquitous use of miRNA regulation, this insight should be highly useful for gene therapy and cellular control circuits.

While it has long been established that higher miRNA expression tends to result in lower reporter expression levels (*15, 68*), we here show that well-processed expression data alone allows for quantitative predictions. Conversely, the typical error introduced by the use of poor expression data hampers design efforts far more than the lack of complex prediction models that take into account the biological peculiarities of particular miRNAs. In fact, the likely single largest remaining source of prediction inaccuracy in our model is the stability impact of entirely miRNA-independent effects caused by changing the transcript sequence when introducing miRNA target sites. While beyond the scope of this manuscript, quantitatively accurate models of, e.g., RBP regulation therefore seem the most promising direction in further increasing prediction accuracy. Given the enormous impact of miRNA expression data quality, residual technical bias probably also plays a large role. And while comparatively weaker, including effects such as seed target abundance or target secondary structure will further improve model accuracy, although they are hard to exactly quantify while much larger sources of error remain.

We designed hard-to-achieve patterns such as restriction to a single cell line or graduated expression. Importantly, we show that such patterns are achievable even with a limited number of target sites, resulting in compact 3’UTRs. We expect such designs to be practically important for increasing the specificity of future mRNA therapies where few other levers for achieving specificity are available. However, the simple nature of miRNA regulation, namely that it is repressive and non-cooperative, still constrains the achievable patterns. Incorporating our insights into the design of multi-gene circuits of miRNA-regulated mRNAs could address this limitation (*9*).

Going beyond cell lines, we have demonstrated that our model correctly predicts miRNAs that distinguish rested and exhausted mouse CD8+ T cell states. For designs beyond immune cells that are easily sorted based on surface markers, the heterogeneity of cell types in a single tissue could pose a significant hurdle because tissue-averaged miRNA expression profiles might not be representative of any single cell type within the tissue. Future single-cell miRNA sequencing techniques (*69*) could help overcome this limitation, though only if they avoid the biases of standard library construction workflows.

We here focus on fully complementary target sites relevant for engineering applications. While biologically relevant seed target sites require more complex models that take into account the secondary structure and miRNA sequence features (*70*), any predictive model will also require debiased expression data as an accurate proxy for concentrations, making the elimination of residual biases equally relevant for basic biology.

In the longer term, we expect that the approach introduced here, i.e., learning quantitative relationships between the levels of trans-regulatory molecules and their targets that allow generalization from one cellular context to the other, can also be applied to RNA binding proteins or transcription factors, enabling the rational design of functional cis-regulatory elements given knowledge of only transregulator expression.

## Supporting information

Design of Library 1

Stability data Library 1

Design of Library 2

Stability data Library 2

Design of Library 3

Stability data Library 3

microRNA expression data

Processed data Actinomycin D

Processed data polysome profiling

## Acknowledgements

We thank Jesse Bloom, Paul Nghiem, and Andrew Hsieh for providing cell lines. We thank the members of the Seelig lab for discussions on this manuscript.

## Author contributions

L.O. and S.C.H. collected the high-throughput stability data. N.L., S.R. and A.K. collected the miRNA expression data generated for this study. A.K. gave advice on the analysis and interpretation of the results. M.B. and H.Y.K. performed the CD8+ T cell differentiation and transduction experiments. L.O. designed the constructs and analyzed the data. L.O., S.C.H., and G.S. designed the project. L.O. and G.S. wrote the manuscript with input from the other authors.

## Data and materials availability

Files specifying library sequences, used microRNA target sites, processed stability data, and other relevant information necessary to reproduce the results are provided as **Supplementary Data**. Pre-processed microRNA expression data generated for this manuscript, as well as merged and crosstalk-filtered microarray and improved sequencing data for the used cell lines and human tissues are also provided as **Supplementary Data**. The analysis code for this manuscript is available on GitHub at loesinghaus/miRNA.design.2025. Raw sequencing data for microRNA expression will be made available on GEO. Raw sequencing and library count data are available on GEO (GSE283210).

## References

1. G. M. Allen, W. A. Lim, Rethinking cancer targeting strategies in the era of smart cell therapeutics. Nat. Rev. Cancer 22, 693–702 (2022).

2. I. I. Taskiran, K. I. Spanier, H. Dickmänken, N. Kempynck, A. Pančíková, E. C. Ekşi, G. Hulselmans, J. N. Ismail, K. Theunis, R. Vandepoel, V. Christiaens, D. Mauduit, S. Aerts, Cell-type-directed design of synthetic enhancers. Nature 626, 212–220 (2024).

3. J. Zhang, K. Salaita, Smart Nucleic Acids as Future Therapeutics. Trends Biotechnol. 39, 1289–1307 (2021).

4. A. P. Teixeira, M. Fussenegger, Synthetic macromolecular switches for precision control of therapeutic cell functions. Nat. Rev. Bioeng., 1005–1022 (2024).

5. B. D. Brown, B. Gentner, A. Cantore, S. Colleoni, M. Amendola, A. Zingale, A. Baccarini, G. Lazzari, C. Galli, L. Naldini, Endogenous microRNA can be broadly exploited to regulate transgene expression according to tissue, lineage and differentiation state. Nat. Biotechnol. 25, 1457–1467 (2007).

6. B. Dhungel, C. Ramlogan-Steel, J. Steel, MicroRNA-Regulated Gene Delivery Systems for Research and Therapeutic Purposes. Molecules 23, 1500 (2018).

7. L. F. R. Gebert, I. J. MacRae, Regulation of microRNA function in animals. Nat. Rev. Mol. Cell Biol. 20, 21–37 (2019).

8. Z. Xie, L. Wroblewska, L. Prochazka, R. Weiss, Y. Benenson, Multi-Input RNAi-Based Logic Circuit for Identification of Specific Cancer Cells. Science 333, 1307–1311 (2011).

9. P. Mohammadi, N. Beerenwinkel, Y. Benenson, Automated Design of Synthetic Cell Classifier Circuits Using a Two-Step Optimization Strategy. Cell Syst. 4, 207–218.e14 (2017).

10. B. Angelici, L. Shen, J. Schreiber, A. Abraham, Y. Benenson, An AAV gene therapy computes over multiple cellular inputs to enable precise targeting of multifocal hepatocellular carcinoma in mice. Sci. Transl. Med. 13, eabh4456 (2021).

11. L. Wang, W. Xu, S. Zhang, G. C. Gundberg, C. R. Zheng, Z. Wan, K. Mustafina, F. Caliendo, H. Sandt, R. Kamm, R. Weiss, Sensing and guiding cell-state transitions by using genetically encoded endoribonuclease-mediated microRNA sensors. Nat. Biomed. Eng., 1730–1743 (2024).

12. R. Jain, J. P. Frederick, E. Y. Huang, K. E. Burke, D. M. Mauger, E. A. Andrianova, S. J. Farlow, S. Siddiqui, J. Pimentel, K. Cheung-Ong, K. M. McKinney, C. Köhrer, M. J. Moore, T. Chakraborty, MicroRNAs Enable mRNA Therapeutics to Selectively Program Cancer Cells to Self-Destruct. Nucleic Acid Ther. 28, 285–296 (2018).

13. S. E. Sinnett, E. Boyle, C. Lyons, S. J. Gray, Engineered microRNA-based regulatory element permits safe high-dose mini MECP2 gene therapy in Rett mice. Brain 144, 3005–3019 (2021).

14. B. Brook, V. Duval, S. Barman, L. Speciner, C. Sweitzer, A. Khanmohammed, M. Menon, K. Foster, P. Ghosh, K. Abedi, J. Koster, E. Nanishi, L. R. Baden, O. Levy, T. VanCott, R. Micol, D. J. Dowling, Adjuvantation of a SARS-CoV-2 mRNA vaccine with controlled tissue-specific expression of an mRNA encoding IL-12p70. Sci. Transl. Med. 16, eadm8451 (2024).

15. G. Mullokandov, A. Baccarini, A. Ruzo, A. D. Jayaprakash, N. Tung, B. Israelow, M. J. Evans, R. Sachidanandam, B. D. Brown, High-throughput assessment of microRNA activity and function using microRNA sensor and decoy libraries. Nat. Methods 9, 840–846 (2012).

16. J. J. Gam, J. Babb, R. Weiss, A mixed antagonistic/synergistic miRNA repression model enables accurate predictions of multi-input miRNA sensor activity. Nat. Commun. 9, 2430 (2018).

17. O. Flores, E. M. Kennedy, R. L. Skalsky, B. R. Cullen, Differential RISC association of endogenous human microRNAs predicts their inhibitory potential. Nucleic Acids Res. 42, 4629–4639 (2014).

18. A. Kozomara, S. Hunt, M. Ninova, S. Griffiths-Jones, M. Ronshaugen, Target Repression Induced by Endogenous microRNAs: Large Differences, Small Effects. PLoS ONE 9, e104286 (2014).

19. I. Vainberg Slutskin, S. Weingarten-Gabbay, R. Nir, A. Weinberger, E. Segal, Unraveling the determinants of microRNA mediated regulation using a massively parallel reporter assay. Nat. Commun. 9, 529 (2018).

20. P. Y. Wang, D. P. Bartel, The guide-RNA sequence dictates the slicing kinetics and conformational dynamics of the Argonaute silencing complex. Mol. Cell 84, 2918–2934.e11 (2024).

21. D. M. Garcia, D. Baek, C. Shin, G. W. Bell, A. Grimson, D. P. Bartel, Weak seed-pairing stability and high target-site abundance decrease the proficiency of lsy-6 and other microRNAs. Nat. Struct. Mol. Biol. 18, 1139–1146 (2011).

22. R. Denzler, S. E. McGeary, A. C. Title, V. Agarwal, D. P. Bartel, M. Stoffel, Impact of MicroRNA Levels, Target-Site Complementarity, and Cooperativity on Competing Endogenous RNA-Regulated Gene Expression. Mol. Cell 64, 565–579 (2016).

23. M. Jie, T. Feng, W. Huang, M. Zhang, Y. Feng, H. Jiang, Z. Wen, Subcellular Localization of miRNAs and Implications in Cellular Homeostasis. Genes 12, 856 (2021).

24. V. Wagner, E. Meese, A. Keller, The intricacies of isomiRs: from classification to clinical relevance. Trends Genet. 40, 784–796 (2024).

25. M. Kertesz, N. Iovino, U. Unnerstall, U. Gaul, E. Segal, The role of site accessibility in microRNA target recognition. Nat. Genet. 39, 1278–1284 (2007).

26. S. Ruijtenberg, S. Sonneveld, T. J. Cui, I. Logister, D. De Steenwinkel, Y. Xiao, I. J. MacRae, C. Joo, M. E. Tanenbaum, mRNA structural dynamics shape Argonaute-target interactions. Nat. Struct. Mol. Biol. 27, 790–801 (2020).

27. V. Agarwal, G. W. Bell, J.-W. Nam, D. P. Bartel, Predicting effective microRNA target sites in mammalian mRNAs. eLife 4, e05005 (2015).

28. J. A. Broderick, W. E. Salomon, S. P. Ryder, N. Aronin, P. D. Zamore, Argonaute protein identity and pairing geometry determine cooperativity in mammalian RNA silencing. RNA 17, 1858–1869 (2011).

29. A. D. Jayaprakash, O. Jabado, B. D. Brown, R. Sachidanandam, Identification and remediation of biases in the activity of RNA ligases in small-RNA deep sequencing. Nucleic Acids Res. 39, e141–e141 (2011).

30. P. Mestdagh, N. Hartmann, L. Baeriswyl, D. Andreasen, N. Bernard, C. Chen, D. Cheo, P. D’Andrade, M. DeMayo, L. Dennis, S. Derveaux, Y. Feng, S. Fulmer-Smentek, B. Gerstmayer, J. Gouffon, C. Grimley, E. Lader, K. Y. Lee, S. Luo, P. Mouritzen, A. Narayanan, S. Patel, S. Peiffer, S. Rüberg, G. Schroth, D. Schuster, J. M. Shaffer, E. J. Shelton, S. Silveria, U. Ulmanella, V. Veeramachaneni, F. Staedtler, T. Peters, T. Guettouche, L. Wong, J. Vandesompele, Evaluation of quantitative miRNA expression platforms in the microRNA quality control (miRQC) study. Nat. Methods 11, 809–815 (2014).

31. M. D. Giraldez, R. M. Spengler, A. Etheridge, P. M. Godoy, A. J. Barczak, S. Srinivasan, P. L. De Hoff, K. Tanriverdi, A. Courtright, S. Lu, J. Khoory, R. Rubio, D. Baxter, T. A. P. Driedonks, H. P. J. Buermans, E. N. M. Nolte-’t Hoen, H. Jiang, K. Wang, I. Ghiran, Y. E. Wang, K. Van Keuren-Jensen, J. E. Freedman, P. G. Woodruff, L. C. Laurent, D. J. Erle, D. J. Galas, M. Tewari, Comprehensive multi-center assessment of small RNA-seq methods for quantitative miRNA profiling. Nat. Biotechnol. 36, 746–757 (2018).

32. H. Kim, J. Kim, K. Kim, H. Chang, K. You, V. N. Kim, Bias-minimized quantification of microRNA reveals widespread alternative processing and 3′ end modification. Nucleic Acids Res. 47, 2630–2640 (2019).

33. A. Kozomara, M. Birgaoanu, S. Griffiths-Jones, miRBase: from microRNA sequences to function. Nucleic Acids Res. 47, D155–D162 (2019).

34. B. Fromm, D. Domanska, E. Høye, V. Ovchinnikov, W. Kang, E. Aparicio-Puerta, M. Johansen, K. Flatmark, A. Mathelier, E. Hovig, M. Hackenberg, M. R. Friedländer, K. J. Peterson, MirGeneDB 2.0: the metazoan microRNA complement. Nucleic Acids Res. 48, D132–D141 (2020).

35. D. Griesemer, J. R. Xue, S. K. Reilly, J. C. Ulirsch, K. Kukreja, J. R. Davis, M. Kanai, D. K. Yang, J. C. Butts, M. H. Guney, J. Luban, S. B. Montgomery, H. K. Finucane, C. D. Novina, R. Tewhey, P. C. Sabeti, Genome-wide functional screen of 3′UTR variants uncovers causal variants for human disease and evolution. Cell 184, 5247–5260.e19 (2021).

36. H. Guo, N. T. Ingolia, J. S. Weissman, D. P. Bartel, Mammalian microRNAs predominantly act to decrease target mRNA levels. Nature 466, 835–840 (2010).

37. J. Alles, T. Fehlmann, U. Fischer, C. Backes, V. Galata, M. Minet, M. Hart, M. Abu-Halima, F. A. Grässer, H.-P. Lenhof, A. Keller, E. Meese, An estimate of the total number of true human miRNAs. Nucleic Acids Res. 47, 3353–3364 (2019).

38. J. Martinez, T. Tuschl, RISC is a 5′ phosphomonoester-producing RNA endonuclease. Genes Dev. 18, 975–980 (2004).

39. C. Aguilar, S. Costa, C. Maudet, R. P. Vivek-Ananth, S. Zaldívar-López, J. J. Garrido, A. Samal, M. Mano, A. Eulalio, Reprogramming of microRNA expression via E2F1 downregulation promotes Salmonella infection both in infected and bystander cells. Nat. Commun. 12, 3392 (2021).

40. B. Panwar, G. S. Omenn, Y. Guan, miRmine: a database of human miRNA expression profiles. Bioinformatics 33, 1554–1560 (2017).

41. Y. Chen, S. Zhu, Y. Pei, J. Hu, Z. Hu, X. Liu, X. Wang, M. Gu, S. Hu, X. Liu, Differential microRNA Expression in Newcastle Disease Virus-Infected HeLa Cells and Its Role in Regulating Virus Replication. Front. Oncol. 11, 616809 (2021).

42. G. Grasso, T. Higuchi, V. Mac, J. Barbier, M. Helsmoortel, C. Lorenzi, G. Sanchez, M. Bello, W. Ritchie, S. Sakamoto, R. Kiernan, NF90 modulates processing of a subset of human pri-miRNAs. Nucleic Acids Res. 48, 6874–6888 (2020).

43. L. Lorenzi, H.-S. Chiu, F. Avila Cobos, S. Gross, P.-J. Volders, R. Cannoodt, J. Nuytens, K. Vanderheyden, J. Anckaert, S. Lefever, A. P. Tay, E. J. De Bony, W. Trypsteen, F. Gysens, M. Vromman, T. Goovaerts, T. B. Hansen, S. Kuersten, N. Nijs, T. Taghon, K. Vermaelen, K. R. Bracke, Y. Saeys, T. De Meyer, N. P. Deshpande, G. Anande, T.-W. Chen, M. R. Wilkins, A. Unnikrishnan, K. De Preter, J. Kjems, J. Koster, G. P. Schroth, J. Vandesompele, P. Sumazin, P. Mestdagh, The RNA Atlas expands the catalog of human non-coding RNAs. Nat. Biotechnol. 39, 1453–1465 (2021).

44. J. Alles, T. Fehlmann, U. Fischer, C. Backes, V. Galata, M. Minet, M. Hart, M. Abu-Halima, F. A. Grässer, H.-P. Lenhof, A. Keller, E. Meese, An estimate of the total number of true human miRNAs. Nucleic Acids Res. 47, 3353–3364 (2019).

45. H. Liu, P. D’Andrade, S. Fulmer-Smentek, P. Lorenzi, K. W. Kohn, J. N. Weinstein, Y. Pommier, W. C. Reinhold, mRNA and microRNA Expression Profiles of the NCI-60 Integrated with Drug Activities. Mol. Cancer Ther. 9, 1080–1091 (2010).

46. C. Camps, H. K. Saini, D. R. Mole, H. Choudhry, M. Reczko, J. Guerra-Assunção, Y.-M. Tian, F. M. Buffa, A. L. Harris, A. G. Hatzigeorgiou, A. J. Enright, J. Ragoussis, Integrated analysis of microRNA and mRNA expression and association with HIF binding reveals the complexity of microRNA expression regulation under hypoxia. Mol. Cancer 13, 28 (2014).

47. X. Qin, S. Yu, L. Zhou, M. Shi, Y. Hu, X. Xu, B. Shen, S. Liu, D. Yan, J. Feng, Cisplatin-resistant lung cancer cell–derived exosomes increase cisplatin resistance of recipient cells in exosomal miR-100–5p-dependent manner. Int. J. Nanomedicine Volume 12, 3721–3733 (2017).

48. S. K. Patnaik, J. Dahlgaard, W. Mazin, E. Kannisto, T. Jensen, S. Knudsen, S. Yendamuri, Expression of microRNAs in the NCI-60 cancer cell-lines. PloS One 7, e49918 (2012).

49. D. Luo, J. M. Wilson, N. Harvel, J. Liu, L. Pei, S. Huang, L. Hawthorn, H. Shi, A systematic evaluation of miRNA:mRNA interactions involved in the migration and invasion of breast cancer cells. J. Transl. Med. 11, 57 (2013).

50. R. T. Fuchs, Z. Sun, F. Zhuang, G. B. Robb, Bias in Ligation-Based Small RNA Sequencing Library Construction Is Determined by Adaptor and RNA Structure. PLOS ONE 10, e0126049 (2015).

51. P. M. Godoy, A. J. Barczak, P. DeHoff, S. Srinivasan, A. Etheridge, D. Galas, S. Das, D. J. Erle, L. C. Laurent, Comparison of Reproducibility, Accuracy, Sensitivity, and Specificity of miRNA Quantification Platforms. Cell Rep. 29, 4212–4222.e5 (2019).

52. D. P. Bartel, MicroRNAs: Target Recognition and Regulatory Functions. Cell 136, 215–233 (2009).

53. S. M. Elbashir, Functional anatomy of siRNAs for mediating efficient RNAi in Drosophila melanogaster embryo lysate. EMBO J. 20, 6877–6888 (2001).

54. L. M. Wee, C. F. Flores-Jasso, W. E. Salomon, P. D. Zamore, Argonaute Divides Its RNA Guide into Domains with Distinct Functions and RNA-Binding Properties. Cell 151, 1055–1067 (2012).

55. A. Grimson, K. K.-H. Farh, W. K. Johnston, P. Garrett-Engele, L. P. Lim, D. P. Bartel, MicroRNA Targeting Specificity in Mammals: Determinants beyond Seed Pairing. Mol. Cell 27, 91–105 (2007).

56. R. J. Bloom, S. M. Winkler, C. D. Smolke, A quantitative framework for the forward design of synthetic miRNA circuits. Nat. Methods 11, 1147–1153 (2014).

57. J. Zhang, Y. Fei, L. Sun, Q. C. Zhang, Advances and opportunities in RNA structure experimental determination and computational modeling. Nat. Methods 19, 1193–1207 (2022).

58. K. Miki, K. Endo, S. Takahashi, S. Funakoshi, I. Takei, S. Katayama, T. Toyoda, M. Kotaka, T. Takaki, M. Umeda, C. Okubo, M. Nishikawa, A. Oishi, M. Narita, I. Miyashita, K. Asano, K. Hayashi, K. Osafune, S. Yamanaka, H. Saito, Y. Yoshida, Efficient Detection and Purification of Cell Populations Using Synthetic MicroRNA Switches. Cell Stem Cell 16, 699–711 (2015).

59. J. Lockhart, J. Canfield, E. F. Mong, J. VanWye, H. Totary-Jain, Nucleotide Modification Alters MicroRNA-Dependent Silencing of MicroRNA Switches. Mol. Ther. Nucleic Acids 14, 339–350 (2019).

60. P. J. Sample, B. Wang, D. W. Reid, V. Presnyak, I. J. McFadyen, D. R. Morris, G. Seelig, Human 5’ UTR design and variant effect prediction from a massively parallel translation assay. Nat. Biotechnol. 37, 803–809 (2019).

61. S. Castillo-Hair, S. Fedak, B. Wang, J. Linder, K. Havens, M. Certo, G. Seelig, Optimizing 5’UTRs for mRNA-delivered gene editing using deep learning. Nat. Commun. 15, 5284 (2024).

62. N. Ludwig, P. Leidinger, K. Becker, C. Backes, T. Fehlmann, C. Pallasch, S. Rheinheimer, B. Meder, C. Stähler, E. Meese, A. Keller, Distribution of miRNA expression across human tissues. Nucleic Acids Res. 44, 3865–3877 (2016).

63. A. Keller, L. Gröger, T. Tschernig, J. Solomon, O. Laham, N. Schaum, V. Wagner, F. Kern, G. P. Schmartz, Y. Li, A. Borcherding, C. Meier, T. Wyss-Coray, E. Meese, T. Fehlmann, N. Ludwig, miRNATissueAtlas2: an update to the human miRNA tissue atlas. Nucleic Acids Res. 50, D211–D221 (2022).

64. A. Kallies, D. Zehn, D. T. Utzschneider, Precursor exhausted T cells: key to successful immunotherapy? Nat. Rev. Immunol. 20, 128–136 (2020).

65. M. Philip, A. Schietinger, CD8+ T cell differentiation and dysfunction in cancer. Nat. Rev. Immunol. 22, 209–223 (2022).

66. S. A. Rose, A. Wroblewska, M. Dhainaut, H. Yoshida, J. M. Shaffer, A. Bektesevic, B. Ben-Zvi, A. Rhoads, E. Y. Kim, B. Yu, Y. Lavin, M. Merad, J. D. Buenrostro, B. D. Brown, A microRNA expression and regulatory element activity atlas of the mouse immune system. Nat. Immunol. 22, 914–927 (2021).

67. U. Lenk, R. Hanke, U. Kräft, K. Grade, I. Grunewald, A. Speer, Non-isotopic analysis of single strand conformation polymorphism (SSCP) in the exon 13 region of the human dystrophin gene. J. Med. Genet. 30, 951–954 (1993).

68. B. D. Brown, B. Gentner, A. Cantore, S. Colleoni, M. Amendola, A. Zingale, A. Baccarini, G. Lazzari, C. Galli, L. Naldini, Endogenous microRNA can be broadly exploited to regulate transgene expression according to tissue, lineage and differentiation state. Nat. Biotechnol. 25, 1457–1467 (2007).

69. S. M. Hücker, T. Fehlmann, C. Werno, K. Weidele, F. Lüke, A. Schlenska-Lange, C. A. Klein, A. Keller, S. Kirsch, Single-cell microRNA sequencing method comparison and application to cell lines and circulating lung tumor cells. Nat. Commun. 12, 4316 (2021).

70. S. E. McGeary, K. S. Lin, C. Y. Shi, T. M. Pham, N. Bisaria, G. M. Kelley, D. P. Bartel, The biochemical basis of microRNA targeting efficacy. Science 366, eaav1741 (2019).

71. V. Agarwal, D. R. Kelley, The genetic and biochemical determinants of mRNA degradation rates in mammals. Genome Biol. 23, 245 (2022).

72. D. Briskin, P. Y. Wang, D. P. Bartel, The biochemical basis for the cooperative action of microRNAs. Proc. Natl. Acad. Sci. 117, 17764–17774 (2020).

73. M. E. Fornace, J. Huang, C. T. Newman, N. J. Porubsky, M. B. Pierce, N. A. Pierce, NUPACK: Analysis and Design of Nucleic Acid Structures, Devices, and Systems. [Preprint] (2022). 10.26434/chemrxiv-2022-xv98l.

74. S. Rishik, P. Hirsch, F. Grandke, T. Fehlmann, A. Keller, miRNATissueAtlas 2025: an update to the uniformly processed and annotated human and mouse non-coding RNA tissue atlas. Nucleic Acids Res. 53, D129–D137 (2025).

75. H. Y. Kueh, M. A. Yui, K. K. H. Ng, S. S. Pease, J. A. Zhang, S. S. Damle, G. Freedman, S. Siu, I. D. Bernstein, M. B. Elowitz, E. V. Rothenberg, Asynchronous combinatorial action of four regulatory factors activates Bcl11b for T cell commitment. Nat. Immunol. 17, 956–965 (2016).

76. M. M. Del Real, E. V. Rothenberg, Architecture of a lymphomyeloid developmental switch controlled by PU.1, Notch and Gata3. Development 140, 1207–1219 (2013).

77. T. Fehlmann, F. Kern, O. Laham, C. Backes, J. Solomon, P. Hirsch, C. Volz, R. Müller, A. Keller, miRMaster 2.0: multi-species non-coding RNA sequencing analyses at scale. Nucleic Acids Res. 49, W397–W408 (2021).

78. K. Abadie, E. C. Clark, R. M. Valanparambil, O. Ukogu, W. Yang, R. M. Daza, K. K. H. Ng, J. Fathima, A. L. Wang, J. Lee, T. H. Nasti, A. Bhandoola, A. Nourmohammad, R. Ahmed, J. Shendure, J. Cao, H. Y. Kueh, Reversible, tunable epigenetic silencing of TCF1 generates flexibility in the T cell memory decision. Immunity 57, 271–286.e13 (2024).

79. B. Muzellec, M. Teleńczuk, V. Cabeli, M. Andreux, PyDESeq2: a python package for bulk RNA-seq differential expression analysis. Bioinformatics 39, btad547 (2023).

80. H. Jin, C. Zhang, M. Zwahlen, K. Von Feilitzen, M. Karlsson, M. Shi, M. Yuan, X. Song, X. Li, H. Yang, H. Turkez, L. Fagerberg, M. Uhlén, A. Mardinoglu, Systematic transcriptional analysis of human cell lines for gene expression landscape and tumor representation. Nat. Commun. 14, 5417 (2023).

81. T. Xu, N. Su, L. Liu, J. Zhang, H. Wang, W. Zhang, J. Gui, K. Yu, J. Li, T. D. Le, miRBaseConverter: an R/Bioconductor package for converting and retrieving miRNA name, accession, sequence and family information in different versions of miRBase. BMC Bioinformatics 19, 514 (2018).

82. C. Backes, F. Sedaghat-Hamedani, K. Frese, M. Hart, N. Ludwig, B. Meder, E. Meese, A. Keller, Bias in High-Throughput Analysis of miRNAs and Implications for Biomarker Studies. Anal. Chem. 88, 2088–2095 (2016).

