## Supplementary Information for "MicroRNA-directed control of complex mRNA stability patterns across cell types"

† These authors contributed equally.

### Materials and Methods

#### Library design

We designed and measured a first test library of 1,000 sequences, a second library of 10,021 sequences in human cell lines, and a third library of 2,000 sequences in mouse cell lines and CD8 T cells. The main text focuses on Library 2. Library 1 was measured in fewer cell lines (HEK293T, HeLa, SKNSH, MCF7) and does not yield any substantial additional insights. Not all sequences included in the libraries are explicitly evaluated in the main text.

##### Library 1 design and purpose

Each sequence has a length of 142 nt, of which 15 nt were fixed at the 5' end (CGAGCTCGCTAGCCT) and 17 nt at the 3' end (AGATCGGAAGAGCGTCG), leaving a total of 110 variable nts. The purpose of Library 1 was for us to gain sufficient information to allow for the design of Library 2. We chose 145 miRNAs annotated as high confidence in miRBase v22 with high expression in the four cell lines. We measured them in two context sequences (data_high_fc_rs848_ref and data_high_fc_rs2303225_ref) of endogenous origin with neutral stability from Griesemer *et al.* (*35*). These measurements were used to fit the transfer function constant $k_{deg}/k_{on}$ that was used to create designs with tailored stability profiles (**Fig. S4A**). Library 1 also contained two and three repeats of 41 miRNAs each, and 197 combinations of between two and five different miRNAs, which allowed for an initial verification of the additive model. It also included control sequences without miRNA target sites (*71*), which were used to generate further control sequences for Library 2. Other parts of Library 1 were not used in this manuscript. Sublibraries are listed in **Table S4** and sequences are provided in **Data S1**.

##### Library 2 design

Each sequence has a length of 200 nt, of which 18 nt were fixed at the 5' end (ACGACGCTCTTCCGATCT; part of the TruSeq Read 1 sequence) and 18 nt (CTCTGGATTTGCAACCGA) at the 3' end, leaving a total of 164 variable nts. Detailed library design, composition, and miRNA selection for Library 2 are described below (see **Table S5** for a list of sublibraries and **Data S3** for library sequences). Unless otherwise mentioned, we used the microarray data by Alles et al. (*37*) when expression data was required, e.g., to choose specific miRNA target sites to test.

#### Target site insertion

The vast majority of sequences were embedded into a single context sequence generated from a combination of two control sequences (high_fc_rs848_ref and high_fc_rs7539036_ref) with high stability and low variability in stability across cell lines from Library 1. Parts of high_fc_rs7539036_ref were added to the 5' and 3' end of high_fc_rs848_ref to generate a context sequence with a total length of 164 nts. MicroRNA targets are inserted into this 'main context' for all designs in Library 2 unless otherwise mentioned. MicroRNA target sites were inserted into context sequences as follows: Single target sites were inserted approximately in the middle of the variable context sequence. Multiple target sites were inserted at a distance of 6 nt to test potential cooperativity effects (*72*). Start codons or poly(A) signals that were created by the insertion of the miRNA target were mutated by replacing the first nucleotide that is not part of the microRNA target site by a C.

**Controls**

In addition to the main context, we took all control sequences (with a length of 101 nt) from Library 1 that were measured as stable in all cell lines (mean stability of at least 1) and ranked them by the variance of their stability across the measured cell lines. We appended the least variable sequences and then trimmed them from the 3' end to get a total length compatible with Library 2 (164 nt). We retained a total of 52 different control sequences.

**Individual full miRNA target sequences**

This sublibrary tests how a single full miRNA target site alters stability and the reliability of different data sources for miRNA sequences. Human miRNA sequences were taken from miRBase (version 22). MicroRNAs longer than 21 nt were truncated to 21 nt from the 3' end. Sequences shorter than 21 nt were padded with Us at their 3' end to reach a total length of 21 nt. A miRNA target is the reverse complement of these homogenized miRNA sequences. Targets containing a canonical poly(A) signal (AAUAAA) were filtered. We used almost all (856) high confidence human miRNAs in miRBase v22 with the exception of those containing canonical poly(A) signals in their targets. We added all sequences in MirGeneDB (166 out of 506 total) that were annotated as low confidence in miRBase. We also sampled 293 additional low confidence miRNAs from miRBase that are not in MirGeneDB, prioritizing miRNAs with large expression values in any of our chosen cell lines.

**Assessing context effects and the impact of secondary structure**

We chose ten miRNAs that cover a range of expression values in our cell lines (let-7a-5p, miR-16-5p, miR-19b-3p, miR-21-5p, miR-22-3p, miR-23a-3p, miR-24-3p, miR-31-5p, miR-107, miR-365a-3p). We embedded full targets for these miRNAs into two types of context sequences: First, we used the top 30 context sequences least variable in stability generated from our controls in Library 1 (see Controls). Second, we inserted 30 short sequences with varying amounts of complementarity to each of these miRNA targets starting 26 nt upstream of the 5' end of the miRNA target into the main context sequence. The degree of complementarity ranges from a block of 5 complementary bases to full complementarity to the miRNA target site. The distribution of $\Delta\Delta G$ values was calculated using NUPACK python package (version 4.0.0.20) (*73*) by deducting the free energy of the two individual strands (the 3'UTR and the miRNA) from the free energy of their complex.

**Individual mutated miRNA sequences**

We chose 11 miRNAs to mutate: 10 microRNAs (miR-16-5p, miR-19b-3p, miR-21-5p, miR-22-3p, miR-23a-3p, miR-24-3p, miR-31-3p, miR-31-5p, miR-107, miR-365a-3p) expected to have little crosstalk and one miRNA (let-7a-5p) with strong expected crosstalk. These microRNAs were chosen because they have highly variable expression values across our cell lines, allowing us to test the impact of mutations at different microRNA expression levels. We generated a total of 1092 mutated microRNA targets: a) a single randomly chosen non-wobble mutation at all possible positions (252 total), b) all possible mutations leading to a single wobble base pair with the miRNA (132 total), c) insertion of an A at position 1 of the seed site (4 total), d) 704 target sites with multiple base changes distributed across the miRNA target for a total of two (231 total), three (187 total), four (132 total), five (99 total), and six (55 total) mutations.

##### Full miRNA target site repeats

We chose 100 miRNAs to generate constructs containing two to six repeats of a full target site. We included all miRNAs for which we tested mutations and all miRNAs of the 5p arm of the let-7 family. To make sure we cover the full range of miRNA expression values across our cell lines, we then classified miRNAs into buckets according to their maximum expression levels across our cell lines: bucket 1 (max > $10^4$ tpm), bucket 2 ($10^4$ tpm ≥ max > $10^3$ tpm), bucket 3 ($10^3$ tpm ≥ max > $10^2$ tpm), and bucket 4 ($10^2$ tpm ≥ max). We chose 14 miRNAs in bucket 1, 46 in bucket 2, 10 in bucket 3, and 5 in bucket 4. This choice ensures that we test all the most impactful miRNA targets across our cell lines.

##### Full miRNA target site combinations

To be able to distinguish different models of target site interactions, it is necessary to have constructs in which the different regulating miRNAs have similar expression levels. For example, in the additive model, if the ratio of expression between two cognate miRNAs for two target sites is 10, their impact on the stability fold change will have approximately the same ratio. Thus, it would be difficult to determine whether the less expressed miRNA has any impact at all. We therefore divided miRNAs into buckets depending on their mean expression across our cell lines: below $10^{2.5}$ tpm, between $10^{2.5}$ and $10^3$ tpm, between $10^3$ and $10^{3.5}$ tpm, and above $10^{3.5}$ tpm. When choosing multiple miRNA targets for a single UTR, we chose targets from the same bucket to maximize the odds of choosing miRNAs with similar expression levels. We started by generating 30, 30, and 20 combinations of two different miRNAs for buckets 2, 3, and 4, respectively. We then sampled an additional miRNA from the same bucket and added it to the previously created designs with one less target site, making sure to discard any duplicate designs with the same target sites. We repeated this procedure to generate constructs with between two and six different microRNA target sequences.

##### Design objectives for tailored stability profiles

We used five types of binary designs (active in one or two cell lines or inactive in one, two, or three cell lines) with target relative stability values of 0 or 1. For binary designs, we generally created 4 designs per cell line and miRNA target site number in two design rounds. In the first round, we generate 2 designs. In the second design round, 2 miRNA targets that were most often used in the first design round were excluded to create 2 additional designs. This prevents the algorithm from choosing identical sites for every single design but also makes it likely that the second set of designs is worse than the first. We also used four types of designs with graduated target stability values: uniformly random values between 0 and 1 sampled independently for each cell line, a range of fixed uniformly spaced values between 0 and 1 randomly assigned to the different cell lines, values distributed uniformly on a log scale between 0.05 and 1 sampled independently for each cell line, and fixed logarithmically spaced values between 0.05 and 1 randomly assigned to the different cell lines. In the evaluation, these are uniformly treated as graduated designs. For graduated designs, we created 93 designs per design type and miRNA target site number with 1 or 2 designs per target pattern (a particular distribution of target stabilities over all cell lines).

##### Model fitting for the design of tailored stability profiles

Sequences for tailored stability profiles were assayed as part of Library 2. Thus, all model design and fitting was done using Library 1 data and the insights gained from it. We derived the constant $\frac{k_{deg}}{k_{on}}$ for the transfer function by fitting the data of Library 1 for the four cell lines (HEK293T, HeLa, MCF7, SKNSH) we measured for that library. We heuristically filtered for crosstalk by first identifying all potentially crosstalking miRNAs as those with no mismatches in the core seed (positions 2 to 7) and up to 4 mismatches total. This crosstalk filtering is different from the one applied for the updated model as we did not have access to our measured crosstalk data at this point. A miRNA target was excluded when its crosstalking miRNAs were expressed to a level of at least 400 tpm and more than 1.35 times its own value in any cell line. We also heuristically filtered false positives in the microarray data. For each cell line, we sorted all miRNA targets with a stability larger than $10^{-0.5}$ and positive deviation by their deviation from the model. miRNAs in the top 30% in at least three out of four cell lines (8 miRNAs total) were filtered. We fitted our transfer function to the filtered data, yielding $k_{deg}/k_{on}=10^{3.65}$, which agrees well with the value later derived from filtered Library 2 data ($k_{deg}/k_{on}=10^{3.68}$).

##### The genetic design algorithm

We used an evolutionary algorithm to create designs with 4, 5, and 6 miRNA target sites for each design type. We started with 300 sets of miRNA sites randomly sampled from all high-confidence miRNAs and ran the algorithm for 30 generations. In each generation, we first predict the stability pattern across our cell lines using the additive model and the transfer function derived from fitting the data for Library 1. We then evaluate the fitness of all designs. A set of 300 new designs was created by selecting 2 parents by tournament selection with a size of 3, merging the designs at a random position, and then randomly changing one of the miRNA targets with a 20% chance. We usually defined the fitness *f* of a design as the inverse of the weighted mean square error (wmse) between the target and the predicted stabilities $s_{target}$ and $s_{pred}$: $f_{wmse} = \left(\sum_{cell\ lines\ i=1}^{n} w_i (s_{target,i} - s_{pred,i})^2 / n\right)^{-1}$. For graduated expression patterns, the weighting was uniform across cell lines. For the binary designs, we weighted the error in the target cell lines 5 times for designs with one or two target cell lines and 3.33 times for designs with three target cell lines. For the designs meant to be active in a single cell type, we also created separate designs with a different fitness criterion $f_{tsi}$ in which we multiply the tissue-specificity index (*62*) of the predicted stability with the stability in the target cell line $s_{target}$: $f_{tsi} = s_{target}\left(\sum_{cell\ lines\ i=1}^{n} 1 - s_i / s_{max,\,1\ldots n}\right)/(n-1)$. This alternative criterion more strongly emphasizes high stability in the target cell line over reduction of stability in the other cell lines. For the graduated expression patterns with logarithmically distributed stability values, we also evaluated the mse on a log scale.

<u>Library 3 design</u>

Each sequence has a length of 200 nt, of which 18 nt were fixed at the 5' end (ACGACGCTCTTCCGATCT; part of the TruSeq Read 1 sequence) and 18 nt (CTCTGGATTTGCAACCGA) at the 3' end, leaving a total of 164 variable nts, same as for Library 2. See **Table S11** for a list of sublibraries and **Data S5** for library sequences.

**Individual miRNA targets and repeats**

Mouse miRNA sequences were taken from miRBase (version 22). MicroRNAs longer than 21 nt were truncated to 21 nt from the 3' end. Sequences shorter than 21 nt were padded with Us at their 3' end to reach a total length of 21 nt. A miRNA target is the reverse complement of these homogenized miRNA sequences. We started with an atlas of 61 mouse immune cell populations generated via qPCR array by Rose et al. (*66*) containing 923 measured miRNAs. We first removed miRNAs that have an expression level below $10^3$ tpm in all of the samples. We then filtered to miRNAs that are either high confidence in miRbase or in MirGeneDB. We then removed targets containing poly(A) signals (AAUAAA or AUUAAA), homopolymers longer than six bases, with >75% GC content, or that are expected to be dominated by crosstalk as calculated using the model derived for Library 2, yielding 163 microRNA targets. We additionally included miRNAs with a mean expression above 3000 tpm in the Tabula Muris Senis or Brain Aging Mouse datasets of miRNATissueAtlas 2025 (*74*), yielding another 50 microRNA targets for a total of 213 targets. We tested three, four and five repeats for each of these miRNA targets. We also added constructs with 1, 3, 5, and 7 seed targets for each unique seed (first 8 bases) of the 163 miRNAs of the first set (124 seed sites). Although we do not discuss the associated data here, it is included in the provided supplementary files.

**Combinations of microRNA targets**

To generate designs with combinations of targets, we divided miRNAs into buckets depending on their expression in the Spleen_CD8+ T cells KLRG1+CD127- +LisOVA, day8 sample of the Rose et al. dataset: below $10^3$ tpm, between $10^3$ and $10^{3.5}$ tpm, between $10^{3.5}$ and $10^4$ tpm, and above $10^4$ tpm. We then sampled 5 unique miRNAs from the bucket 26, 40, 40, and 30 times, respectively, for a total of 136 combinations. We additionally generated various designs with up to 5 target sites to potentially distinguish between human tissues, mouse immune cells, mouse brain cells, and mouse tissues (774 total) using the design algorithm described above for Library 2.

**<u>Library Cloning</u>**

**Table S2** lists all oligos used for cloning and library preparation. Libraries were ordered as oligo pools from Twist. We resuspended the oligos to 5 ng/µl in 10 mM Tris-HCl, pH 8.0. We amplified the library using Phusion High-Fidelity PCR Master Mix with HF Buffer (NEB, M0531L) using 0.25x EvaGreen (Biotium, #31000), 1 ng/µl resuspended library, and 0.5 µM of oligos 53 and 54 (Library 1) or oligos 51 and 52 (Library 2 and Library 3). We initially ran a test qPCR amplification for 25 cycles in a total of 10 µl to determine the optimal cycle number before the end of exponential amplification. Then we ran a larger reaction with this cycle number. The amplification protocol was 96°C for 40s, cycles of 96°C for 15s, 61°C for 20s, 72°C for 20s until the end of the exponential phase (approximately 10 cycles for all libraries), then a final extension at 72°C for 8 min. The DNA was purified using 1.5x SPRIselect (Beckman Coulter, B23319) bead cleanup according to the manufacturer's instruction.

**Table S6** lists the used plasmids. We used plasmid 1 (Addgene #176640 (*35*)) as the base plasmid for Library 1. For Library 2, BsaI cutting sites were inserted into this plasmid via overhang PCR to create plasmid 4 (oligos 49 and 50). For Library 3, we used plasmid 8 as the base cloning plasmid which includes the murine stem cell virus backbone for retroviral transgene production (*75*). The base plasmids were digested with BmtI and XbaI (Library 1), BsaI (Library 2) or HindIII (Library 3) and purified on a 1% agarose gel using Monarch® DNA Gel Extraction Kit (NEB, T1020L). We used a Gibson Assembly reaction (NEBuilder® HiFi DNA Assembly Master Mix, NEB, E2621L) using 800 ng of plasmid digest and 210 ng of amplified library at 50°C for 1h to assemble the libraries. The assembled plasmids were purified using a 1x SPRIselect bead cleanup.

We transformed 480 ng of assembled library into a vial of NEB® 10-beta Electrocompetent E. coli (NEB, C3020K). After resuspension in 1 ml of SOC and incubation at 37°C for 1h, we plated a 1:100,000 dilution to determine the transformation efficiency and grew the remaining cells in 200 ml of LB supplemented with 50 µg/ml kanamycin (Library 1 and Library 2) or 100 µg/ml carbenicillin (Library 3) overnight. The libraries were purified using QIAGEN Plasmid Maxi Kit (Qiagen, 12162).

#### **Individual construct cloning for cell line flow cytometry**

Starting from plasmid 1 digested with BmtI and XbaI, we inserted the main context sequence for Library 1 via Gibson assembly with oligo 26. The backbone plasmid for insertion of miRNA target sites was created from this plasmid by an overhang PCR with oligos 27 and oligo 28. This backbone plasmid was then digested with BsaI, and the individual constructs were created by ligation of the digested plasmid with oligos 29 to 48. The resulting constructs precisely match the associated sequences in Library 1.

#### **Cell culture for high-throughput stability measurements in cell lines**

HEK293T, HeLa, HUH7, MCF7, K562, SKNSH, JEG3, Tera1, and 3T3 cells were purchased from ATCC. A549 cells were a kind gift from Jesse Bloom. HaCaT cells were a kind gift from Paul Nghiem. PC3 cells were a kind gift from Andrew Hsieh. P2C2 cells were originally sourced from Ellen Rothenberg and have been described (*76*). HEK293T, HeLa, MCF7, A549, HUH7, 3T3 and HaCaT cells were cultured in DMEM supplemented with 10% FBS and 100 U/ml Penicillin/Streptomycin (ThermoFisher 15140122). SKNSH and JEG3 cells were cultured in EMEM supplemented with 10% FBS and 100 U/ml Penicillin-Streptomycin. K562 and PC3 cells were cultured in RPMI supplemented with 10% FBS and 100 U/ml Penicillin-Streptomycin. Tera1 cells were cultured in McCoy's 5A medium supplemented with 15% FBS and 100 U/ml Penicillin-Streptomycin. P2C2 cells were cultured in RPMI supplemented with 10% FBS, 1% MEM Non-Essential Amino Acids Solution (100X), and 0.5 mg/ml Penicillin-Streptomycin-Glutamine.

#### **MicroRNA expression measurements (improved sequencing data)**

The same RNA samples measured using microarrays in Alles et al. (*37*) were used for miRNA sequencing most human cell lines in this study. Cell culture conditions and the RNA isolation procedure have been described there. For HeLa, JEG3, and Tera-1, we additionally used the same total RNA extracted during the ratiometric stability assay for Library 2. For mouse cell lines and CD8 T cells, total RNA was extracted using the Monarch Total RNA Miniprep Kit (NEB). Small RNA expression libraries were generated using MGIEasy Small RNA Library Prep Kit (MGI Tech) according to the manufacturer's protocol. Briefly, 100 ng of total RNA were used for 3’ adapter ligation. After removal of unused adapter by digestion, 5’ adapters were ligated and the product was reverse transcribed using uniquely barcoded RT primers. The cDNA was PCR amplified for 21 cycles and PCR products derived from miRNAs were size selected by gel electrophoresis using Novex 6% TBE PAGE gels (Invitrogen). PCR products were pooled into a single library, circularized, and sequenced using DNBSEQ-G400RS High-throughput Sequencing Reagent Set (G400 sRNA FCL SE50) on a DNBSEQ-G400RS sequencer (both MGI Tech).

Fastq files were trimmed and quantified using the miRMaster 2.0 pipeline (*77*). The adapter sequence used for trimming was AGTCGGAGGCCAAGCGGTCTTAGG with a minimum overlap of 10 nt. We set a maximum edit distance of 1 nt for adapter matches and no ambiguous nucleotides were allowed. For each read 3 nucleotides were trimmed from the leading and trailing ends. Trimming was further performed with a sliding window of 4 nt with a phred score quality threshold of 20. Read lengths smaller than 17 nt were removed. Reads were collapsed and mapped against GRCh38 or GRCm38 with a maximum mismatch of 1 using bowtie 1.1.2 using the options –mm -v1 –m 100 –best –strata –fullref. MiRNA and isomiR quantification was performed against the *H. sapiens* or *M. musculus* miRNA set from miRBase v22.1.

##### Flow cytometry for cell line control experiments

We seeded 100,000 HEK293T or HeLa cells in 24-well plates 24 h before transfection. We transfected 500 ng total of plasmid in each well. To screen the effect of plasmid concentration on miRNA activity, we used between 16.7 and 200 ng of reporter plasmid, 100 ng of a CMV-mCherry transfection control plasmid (plasmid 6), and enough pUC19 filler plasmid to reach 500 ng total. To measure the impact of different miRNA target sites, we used 30 ng (HEK293T) or 60 ng (HeLa) reporter plasmid, 100 ng of a CMV-mCherry plasmid (plasmid 6), and enough pUC19 filler plasmid to reach 500 ng total. Transfections were carried out using Lipofectamine™ 3000 Transfection Reagent (ThermoFisher, L3000001) according to the manufacturer's instructions. We exchanged media 4 h after transfection. Two days later, we detached cells with TrypLE, added media, then centrifuged for 5 min at 210 g. Cells were resuspended in 1 ml of PBS. We measured 200 µl of cell suspension either undiluted (HeLa) or diluted 1:1 in PBS (HEK293T) using an Attune NxT Flow Cytometer (ThermoFisher). We first gated cells on SSC-A and FSC-A (**Fig. S5B**), then on the mCherry fluorescence such that almost all non-transfected cells from a negative control were excluded (**Fig. S5C**). We calculated the median GFP fluorescence of the gated cells, subtracted the median GFP fluorescence of an untransfected negative control, then normalized to a positive control of the same plasmid without miRNA target sites (**Fig. S5E**). No compensation was required because fluorophores are spectrally separated.

##### Cell line plasmid library transfection and electroporation

We measured two replicates for each cell line. For all adherent cell lines, we seeded between 350,000 and 5 million cells two days (Tera1 cell line) or one day (all other cell lines) before transfection. The precise transfection parameters per cell line can be found in **Table S1**. Between 2.5 and 15 µg of plasmid library were transfected with Lipofectamine™ 3000 Transfection Reagent (ThermoFisher, L3000001) according to the manufacturer's instructions. We exchanged media 4 h after transfection. 48 hours later, we detached cells with TrypLE and added media..

For K562, 5 million cells were transfected with 10ug of the 3’UTR library 2 using the Neon Transfection System (Invitrogen MPK5000) following the manufacturer’s instructions, with E2 buffer, 1350V, and 3x 10ms pulses. Electroporated cells were placed in 6 ml RPMI + 10% FBS (no P/S). We processed the cells 48 hours after electroporation.

For all cell lines, cells in media spun down for 5 min at 210 g. We resuspended the cells in 1 ml of cold PBS and placed them on ice. We took 25 µl of each sample, added 175 µl of PBS, and measured the GFP fluorescence on the Attune NxT flow cytometer to determine cell numbers and transfection efficiencies.

#### **mRNA purification and library amplification for ratiometric stability data in cell lines**

We used the Monarch Total RNA Miniprep Kit (NEB, T2010S) to purify total RNA. We eluted in 100 µl of nuclease-free water and determined concentrations using a NanoDrop 2000c (Thermo Fisher). mRNA was purified using the NEBNext® Poly(A) mRNA Magnetic Isolation Module (NEB, E7490L). We eluted in 18 µl of nuclease-free water and performed reverse transcription (RT) using 0.75 µM oligo 1 (Library 1) or oligo 11 (Library 2, Library 3), 0.5 mM dNTPs (NEB), 0.5 U/µl SUPERase·In™ RNase Inhibitor (ThermoFisher, AM2696), and 10 U/µl Maxima H Minus Reverse Transcriptase (ThermoFisher, EP0752). Before adding the RT buffer and enzyme, the mix was incubated at 65°C for 5 min. Reverse transcription was carried out at 50°C for 15 min, then 85°C for 5 min. We added 10 U RNase I (ThermoFisher, EN0601) and 5 U RNase H (NEB, M0297S) and incubated at 37°C for 30 min. cDNA was purified using DNA Clean & Concentrator-5 (Zymo, D4014) with 7x binding buffer, eluted in 10 µl nuclease-free water, and stored at -20°C until library amplification.

Libraries were amplified using KAPA HiFi HotStart ReadyMix (Roche, KK2602) with 0.5 µM of each index primer (Oligos 2 to and 8 and 12 to 25) and 1x EvaGreen (Biotium, #31000). The protocol was 95°C for 3 min, a variable number of cycles of 96°C for 25s, 69°C for 15s, 72°C for 30s, and a final extension at 72°C for 5 min. We ran a pilot qPCR for 30 cycles with 1.09 µl of library in a total of 5 µl to determine the appropriate cycle number. The full library was amplified in 50 µl until the last cycle of exponential amplification. The PCR reaction was run on a 1.8% agarose gel at 120V for 35 min. The band matching the expected size was cut out, purified using Monarch® DNA Gel Extraction Kit (NEB, T1020L) and eluted in 20 µl of elution buffer. Concentrations were determined using a Qubit™ dsDNA Quantification Assay (ThermoFisher Q32851).

To introduce unique molecular identifiers (UMIs) for the plasmid DNA library, library amplification was preceded by two cycles of PCR amplification with the reverse transcription primer (oligo 1 or 11) and the library amplification i5 primers (oligos 2 and 3 or 12 and 13). The

DNA was purified using DNA Clean & Concentrator-5 (Zymo, D4014), eluted in 10 µl, and amplified as above.

**Time course-based stability measurements**

To compare mRNA amounts across different sequencing libraries, a small set of 10 constructs with known sequences was used as spike-in controls. Each construct was individually cloned into the same 3'UTR library plasmid backbone to be compatible with their sequencing library preparation pipeline. After cloning, the GFP cassette was PCR-amplified with a forward primer containing a T7 promoter (gcgaaattaatacgactcactataGGGCTCTTCCTCATCTCCGGGC) binding upstream of the 5'UTR, and a reverse primer containing a large polyT segment ([T70]GTTGTTAACTTGTTTATTGCAGCTT) binding downstream of the 3'UTR. Amplicons were purified and mixed at concentrations of 12, 12, 1.2, 1.2, 0.12, 0.12, 0.012, 0.012, 0.0012, and 0.0012 ng/uL. Finally, spike-in *in vitro* RNA was synthesized using the HiScribe T7 RNA synthesis kit (NEB E2040S).

For the time-course experiment, 5 million K562 cells were transfected with 10 ug of the 3'UTR library 2 using the Neon Transfection System (Invitrogen MPK5000) following the manufacturer's instructions, with E2 buffer, 1350V, and 3x 10ms pulses. Electroporated cells were placed in 75mL RPMI + 10% FBS (no P/S). 48 hours later, 5mL were extracted and successful transfection was verified via flow cytometry. Then, 2 samples of 6 ml were extracted and Actinomycin D (Thermo A7592) was added to the remaining culture to 5 ug/ml. Extracted samples were placed in 15 ml conical tubes each, centrifuged at 100g and room temperature for 5 minutes, and the supernatant was discarded (no aspiration). Then, cells were resuspended in 1 ml ice cold PBS, centrifuged at 1000g 4°C, and the supernatant was discarded. The resulting pellet was transferred immediately to -80°C. At 30 min, 1h, 2h, 4h, 8h, 12h, and 24h after the addition of ActD, 6 ml of culture were extracted from the culture and centrifugation and sample freezing was performed as above. Since sample preparation at room temperature took ~10 minutes, sample extraction was actually performed 10 minutes before the indicated timepoint. Two experimental replicates were performed.

For sequencing library preparation, we first determined the appropriate amount of IVT RNA spike-in to add using one of the t=0 samples, with the goal of having the most abundant spike-in species be at ~2x the concentration of the most abundant cell mRNA species at t=0. To do this, we removed the cell pellet from the freezer, equilibrated it to room temperature, lysed it with 800 ul of the NEB RNA miniprep kit's lysis buffer, split into 8 tubes, added 5 ul of a different spike-in dilution to each (from 1x to 1e6x dilutions, and one sample receiving no spike-in), and proceeded with library preparation until the saturation qPCR step as above. qPCR amplification curves were used to estimate the relative amount of cell library mRNA compared to spike-in RNA, and the appropriate amount of spike-in RNA to add to a whole sample was calculated. With this information, all timepoint samples were processed by removing tubes from the freezer, lysing as above (no splitting), adding spike-in to every tube, and proceeding with the standard library preparation and sequencing protocol as for the other libraries.

**Polysome profiling measurements**

Before seeding, plates for HEK293T were coated with Poly-D-Lysine to improve adherence during the washing steps. HEK293T and HeLa cultures at ~30% and ~80% confluency in 10 cm plates were transfected with a mixture of 7.5ug 3'UTR + 7.5ug 5'UTR libraries using Lipofectamine 3000 as above. 24h later, cells were lysed in the presence of cycloheximide as previously described (*60*, *61*). In short, we replaced plate media with 5 ml room temperature Wash+CHX buffer (RNase-free DPBS + 100 ug/mL cycloheximide), incubated at 37°C for 5 min, replaced buffer with 5 ml ice cold Wash+CHX buffer, placed plate on ice and removed buffer, added 300uL cold lysis buffer (1x salt solution, 1% Triton X-100, 1mM DTT, 0.2 U/uL SUPERase-In, 100 ug/ml cycloheximide, and water to 300 ul; where 10x salt solution contains 100 mM NaCl, 100 mM $MgCl_2$, and 100 mM Tris-HCl pH 7.5 in RNase-free water), scraped cells, transferred to a 1.5ml centrifuge tube, incubated on ice for 10 min, triturated cells by aspirating and pushing 10 times with a 25-G needle and syringe, centrifuged at 16,000 g for 5 min at 4°C, transferred supernatant to a different tube, added DNase I to 0.005 U/ul, incubated for 30 min on ice, and stored at -80°C.

Polysome profiling was performed as previously described (*60*, *61*). Briefly, ultracentrifuge tubes were prepared by layering 5.4 ml 20% sucrose solution (20% sucrose plus 100 mM KCl, 20 mM HEPES pH 7.2 and 10 mM $MgCl_2$) on top of 5.4 ml of a similar 55% sucrose solution, covering the opening with parafilm, and leaving tubes on their side overnight at 4°C. The next day, tubes were returned to a vertical position, incubated for 1-2h, and cell lysate was carefully pipetted on top of the sucrose gradient. ~50 ul of cell lysate was left in its original tube and stored back to be used as a control later. Ultracentrifugation was performed for 3h at 151,000g using a Beckman SW-41 Ti rotor. Finally, ~25 fractions were collected, and fractions corresponding to free RNA and all visible polysome peaks were used in downstream processing (**Fig. S27A-B**).

For sequencing library preparation, a portion of the remaining total cell lysate was used to calibrate the amount of spike-in to be added, with the goal of having the most abundant spike-in species be at ~2x the concentration of the most abundant cell mRNA species in the average fraction tube. To this end, we removed 10 ul lysate, added 21 ul $H_2O$, and split into 6 tubes, 5 ul each. Next, we added 5 ul of spike-in dilution to each tube (10x, 100x, 1e3x, 1e4x, 1e5x, and no spike-in) for a total of 10 ul. For lysis, 250 ul Trizol (Invitrogen 15-596-026) was added and the mixture was vortexed and incubated at room temperature for 5 minutes. Then, 50 ul chloroform was added, vortexed, and incubated for 3 minutes. The mixture was centrifuged at 16,000 rcf for 10 minutes, the supernatant (~125uL) was transferred to another RNase-free tube, and an identical amount of 100% ethanol was added. RNA purification was then completed with the Zymo RNA C&C-5 kit (Zymo R1013), by transferring the mixture to a column and centrifuging, adding 400 ul RNA prep buffer and spinning for 30 sec, adding 700 ul RNA wash buffer and spinning for 2 minutes, and eluting in 53uL RNase-free water. Library preparation, from mRNA isolation using magnetic beads to saturation qPCR, proceeded as described above. qPCR curves were used to calculate the appropriate amount of spike-in as in the time-course experiment. With this information, all fractions were processed by removing tubes from the freezer, adding spike-in to every tube, lysing as above with 500 ul of Trizol per 500 ul of lysate, and using appropriately scaled amounts of chloroform and ethanol (no splitting). After purification, we proceeded with the standard library preparation and sequencing protocol. Fractions corresponding to the same polysome peak were combined by passing Trizol + chloroform + ethanol mixtures through the same Zymo column as appropriate. Note that mRNA isolation was only performed on the total lysate and the free ribosome fraction under the assumption that ribosome-bound fractions are already enriched for mRNA.

#### Mouse husbandry

All mice used in this study were homozygous for the *Tcf7*-YFP reporter (*78*) and the $H11^{Cas9}$ allele (The Jackson Laboratory Strain #:027650). Both male and female mice aged 4 to 5 months were used for experiments. Experimental groups were age and sex matched. All mice were housed, maintained, and euthanized in accordance with Institutional Animal Care and Use Committee (IACUC) guidelines for the University of Washington.

#### MSCV virus production

MSCV retrovirus was made for Library 3. HEK293T cells cultured in DMEM supplemented with 10% FBS, 1 mM Sodium Pyruvate, and 0.5 mg/ml Penicillin-Streptomycin-Glutamine were seeded on either 10 cm or 6-well tissue culture treated plates the day before transfection. When they reached ~70% confluence, the HEK293T cells were transfected with a mixture of Library 3 and pCL-Eco (Addgene #12371) plasmids using the FuGENE® 6 Transfection Reagent (Promega). Approximately 24 hours later, the HEK293T media was exchanged for fresh growth media. 48 hours after media change, the supernatant was removed from the HEK293T culture and filtered through a 0.45 uM PES syringe filter. The filtered viral supernatant was stored on ice until used for CD8+ T cell transductions which were performed on the same day.

#### Mouse CD8 T cell isolation, transduction, culturing, and differentiation

Spleens were harvested and ground between two sterile microscope slides into a single-cell suspension in HBH solution (HBSS, 10 mM HEPES, 0.5% BSA, pH 7.4). The suspension was filtered through a 40 um nylon mesh filter and pelletted via centrifugation at 300g for 5 min. Cells were resuspended in 3 ml red blood cell (RBC) lysis buffer per spleen (150 mM $NH_4Cl$, 10 mM $NaHCO_3$, 1 mM EDTA) and incubated at room temperature for 5 min, then quenched with HBH at 2x the volume of RBC lysis buffer. Cells were spun down at 300g for 5 min, resuspended in HBH with 2.4G2 Fc blocking solution, counted, and incubated on ice for 30 min. Splenocytes were enriched for CD8+ T cells via a CD8a+ T cell Isolation Kit, mouse (Miltenyi #130-104-075), with the antibodies and microbeads scaled down to 70% of the specified amounts in the manufacturer's protocol. One to three spleens were loaded onto each LS column (Miltenyi #130-042-401).

One day before spleen harvest and CD8+ T cell isolation, tissue culture treated flat bottom culture plates were coated with 1 ug/ml anti-CD3e (Tonbo, #40-0031-U100) and 0.5 ug/ml anti-CD28 (Tonbo, #40-0281-U100) antibodies in PBS, wrapped with parafilm, and incubated at 4°C overnight. Antibody coated plates were washed twice with PBS before plating isolated CD8+ T cell cultures at 1-1.7 million cells/ml of T cell media (RPMI 1640 with L-glutamine media supplemented with 10% FBS, 20 mM HEPES, 1 mM Sodium Pyruvate, 1% MEM Non-Essential Amino Acids Solution (100X), 50 μM beta-mercaptoethanol, and 0.5 mg/ml Penicillin-Streptomycin-Glutamine). For all experimental conditions, CD8+ T cells were cultured in TCM supplemented with 10 ng/ml IL-2 (PeproTech, # 200-02) at 37°C, 5% $CO_2$ through initial activation and transduction.

In the first CD8+ T cell experiment, one day before transduction, non-tissue culture treated plates were coated with 12.5 ug/ml RetroNectin (Takara, #T100B) in PBS, wrapped with parafilm, and incubated at 4°C overnight. The next day, RetroNectin-coated plates were blocked with Bovine Serum Albumin (BSA) (Fisher BioReagents™ #BP1600-100) by aspirating the RetroNectin solution, adding 2% w/v BSA in PBS, and incubating at 37°C for 30 min. BSA-blocked plates were rinsed once with PBS, then coated with retrovirus by adding filtered retroviral supernatant (as described above) and spinning in a centrifuge at 3000g, 32°C for 2 hrs. No sooner than 24 hrs after plating the purified CD8+ T cells on the initial activation plate, the cells were removed from activation, pooled, and counted. After retrovirus coating, the supernatant was aspirated from the plates and they were rinsed once with PBS. CD8+ T cells in TCM supplemented with 10 ng/ml IL-2 were transduced by transferring them to virus-coated plates at a concentration of ~1x$10^6$ cells/ml and spinning at 800g, 32°C for 30 min, then incubating overnight at 37°C, 5% $CO_2$. On the day of transduction, tissue culture treated activation plates were coated with 1 ug/ml anti-CD3e and 0.5 ug/ml anti-CD28 antibodies in PBS, wrapped with parafilm, and incubated at 4°C overnight. The next day, the initially activated and transduced CD8+ T cells were pooled, split between two culture conditions (rested and exhausted), spun down at 300g for 5 min, and resuspended in TCM supplemented with the appropriate cytokines (rested: IL-15 (PeproTech # 210-15) at 50 ng/ml; exhausted: IL-2 at 10 ng/ml and IFN-α (Biolegend #752804) at 10 ng/ml). Rested samples were plated on non-antibody coated tissue culture plates and harvested two days later for downstream analyses. Exhausted samples were plated on antibody-coated plates, passaged every two days onto freshly-coated activation plates, and harvested five days later for downstream analyses.

In the second CD8+ T cell experiment, one day before transduction, non-tissue culture treated culture plates were coated with 12.5 ug/ml RetroNectin, 1 ug/ml anti-CD3e, and 0.5 ug/ml anti-CD28 in PBS, wrapped with parafilm, and incubated at 4°C overnight. The next day, RetroNectin/antibody-coated plates were blocked with BSA, rinsed, coated with virus, and plated with CD8+ T cells as described above. On the day of transduction, tissue culture treated activation plates were prepared as described above. The next day, activated and transduced CD8+ T cells were pooled, split between four culture conditions (rested, early activated, late activated, and exhausted), spun down at 300g for 5 min, and resuspended in TCM supplemented with the appropriate cytokines (rested: IL-15 at 50 ng/ml; early and late activated: IL-2 at 10 ng/ml; exhausted: IL-2 at 10 ng/ml and IFN-α at 10 ng/ml). Rested samples were plated on non-antibody coated tissue culture plates. Early and late activated as well as exhausted samples were plated on antibody-coated plates. Rested and early activated samples were harvested two days later for downstream analyses. Late activated and exhausted samples were passaged every two days onto freshly-coated activation plates and harvested six days later for downstream analyses.

**Flow cytometry measurements for mouse CD8+ T cells**

CD8+ T cell samples were prepared for flow cytometry measurement by spinning down at 300g for 5 mins in a 96-well round bottom plate, resuspending in 2.4G2 blocking solution, and incubating on ice for 15-30 mins. Antibody stains for PD-1 and TIM-3 were added at appropriate concentrations (1:400 PD-1: e450 Invitrogen # 48-9981-82; 1:100 TIM-3: APC Biolegend # 119706) and the antibody/cell suspension was incubated on ice for an additional 15-30 mins.

Cells were spun down at 300g for 5 mins and resuspended in HBH prior to acquisition on an Attune NxT Flow Cytometer (ThermoFisher Scientific). CD8+ T cell flow cytometry data were analyzed using FlowJo (BD) software.

**P2C2 and mouse CD8+ T cell mRNA, gDNA, and viral RNA library preparation**

Mouse CD8+ T cell and P2C2 cultures were spun down for 3 min at 500g at 4°C in a swinging bucket centrifuge. To extract library mRNA, we proceeded with the same RNA purification protocol as described above for human cell lines. To isolate gDNA, we used the Quick-DNA Kit (Zymo, D3024) and eluted in 52 ul. A UMI was added as described above for plasmid DNA with a final concentration of 30 ng/ul gDNA in a 200 ul PCR reaction. The reaction was purified using a two-sided size selection with SPRI beads (0.6x - 1.2x) and eluted in 15 ul. Following this, we proceeded with library amplification as for mRNA libraries. To isolate viral RNA, we concentrated viral samples using Retro-X Concentrator (Takara Bio, #631455). We started with 30 ul of concentrated viral RNA, added 800 ul of TRIzol® reagent (ThermoFisher) and incubated at RT for 5 min. We added 160 ul of chloroform and incubated at RT for 3 min. We then spun at 16000g for 10 min and transferred the supernatant (~450 ul) to a new tube. The supernatant was then further purified using the RNA Clean & Concentrator-5 kit (Zymo, R1013) and eluted in 18 ul water. At this point, we proceeded with the NEBNext® Poly(A) mRNA Magnetic Isolation Module (NEB, E7490L) and downstream steps as described above for cell line mRNA.

**Library sequencing and data processing**

Ratiometric data for Library 1 was sequenced in-house on a NextSeq 500/550 Mid Output kit (150 cycles, Illumina) with custom primers (Oligos 9 and 10). Ratiometric data for Library 2 and Library 3 as well as time-course stability data and polysome profiling data for Library 2 were sequenced on a NovaSeq X Plus (Illumina) using a paired-end 300 cycle kit by Novogene. Cell line and replicate data was split according to the i5 and i7 indices. The UMI was extracted and added to the read name. Libraries were aligned to the reference sequences using BWA-MEM. Alignments were filtered using a custom python script. First, we removed ambiguous alignments: Reads were discarded when one of the two reads failed to align, when both reads had multiple top-scoring alignments, or when one read aligned uniquely but did not match any of the top alignments of the other strand. We then filtered to at most one total insertion or deletion or four substitutions relative to the aligned reference sequence. The alignments were sorted and indexed using samtools. UMIs were deduplicated using UMI-tools with unique matching. Alignments for each reference were counted using a custom python script.

**Ratiometric stability data analysis**

All analyses are performed with custom Python code.

Fold-change values were calculated from the count data of the two replicates using PyDESeq2 (*79*). The calculation was performed relative to plasmid library counts for stability assays in cell lines (Library 1 and Library 2, HEK293T and 3T3 data for Library 3) as well as to assess biases and MSCV virus production. For MSCV integration in P2C2 and mouse CD8 T cells, the calculation was performed relative to gDNA counts.

#### Ratiometric stability data analysis (Library 1 and 2)

##### Filtering, calculation of stability values and normalization

We filtered all sequences with less than 100 combined counts in the two plasmid DNA library replicates (one sequence out of 10,001 total unique sequences in Library 2). The fold change values calculated by PyDESeq2 were normalized to make the average stability for non-regulated target sites in the main context equal to 1. Normalization was performed separately for each cell line. We initially divided all fold change values by the median of the 300 most stable constructs containing individual high confidence miRNA target sites in the main context, most of which are inactive. This shifts the overall distribution of DESeq2-calculated fold changes to be around 1. For all UTRs using the main context sequence (i.e., everything except the context controls), we then grouped designs by the number of miRNA target sites (1 to 6). Within each group, we divide fold change values by the median of the top 15% of most stable designs. This rescales fold change (stability) values to the mean observed for a certain number of inactive target sites and has a very minor effect for cell lines outside of MCF7 and HUH7. Designs using other context sequences are only processed using the first normalization step. Normalized stability data for Library 1 and 2 are provided as **Data S2** and **Data S4**.

##### The choice of miRNA expression data

We used two miRNA expression datasets for most analyses: microarray data by Alles et al. (*37*) and BGISEQ-500-based sequencing data generated for this study. Both datasets cover all used cell lines. Although the latter was generated for this study, it was collected in a different laboratory from the one where the reporter experiments were performed. We deliberately made no effort to harmonize cell lines or culture conditions to avoid biasing our results in favor of this newly collected data. Of note, both datasets contain expression data for SH-SY5Y, while the stability data was collected in SKNSH cells. Since SH-SY5Y were created from the subclone of SKNSH, we expect their miRNA expression profiles to be sufficiently similar for this to be valid.

##### Fitting a function for a single target site in the main context

For the analysis of single full target sites, we only use high confidence miRNAs in miRBase and low confidence miRNAs in miRBase that are in MirGeneDB, but not low confidence miRNAs in miRBase that are not in MirGeneDB. First, we normalized the miRNA expression data to transcripts per million (tpm) on this subset of miRNAs. We then used SciPy to fit our transfer function either for fixed total miRNA levels or variable total miRNA levels, i.e., we introduced an additional fitting parameter for each cell line that scales the total amount of miRNA in that cell line. For the latter, the total miRNA levels for HEK293T cells were kept constant to establish a reference point. We calculated deviation values for each miRNA target as the difference between the measured stabilities and the stabilities estimated by the transfer function on a logarithmic scale.

##### Single target sites in different context sequences

We used the fitted transfer function for target sites in the main context to predict the stability in all other contexts. Because the transfer function predicts relative stability changes due to the presence of miRNA target sites, measured stability values for miRNA targets $s_{miRNA, c}$ in different context sequences c were divided by the measured baseline stabilities $s_{0, c}$ of the surrounding context sequences without a target site in the same cell line: $s_{miRNA, rel} = s_{miRNA, c} / s_{0, c}$.

Estimation of the scale factor

We hypothesized that systematic global deviations from the transfer function are the result of both genuine biological variations in total miRNA levels between cell lines and differences in baseline context stabilities $k_{deg}$ between cell lines. Notably, Ago2 expression, which could serve as a proxy for the total slicing-competent miRNA level, is highest in PC3, where we do observe a stronger effect of miRNA (**Fig. S11A**). To estimate total miRNA levels, we therefore retrieved expression data (nTPM) for Ago proteins from the human protein atlas (*80*), which contains data for 8 of 10 of our cell lines. We then normalized these data to those in HEK293T cells. To estimate the relative stability of the main context, we took the geometric mean of 51 different measured context sequences and normalized this mean to the HEK293T value. For HUH7, stability of the main context relative to 51 other endogenously-derived context sequences is low, indicating that this effect might be dominant there (**Fig. S11B**). This approach assumes that the range of stability values covered by the different context sequences is similar across cell lines. The overall scale factor is then estimated by dividing the relative Ago2 expression by the relative mean context stability (**Fig. S11C**). We note that mathematically only a single scaling factor is necessary because the model only depends on the ratio of the miRNA-induced degradation and the baseline degradation (**Supplementary Text 1**).

Evaluation of the impact of secondary structure for single target sites

All $\Delta\Delta G$ values were calculated using the Python package for NUPACK version 4.0.0.20 (*73*) using the free_energy result of the complex_analysis function. When calculating these values, one must make a choice of how many bases to the 5' and 3' end of the miRNA target site to include. We used four different cutoffs (150, 100, 60, and 40 bases to the 5' and 3' end) and averaged the final results of the following procedure for increased robustness. We calculated the free energy of the miRNA $\Delta G_{miRNA}$), the target without context ($\Delta G_{target, ideal}$), the target with varying amounts of context ($\Delta G_{target, actual}$), and the miRNA in complex with the target both without ($\Delta G_{miRNA\text{-}target, ideal}$) and with context ($\Delta G_{miRNA\text{-}target, actual}$). We then subtracted the free energy of the miRNA and of the target from the free energy of their complex to calculate the binding $\Delta\Delta G$ between the miRNA and its target: $\Delta\Delta G_{ideal/actual} = \Delta G_{miRNA-target, ideal/actual} - \Delta G_{miRNA} - \Delta G_{target, ideal/actual}$. The $\Delta\Delta G$ difference value that we use in figures is then calculated by subtracting the binding $\Delta\Delta G$ without context from the binding $\Delta\Delta G$ with context: $\Delta\Delta G_{diff} = \Delta\Delta G_{actual} - \Delta\Delta G_{ideal}$. For individual target sites, $\Delta\Delta G$ differences larger than 10 kcal/mole were heuristically classified as indicating strong secondary structure. P-values for the deviation differences between strong and weak secondary structure were calculated using a one-sided Mann–Whitney U test. We constrained the analysis of deviation values to miRNAs expressed to a level larger than $10^{3.5}$ tpm, i.e., those where we expect a measurable impact on stability in the absence of secondary structure. In the analysis of designed secondary structures, we excluded the two constructs containing the strongest designed secondary structure for each miRNA target. These constructs contain contiguous 21 bp and 24 bp RNA duplexes, which cause a decrease in measured stability that we attribute either to a destabilizing effect of having a very strong hairpin in the 3'UTR or to the negative impact of such hairpins on library preparation, rather than an unexpected increase in miRNA activity at stronger secondary structures.

##### Processing and comparison of miRNA expression datasets

We used miRNA expression values provided by the authors either in the original publication or deposited on the GEO database. The details of the used datasets are available in **Table S3**. The first step of dataset processing depended heavily on the individual dataset and was performed with a custom python script. The authors' files were converted into a CSV file containing miRNA names, cell line identifiers, and non-normalized expression values. We used a custom R script and miRBaseConverter (*81*) to map older miRNA names to a MIMAT ID and to check whether the miRNA in question is still considered real in the current version of miRBase. Invalid miRNAs were discarded. We then converted the MIMAT ID back to the miRNA identifier in miRBase v22. Datasets were normalized to tpm, a baseline value of 1 was added to the expression level of each miRNA, then the dataset was normalized to tpm again and converted to log10 values. To compare datasets, we limited the miRNAs in each dataset to high confidence miRNAs in miRBase present in all compared datasets (242 miRNAs total). Microarray datasets were classified according to the manufacturer (Agilent, Affymetrix). Sequencing datasets were classified as either standard or improved based on their use of degenerate adapters to reduce ligation bias. We do not have sufficient cell line data to distinguish more subtle differences between datasets such as the use of UMIs. To calculate correlation values between datasets, correlations were averaged across all cell lines common to both datasets. To check the agreement with our stability dataset, the transfer function was fitted individually for each cell line. The resulting correlation values were averaged across cell lines for each dataset and then grouped by the dataset collection method.

##### Identification of false positives and negatives

For false positives, we first filtered miRNA targets for which we measured stabilities greater than $10^{-0.25}$ and expression levels larger than $10^{3.8}$ tpm. We then flagged those which display less activity than expected by the model (deviation > $10^{0.3}$) as potential false positives. For false negatives, we first filtered miRNA targets for which we measured stabilities lesser than $10^{-0.25}$ and expression levels smaller than $10^{3.5}$ tpm. We then identified those which display more activity than expected by the model (deviation > $10^{-0.3}$) as potential false negatives. We compared potential false positives and false negatives between the two expression datasets. When a miRNA expression measurement was identified as a potential false positive/negative in one dataset but not the other, it was retained as a potential false positive/negative. If a miRNA expression measurement was identified as a potential false positive/negative in both datasets, we concluded that it is not a genuine false positive/negative. We labeled miRNAs that appear as a potential false positive or negative in at least 4 cell lines after this filtering step as genuine false positives or negatives. **Table S7** contains a list of these identified false positives and negatives.

##### Analysis of mutated target sites and generation of a crosstalk model

Addressing crosstalk serves to increase the predictive accuracy of our model. In principle, one could simply filter all but the most highly expressed miRNA that share a core seed sequence (nts 2 to 7). However, this leads to a strong reduction in the available number of potential miRNA target sites. This problem gets worse as the number of target cell types increases because the highest expressed miRNAs of a given family can differ between tissues. Thus, crosstalk filtering is a balance between increasing predictive accuracy and retention of as many potential target sites as possible. We note that simply adding expression levels of the same family is often worse than not addressing crosstalk at all and not a viable option for quantitative prediction.

The measured stabilities s were converted to relative knockdown values, which we define as $\frac{s_{full\ target}}{1-s_{full\ target}}\frac{1-s_{mut\ target}}{s_{mut\ target}}$. This expression yields a value of 1 when the mutated target site is as strong as the full target site and 0 when the mutated target site is completely inactive. Analyses were generally performed on miRNA target sites where the stability value for the full target site is less than ⅓ in a given cell line to ensure that this value can be accurately determined. We determined the distribution of these relative knockdown values depending on where in the sequence mutations and wobbles occur ignoring the specific miRNA sequence and identity. We then classify individual mutations and wobbles according to the median relative knockdown at a given position with values less than 0.08, 0.3, and 0.65, or greater than 0.65 being classified as high, medium, low, and no impact mutations and wobbles, respectively. This classification was performed independently for true mismatches and for wobbles. The classification thresholds for the median were chosen heuristically such that each category contains multiple positions (10, 12, 16, and 4 for high, medium, low, and no impact mutations). When the classification for a wobble base pair at a specific location was stronger than for the corresponding mismatch, the classification for the wobble was lowered to the level of the mismatch. P-values between the mismatch and wobble relative knockdown distributions were calculated using a two-sided Mann-Whitney U test.

To train a tree model, we applied the previously derived mutation classification to targets with multiple mutations and counted how often each mutation type occurs. The mutation counts by mutation strength were the input values for the tree model. Training was performed on relative knockdown values constrained to mutated target sites where the non-mutated target site had a stability value less than ⅓. We held out all mutated hsa-miR-31-5p target sites as test data. We trained both an XGBoost model (xgboost version 1.7.3) and a DecisionTreeRegressor (sklearn version 1.3.0) and found nearly identical performance. The decision tree predicts relative knockdown values, which are converted to stabilities afterwards. The predictions shown **Fig. S17C** were performed on all measured mutated target sites irrespective of the stability of the non-mutated target.

Because the impact of mutations turned out to be highly sequence-specific, we decided to largely filter miRNA target sites where significant crosstalk is expected instead of trying to quantitatively adjust predictions for target sites for which crosstalk is expected: First, we added the expression levels for miRNAs that are identical for the first 18 bases, which we expect to cause nearly identical behavior (**Table S8**). We note that this is not necessarily fully accurate as mutations after base 18 can still have an impact (**Fig. S16C-E**). We nevertheless consider it preferable to filtering these target sites. We then identified miRNAs with potential crosstalk as all pairs of miRNAs and miRNA targets separated by fewer than 2 high impact mutations, 4 high and medium impact mutations, and 5 total mutations (**Table S9**). This is based on the maximum number of mutations of a given impact that still allows for crosstalk (**Fig. S16E**). These sites with potential crosstalk were filtered if the stability predicted by the transfer function for the full target site of the crosstalking miRNA is smaller than $10^{-0.5}$ and less than ⅓ of the predicted stability of the fully complementary miRNA for the target site in question.

<u>Analysis of seed-pairing stability and transcriptomic target abundance</u>

To calculate seed-pairing stability (SPS), we took bases 2 to 8 of each miRNA and calculated the binding energy with its reverse complement using the free_energy result of NUPACK's complex_analysis function. To calculate the transcriptomic target abundance we again took bases 2 to 8 of each miRNA. The human reference transcriptome was obtained from the NCBI RefSeq database (accession GCF_000001405.40, assembly GRCh38.p14) in GenBank format (rna.gbff file), which contains annotated transcript and gene sequences for the GRCh38.p14 build. We calculated seed abundances in the human genome either for the entire mRNA or for just the 3'UTR as the sequence between the end of the annotated CDS and the start of the poly(A) site. The relevant figures show the result for using just the 3'UTR but the results are not sensitive to this choice. To account for transcript abundance by cell line, we obtained expression values (nTPM) for each mRNA for a subset of our cell lines from the human protein atlas (*80*). Expression-adjusted seed counts were obtained by a weighted sum of mRNA expression values and seed counts per mRNA.

<u>Repeat and combination data</u>

We used the bias-corrected expression data demonstrated in **Fig. 1H**. We predicted the expected stability using the transfer function and scale factors derived for single target sites and using either the additive model or an antagonistic model where only the miRNA target site with the highest associated expression level is used. For the inverted transfer function, expression levels for a given microRNA in a given cell line are calculated from stabilities $s$ via $t^{-1}(s) = \frac{k_{deg}}{k_{on}} \frac{1-s}{s}$. Since the transfer function only outputs values between 0 and 1, stability values larger than 1 were reduced to 0.999 for calculations using the inverted transfer function. We then input these calculated expression levels into the transfer function using the additive model.

<u>Analysis of outliers in the repeat and combination data</u>

Unusual patterns in the repeat data were identified as miRNAs for which two and four repeats were at least a factor of $10^{0.1}$ more stable than one and three repeats, or three and five repeats were at least a factor of $10^{0.1}$ more stable than two and four repeats. Constructs with a stability value of at least 1.5 were identified as highly stable. This yielded 13, 3, and 14 miRNAs where these conditions were fulfilled in at least one cell line.

To identify the origin of the unusual patterns in the repeat data, we calculated pairwise interaction strengths between the individual target sites as the free energy difference between a complex of two targets and the individual targets. To more generally detect whether secondary structure plays a role for any construct with multiple target sites, ΔΔG difference values between miRNAs and 3'UTRs with multiple target sites were calculated as for individual target sites but with 150 or 100 nts on either side of the center of the variable 3'UTR region. Here, we heuristically classified sites with ΔΔG differences larger than 14 kcal/mole as strong secondary structure sites based on the pattern observed in **Fig. S8**. Constructs were identified as likely subject to prediction deviations due to secondary structure when the site with the largest cognate miRNA expression value was classified as having strong secondary structure.

Performance prediction and evaluation of constructs with designed stabilities

We compare three prediction methods: First, the baseline prediction based purely on microarray data (*37*) and the transfer function fitted using the measurements from Library 1. Second, an updated model based on bias-aware merging of microarray and improved sequencing data as demonstrated in Figure 1 of this manuscript. The updated model also applies the scale factors derived from individual target sites. Third, predictions based on estimating expression levels via the inverted transfer function as described above for repeat and combination data. All predictions use the additive model. We filtered designs where the measured stability in any cell line is larger than 1.5 as this makes it difficult to estimate true relative stabilities (15 out of 3753 total designs). To evaluate design performance, stabilities larger than 1 were set to 1. When choosing the best-performing designs or when ranking design success into quantiles, we used the weighted mean square error as explained above in the design section. When comparing the design success across design types, we used the unweighted root mean square error between the target and measured stabilities.

**Processing and merging of human tissue miRNA expression datasets**

We used an Agilent microarray dataset from two human subjects by Ludwig et al. (*62*) and a BGISEQ-500-based sequencing dataset from six human subjects by Keller et al. (*63*). The data collection methods match the two main datasets used for the rest of this study, making it likely that biases behave at least similarly. We first filtered to miRNAs that are either high confidence in miRBase or in MirGeneDB. Data were then normalized to tpm. To calculate correlation values between subjects for the same tissue and dataset, we set expression values less than 100 tpm to 100 tpm. To integrate data from the different subjects and produce a single expression dataset across tissues, we then calculated the geometric mean of expression values for each tissue across all subjects and again normalized to tpm (**Fig. S38**). To compare the two datasets, we first harmonized tissue names and again set values less than 100 tpm to 100 tpm. Tissues that were not in both datasets were discarded. To calculate brain expression values for the NGS dataset, we averaged over temporal lobe, occipital lobe, frontal lobe, white matter, and gray matter values. We found that there are many miRNAs that behave differently in the two datasets. To identify these outlier miRNAs, we first renormalized the expression values because the microarray dataset has more highly expressed miRNAs overall. Regular normalization to tpm therefore leads to an underestimation of individual expression levels relative to the NGS data. We determined the top 30 most different miRNA by expression in each tissue. We identified miRNAs that are in the top 30 in at least three tissues, then renormalized the dataset to tpm while excluding them from the calculation. We again compared miRNA expression levels across tissues and datasets. We then determined outliers in a specific tissue as those miRNAs where the renormalized expression values differ by more than a factor of 10. miRNAs which are consistent outliers in 5 or more tissues were classified as consistently larger in NGS (5 total) or microarray (94 total) data. We then calculated Pearson correlation values with and without these outliers on a log expression scale (**Fig. S39A-B**).

To merge the two datasets, we asked whether it is possible to guess which of the two datasets is correct for which outlier. First, we calculated deviation values in the cell line stability for each outlier miRNA for both our microarray and NGS data cell line expression data (**Fig. S40A**). We excluded cell lines (JEG3, Tera1) with poorer prediction performance from the analysis. We subtracted absolute deviation values for the two cell line datasets and considered a difference greater than $10^{0.2}$ as signifying substantially more correct prediction by one or the other method (**Fig. S40B-D**). Second, we added false positive and negative information from our cell line data. Third, we inspected the sequence features of outliers. We noticed that many of the strongest outliers had a G-content greater than 50% (**Fig. S40E-F**). An earlier publication (*82*) found similar outliers between microarray and NGS data. There, RT-qPCR agreed with the NGS data over the microarray data. We therefore also labeled miRNAs with a G content of over 50% that were previously identified as consistent outliers as more likely to be correctly measured by improved NGS data. In total, we classified 12 miRNAs as incorrect in the NGS data and 16 miRNAs as incorrect in the microarray data. We then performed a bias-aware merging based on the geometric mean as for the cell line data (**Fig. S40G**). The resulting data was then further merged and filtered for crosstalk as for the cell line data.

**<u>Designs with tailored stability profiles in human tissues</u>**

We used the merged and crosstalk-filtered human tissue expression dataset. Because this dataset contains fewer miRNAs than the merged cell line data due to fewer input miRNAs in the microarray tissue data, we performed another round of fitting to the cell line stability data for this subset of miRNAs. The design process was generally performed as for cell lines with a few differences. First, we only generated a single design per design target and miRNA target number (1 to 8). Second, we filtered the expression dataset to miRNAs with a maximum expression greater than 3000 tpm across target tissues, which reduces computational complexity and focuses the algorithm on potentially relevant miRNAs. Third, we introduced an 'empty' target site with zero expression in all tissues, which allows the algorithm to use fewer than the maximum number of target sites. We generated designs that are either active or inactive in a single tissue. The stability in that tissue was given a weight of 6.5 in calculating the mse. To compare the design quality, defined as the inverse of the weighted mse between the prediction and the target stability profile, between different target site numbers, we divided all quality values for designs targeting the same tissue by the maximum quality across the allowed target site numbers. Design for up to six target sites are provided in **Table S10**.

**<u>Ratiometric stability data analysis (Library 3)</u>**

Analysis of ratiometric stability data for Library 3 largely followed the same steps as for Library 2, where applicable. When fitting the transfer function with two constants, we initially fitted the first constant excluding the saturation term, then determined the saturation term. We identified four microRNAs that were completely inactive in 3T3 and P2C2 cells (mmu-miR-378a-3p, mmu-miR-191-5p, mmu-miR-181b-5p, mmu-miR-181a-5p) that were consequently excluded from the analysis in mouse CD8 T cells due to likely bias. For comparing the transfer function to

HEK293T data, we subsetted the analysis to mouse miRNAs with near identical human miRNAs (identical for the first 19 bases).

To find consistent outliers in the repeat data for non-expressed microRNAs, we subset to miRNAs with $<10^{2.5}$ tpm, then counted how often each five target repeat has a stability $>10^{0.3}$ or $<10^{-0.3}$ across T cell conditions (six total, two for E1 and four for E2). We highlight cases where the stability crosses the threshold for high stability at least two times or the threshold for low stability at least three times.

To determine the microRNAs that best distinguish T cells from mouse cell lines (3T3, P2C2), we subset to designs with a predicted stability of at least 0.66 for a given repeat number in the target cell lines, then sorted by the margin between on- and off-target cells (minimum in the on-target cells minus maximum in the off-target cells).

#### Time-course stability data analysis

We removed all constructs with less than 20 UMIs at time 0, retaining 10007 out of 10028 total constructs. We then normalized counts using the spike-ins by fitting a linear model y=a*x+b, where y is the observed sequencing UMIs and x is the expected spike-in abundance. UMIs for each time point t were then scaled by the ratio $a_0/a_t$. We truncated the time series for each construct at the point where the relative abundance compared to time 0 fell below 0.1. This is necessary to deal with very low UMI counts at later time points for low stability constructs. We then fit an exponential decay function $m_0*\exp(-k*t)$ to all constructs and calculated half-lives as ln(2)/k. We filtered any construct where the Pearson $r^2$ between the log10 of fitted and measured values was below 0.85 (146 constructs total).

#### Polysome profiling data analysis

We removed all constructs with less than 30 UMIs at time 0, filtering out 4 constructs for HEK293T and 51 for HeLa. Spike-in UMI counts were divided by the number of independent tubes contributing to each polysome fraction, correcting for the amount of spike-in added per fraction. We then normalized counts using the spike-ins by fitting a linear model y=a*x+b, where y is the observed sequencing UMIs and x is the expected spike-in abundance. UMIs for each time point t were then scaled by the ratio $a_0/a_t$. To calculate the mean ribosome load (MRL) for a particular construct, we performed a weighted sum over spike-in adjusted UMI counts $c$ and observed ribosome counts $r$ per fraction: $MRL = \frac{\sum_{i=0}^{n} r_i c_i}{\sum_{i=0}^{n} c_i}$. The number of ribosomes per fraction is estimated based on the absorbance curves (**Fig. S27A-B**). The 0 ribosome fraction is included in this calculation.

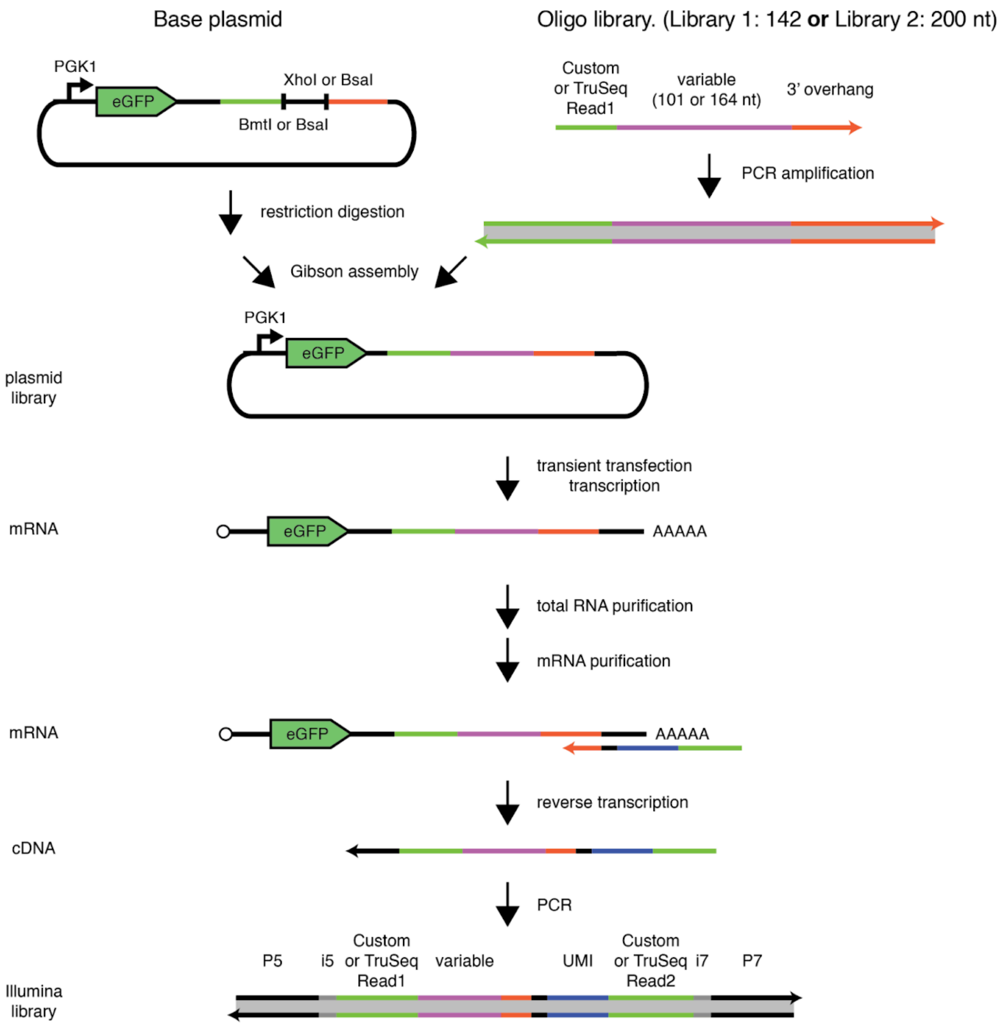


**Fig. S1. Library construction and purification scheme.** Assembly and processing of Library 1 and Library 2. Differences between the workflow for the two libraries are marked by an "or" (Library 1 or Library 2). Library oligo pools are amplified via PCR to introduce overhangs. The base cloning plasmid is digested with restriction enzymes and the amplified oligo pool is inserted via Gibson assembly to generate plasmid libraries. After transient transfection into the target cell lines, mRNAs transcribed from the library plasmids are first purified as total RNA, then with a poly(A)-based mRNA isolation kit. Reverse transcription is performed with a primer containing a unique molecular identifier (UMI). The cDNA is amplified with i5 and i7 index primers and sequenced. The primary differences between the library preparation for the two libraries are the restriction sites used for digestion of the base plasmid, the length of the variable region, and the use of custom read 1 and read 2 sequencing primers for Library 1.

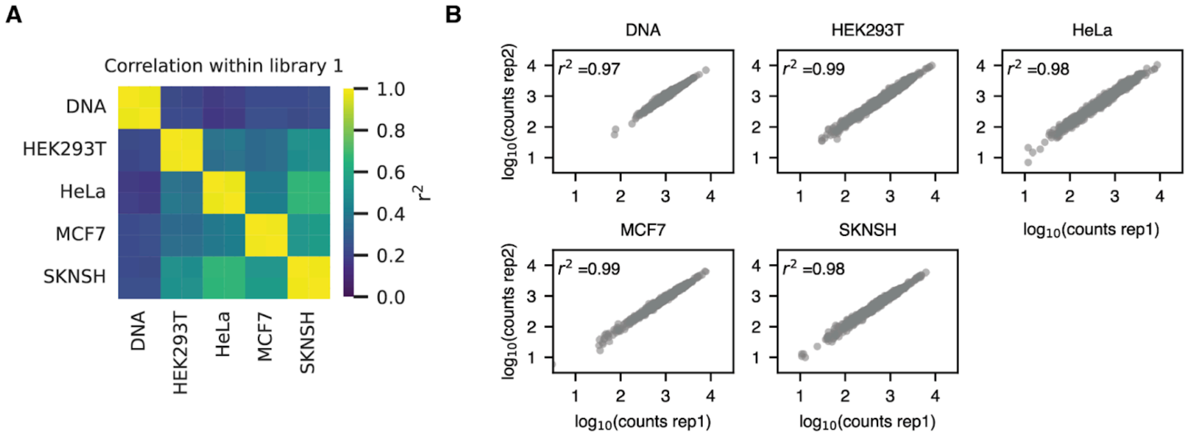


**Fig. S2. Correlation between Library 1 replicates.** (**A**) Correlation between counts across replicates and cell lines for Library 1. (**B**) Comparison of counts for individual designs between the two library replicates for different cell lines.

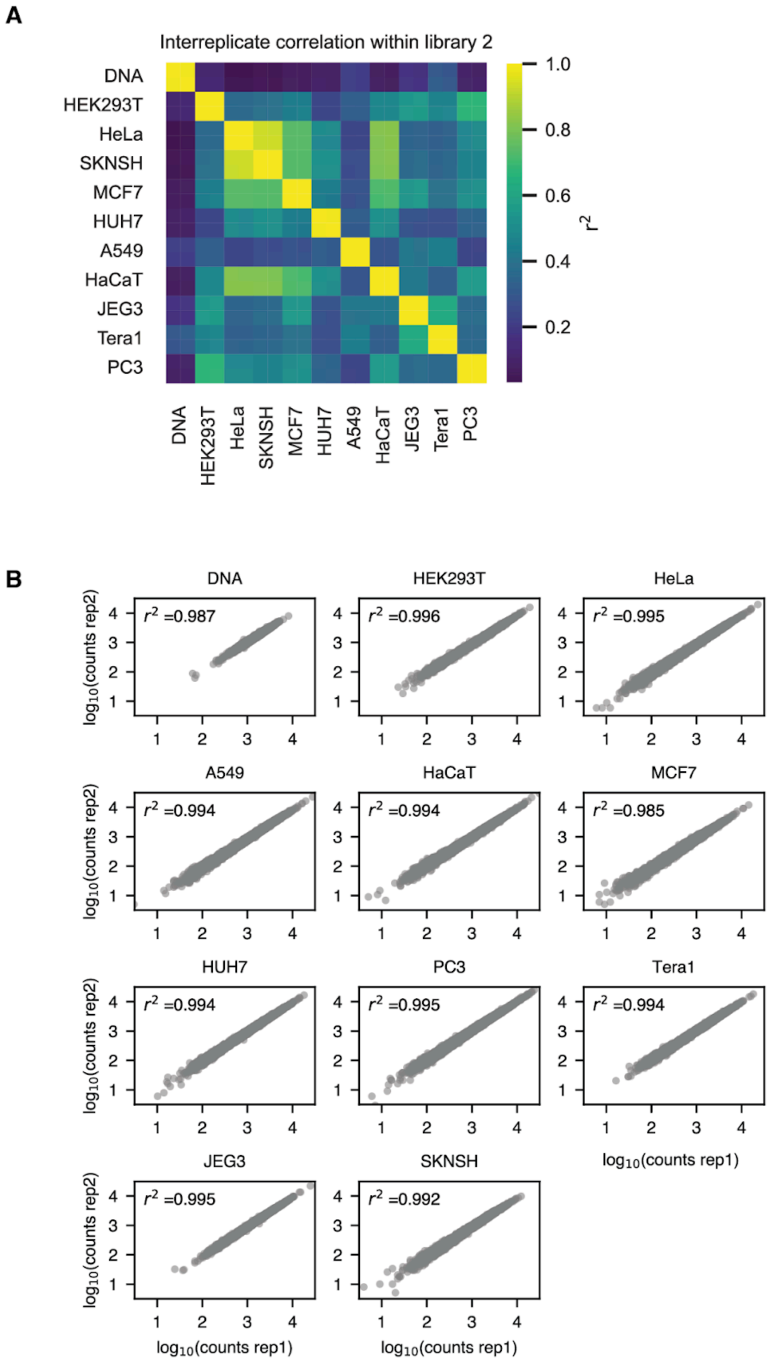


**Fig. S3. Correlation between Library 2 replicates.** (**A**) Correlation between counts across replicates and cell lines for Library 2. (**B**) Comparison of counts for individual designs between the two library replicates for different cell lines.

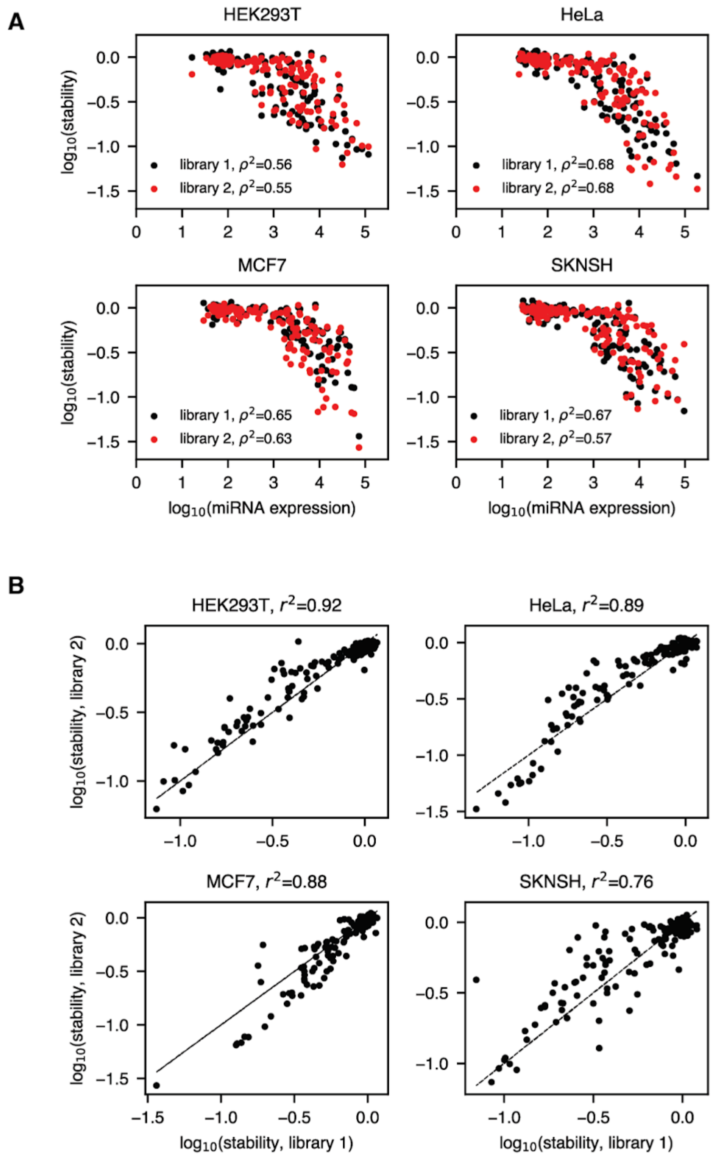


**Fig. S4. Comparison of the results for Library 1 and Library 2. (A)** Microarray-measured miRNA expression (*37*) versus the measured stability for miRNA targets and cell lines shared between Library 1 and 2. The distributions largely agree. The legend shows the Spearman $\rho^2$ between expression and stability. (**B**) The measured stabilities for both libraries. Note that the main context sequences are different for the two libraries. The relatively large variation for SKNSH could imply biological differences in the cell line miRNA concentrations at the time of measurement for the two libraries.

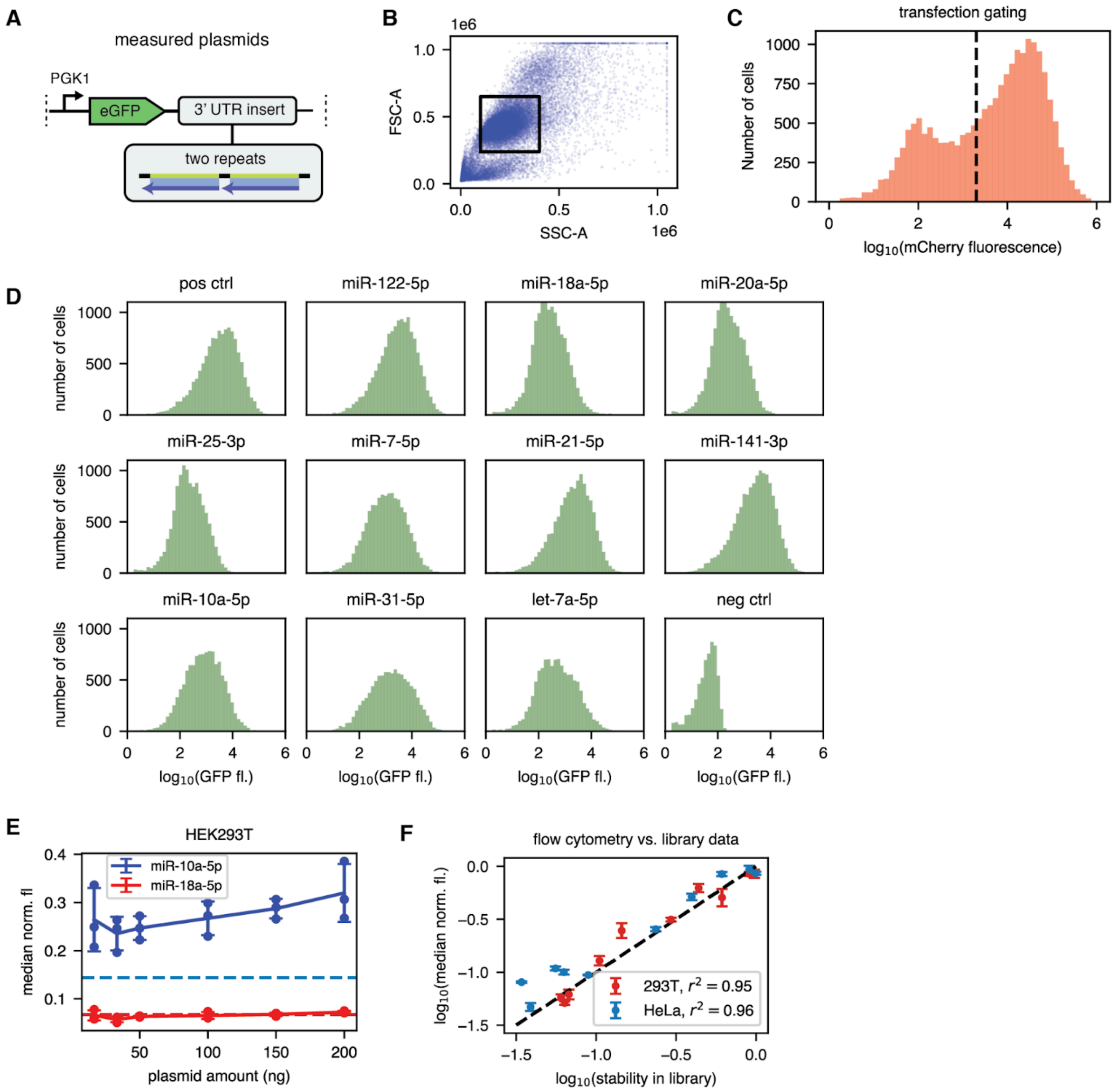


**Fig. S5. Flow cytometry results for individual constructs from library 1.** (**A**) We measured constructs containing two repeats of a microRNA target site. The sequence is identical to the same constructs in Library 1. (**B**) Gating strategy on SSC-A and FSC-A for HEK293T cells. (**C**) The gating for transfected cells using an mCherry-expressing transfection control. (**D**) GFP fluorescence distribution for one replicate in HEK293T after filtering for transfection using the mCherry signal. The negative control is shown without mCherry gating. (**E**) Median fluorescence normalized to the median fluorescence of the positive control for two different microRNA target types versus the transfected plasmid concentration. The result is largely independent of plasmid concentrations when using the median fluorescence. N=3 (**F**) Relationship between the stability measured for Library 1 for our high-throughput stability data and the median GFP fluorescence level normalized to the median fluorescence of the positive control measured using flow cytometry for two different cell lines (HEK293T, HeLa). N=3

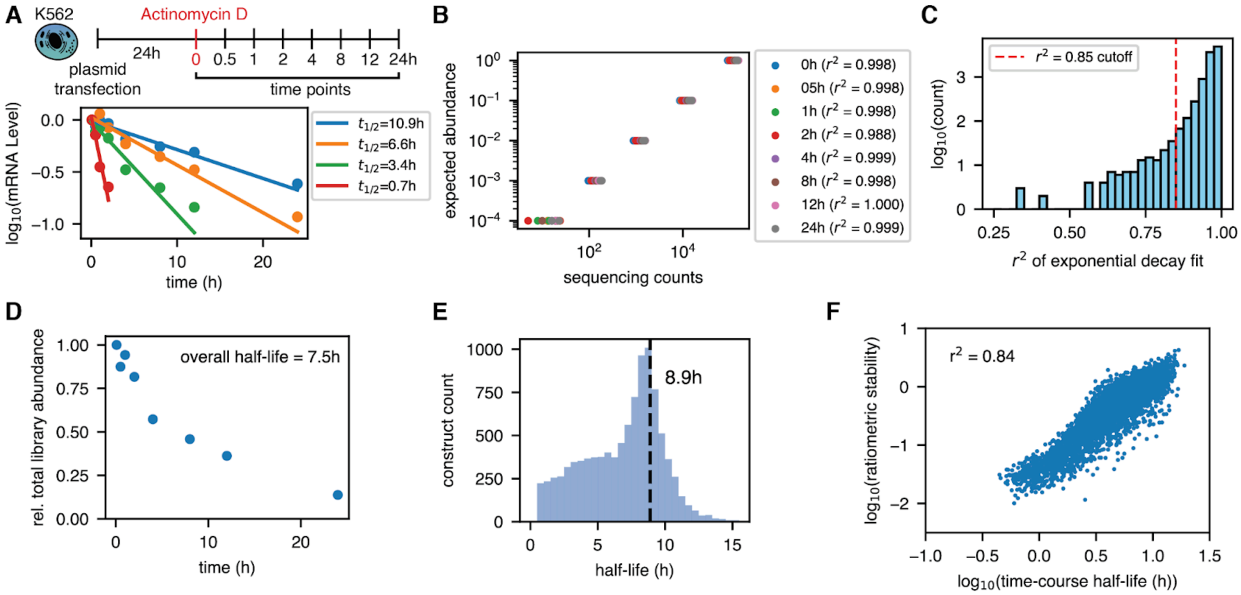


**Fig. S6. Stability time course measurements.** (**A**) Top: Schematic of the experiment. Bottom: Examples of exponential decay fits for four different constructs. (**B**) Expected abundance and sequencing counts for spike-ins used to normalize the counts for different time points. (**C**) Distribution of the Pearson $r^2$ of exponential decay model fits. Constructs with values below the thresholds were excluded from the analysis. (**D**) Overall half-life averaged across all library members. (**E**) Distribution of measured half-lives in the library. The most common half-life corresponding to a stability of 1 in the ratiometric assay, is 8.9h. (**F**) Correlation of measured half-life in the time-course based measurement.

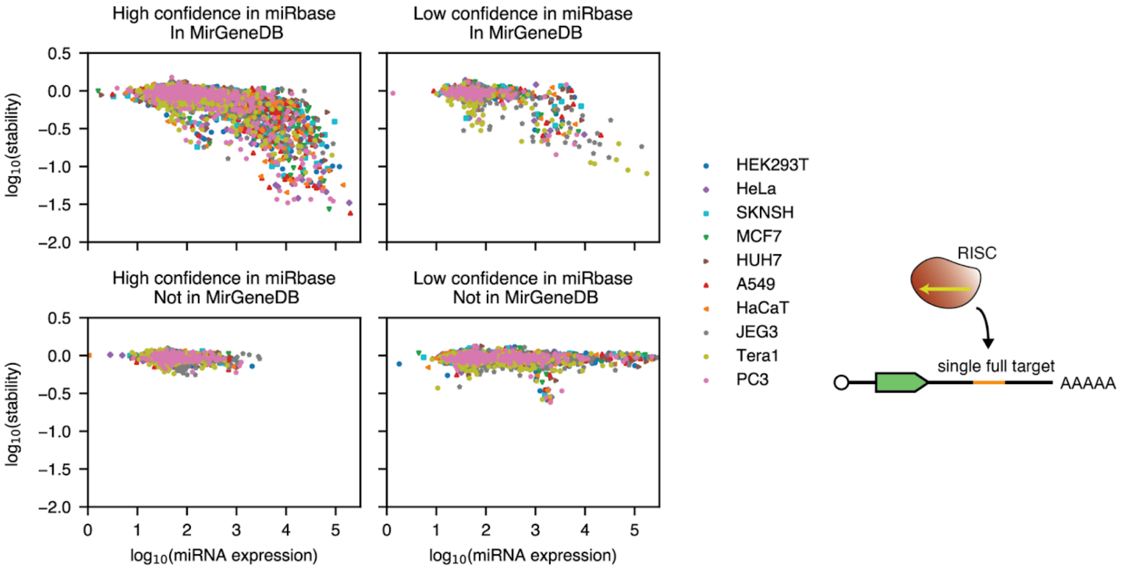


**Fig. S7. The impact of the microRNA annotation data source on the observed behavior for single target sites.** The microRNA expression data is from Alles et al. (*37*). MicroRNA targets in miRBase are annotated as high confidence or low confidence depending on the available level of experimental evidence for their existence. Later additions to miRBase are often low confidence sequences derived purely from sequencing data without any direct biological evidence. MirGeneDB is a manually curated microRNA database that more closely considers clear experimental evidence. Most, though not all, microRNAs that are in MirGeneDB are annotated as high confidence in miRBase. We find that the association between miRNA expression and target stability depends strongly on the annotation data source: All microRNAs in MirGeneDB approximately follow a monotonous relationship regardless of their status in miRBase, validating them as likely real microRNA. The microRNA that are annotated as high confidence in miRBase but not in MirGeneDB have insufficient expression in our measured cell lines to draw definitive conclusions. Low confidence microRNA that are not in MirGeneDB generally show no activity whatsoever even when (erroneously) measured as highly expressed. We therefore exclude them from the analysis.

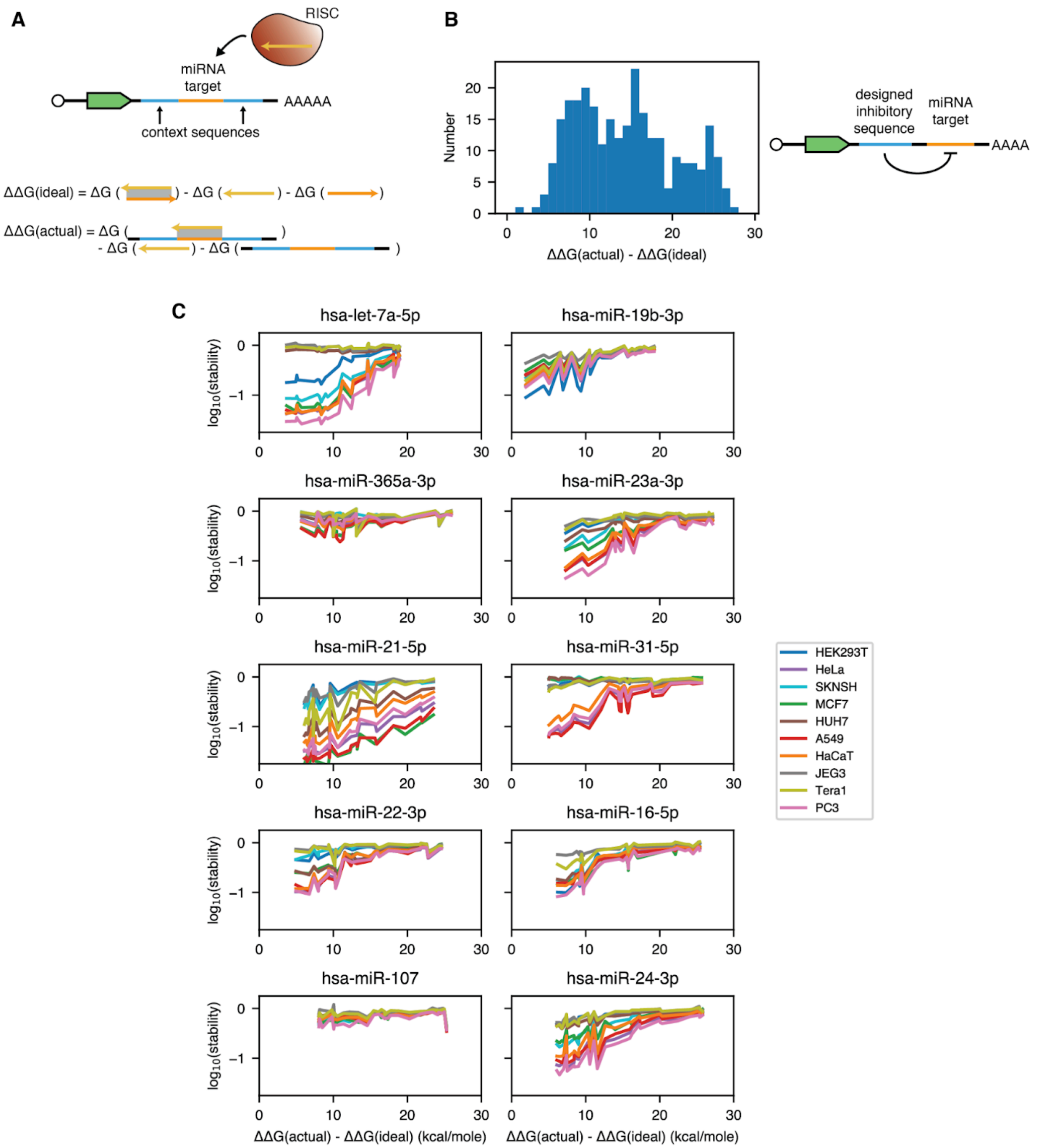


**Fig. S8. Strong target secondary structure inhibits microRNA function.** (**A**) We calculate the binding energy (ΔΔG) between the miRNA and its target without and with the surrounding context sequence (*73*). The ΔΔG value shown on the x-axis for all graphs investigating secondary structure in this publication is the difference between these two values. (**B**) Our engineered context sequences display a wide distribution of ΔΔG differences, allowing us to investigate the dependence of miRNA activity on target secondary structure. (**C**) The measured stabilities of identical miRNA targets depend on the ΔΔG difference between ideal and actual binding energies. The effect is relatively consistent across cell lines and microRNAs. A single connected line in this graph belongs to a single microRNA target site occluded by different amounts of secondary structure in a single cell line.

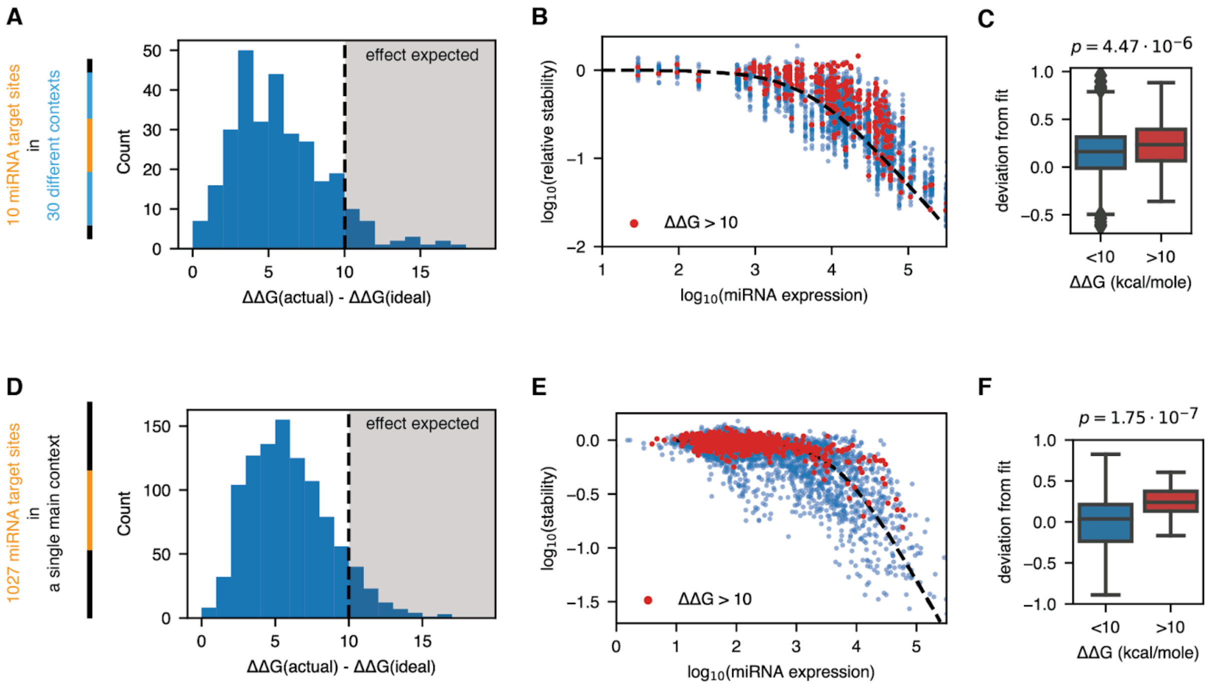


**Fig. S9. The impact of secondary structure is generally small for natural context sequences.** (**A**)-(**C**): Results for 10 miRNA target sites in 30 context sequences. (**D**)-(**F**): Results for 1027 miRNA target sites in the main context sequence. (**A**)/(**D**) Distribution of the binding energy differences. (**B**)/(**E**) MicroRNA expression and target transcript stabilities for low and high (>10 kcal/mole) binding energy differences. (**C**)/(**F**) Distribution of the fit deviation values for low and high binding energy differences. We constrained the calculation to miRNAs with an expression greater than $10^{3.5}$ tpm to exclude miRNA targets with no expected effect on stability. The p-value was calculated using a one-sided Mann-Whitney U test.

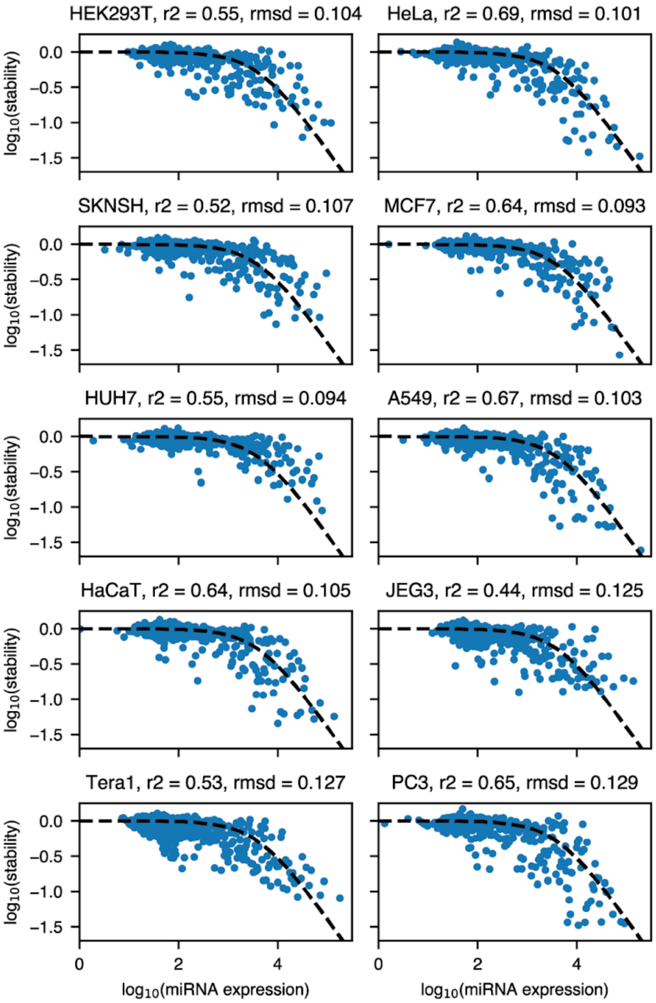


**Fig. S10. Fit results for individual cell lines and normalized microRNA expression data.** The x-axis contains microarray microRNA expression data (*37*). We fit a single universal transfer function for all cell lines. The titles show the Pearson $r^2$ values and the root mean square deviation (rmsd) between the measurements and the transfer function fit.

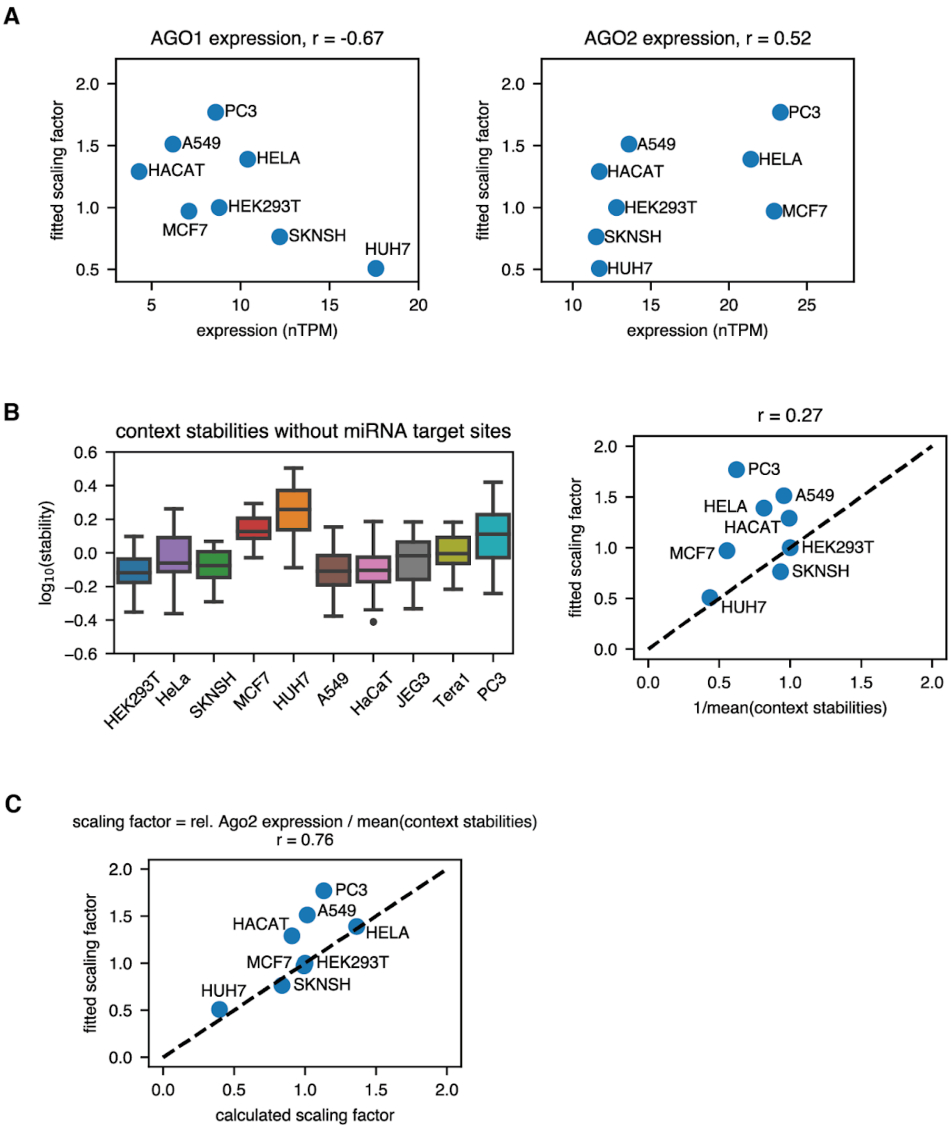


**Fig. S11. Ago2 expression and context stability explain the scaling factor.** (**A**) Ago1 and Ago2 expression according to protein atlas data versus the fitted scaling factor for individual target sites. Ago1 expression is negatively and Ago2 expression is positively correlated with the scaling factor. We used Ago2 expression as a proxy for the total amount of slicing-competent miRNA in a cell. (**B**) Left: The distribution of baseline stabilities for 51 different 3'UTRs (Methods) without miRNA target sites. Because the normalization is performed relative to the stability of the main context, the shown stabilities are also relative to the stability of the main context sequence. Right: The inverse of the geometric mean of the stabilities shown on the left versus the fitted scaling factor. This value approximates the relative stability of the main context across cell lines. (**C**) Dividing the relative Ago2 expression normalized by the HEK293T value by the geometric mean of the context stabilities yields a good estimate of the overall scaling factor.

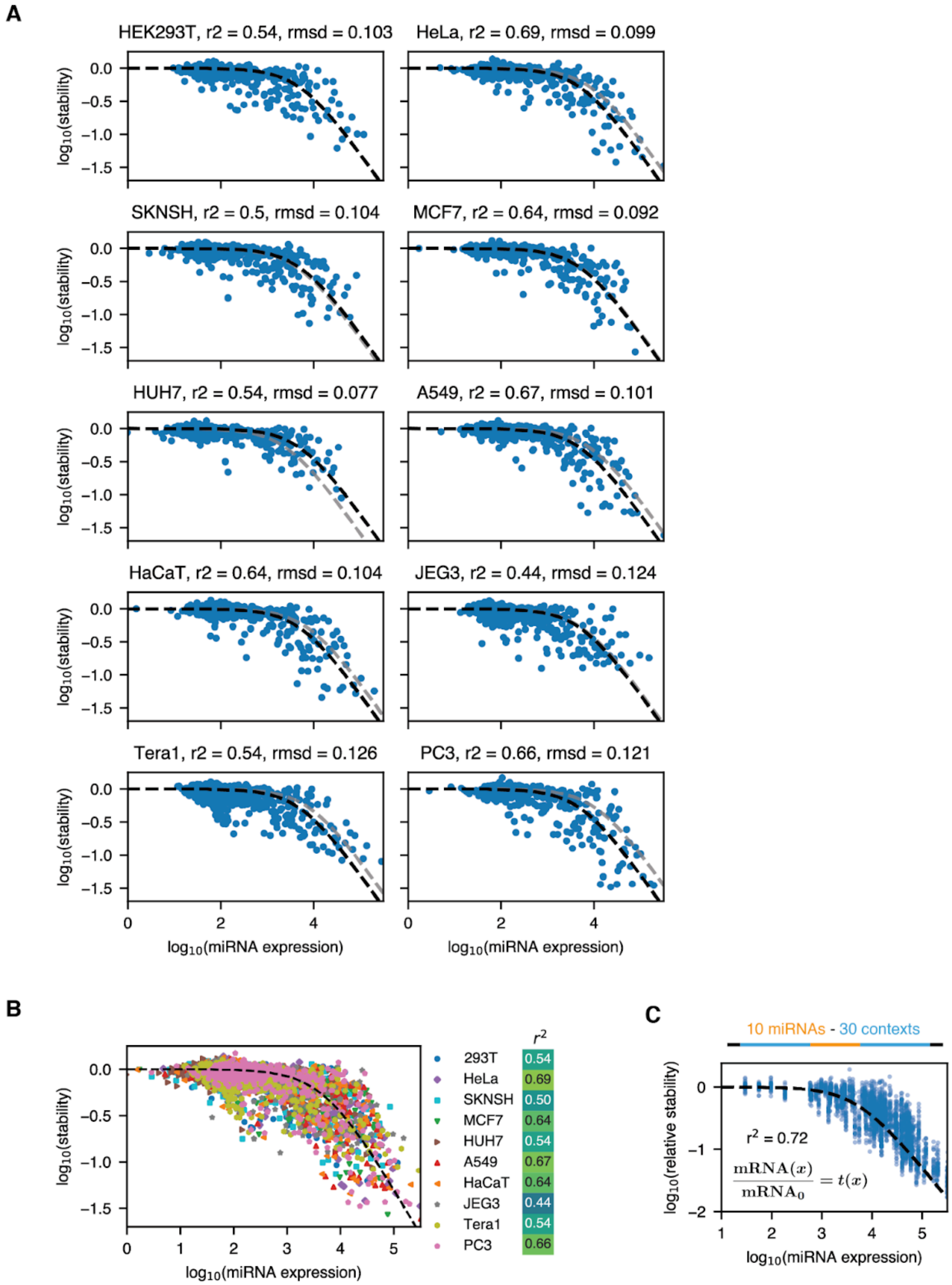


**Fig. S12. A scaling factor for the total microRNA expression levels improves the model fit.** (**A**)/(**B**) Scaled microRNA expression versus stability for target sites in the main context sequence. The titles and the heatmap show the Pearson $r^2$ values and the root mean square deviation (rmsd) between the measurements and the transfer function fit. The adjusted transfer function is shown in black and the unscaled transfer function in gray. (**C**) The scaling factor also improves the predictions of the relative stability of 3'UTRs containing miRNA targets in different context sequences.

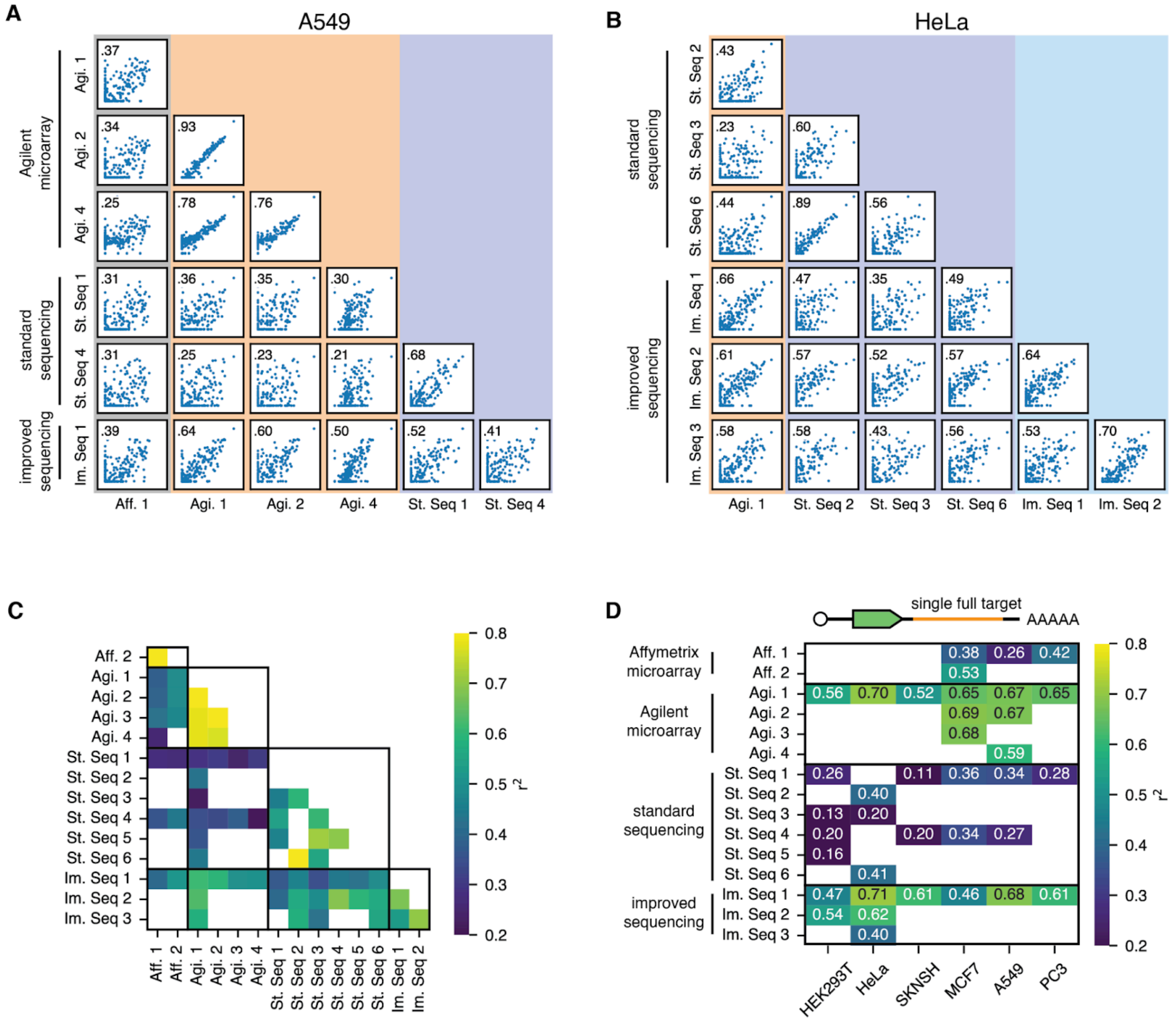


**Fig. S13. Cross-dataset correlation predicts well-performing datasets.** Cross-dataset correlation for (**A**) A549 and (**B**) HeLa cells. All values are set to a minimum of 100 tpm and the analysis is constrained to miRNAs common to all datasets. (**C**) Cross-dataset Pearson $r^2$ averaged across all cell lines. White squares indicate a lack of overlapping cell lines between two publications. (**D**) Pearson $r^2$ between the fitted transfer function using different microRNA expression datasets and measured stability for single microRNA target sites. White squares indicate that a cell line is missing for a given cell line. Well-performing datasets have both high correlation with other datasets within their collection method and also with some other collection methods.

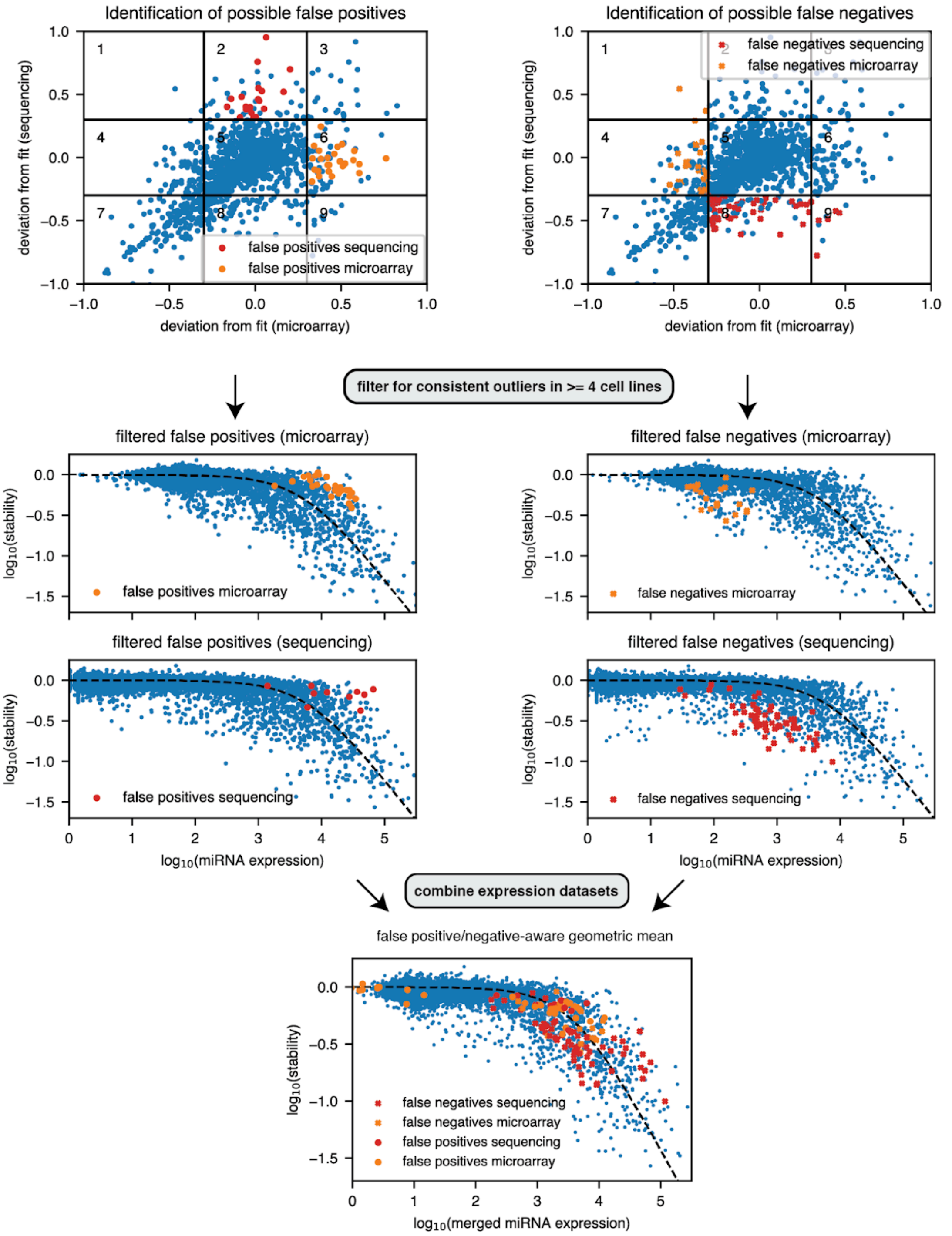


**Fig. S14. A comparison of datasets identifies biased data points.** We compare the transfer function fit for a microarray (*37*) dataset and the sequencing dataset collected for this study for all cell lines. First, we identified potential false positives and negatives in both datasets as outliers from the fit in only one of the two datasets (top plots, see Methods for details). Then, these potential outliers were filtered for agreement across our cell lines: An outlier in a single cell line could represent measurement noise or genuine biological differences in the microRNA expression at measurement time. An outlier miRNA that is consistent across multiple cell lines is likely to be due to technical differences in the expression measurement method. The middle plots highlight miRNAs that were identified as false positives and negatives in the two datasets in the expression versus stability plot. After merging, the previous false positives and negatives were predicted well by the transfer function, indicating that they were indeed incorrectly measured in the other dataset (bottom plot).

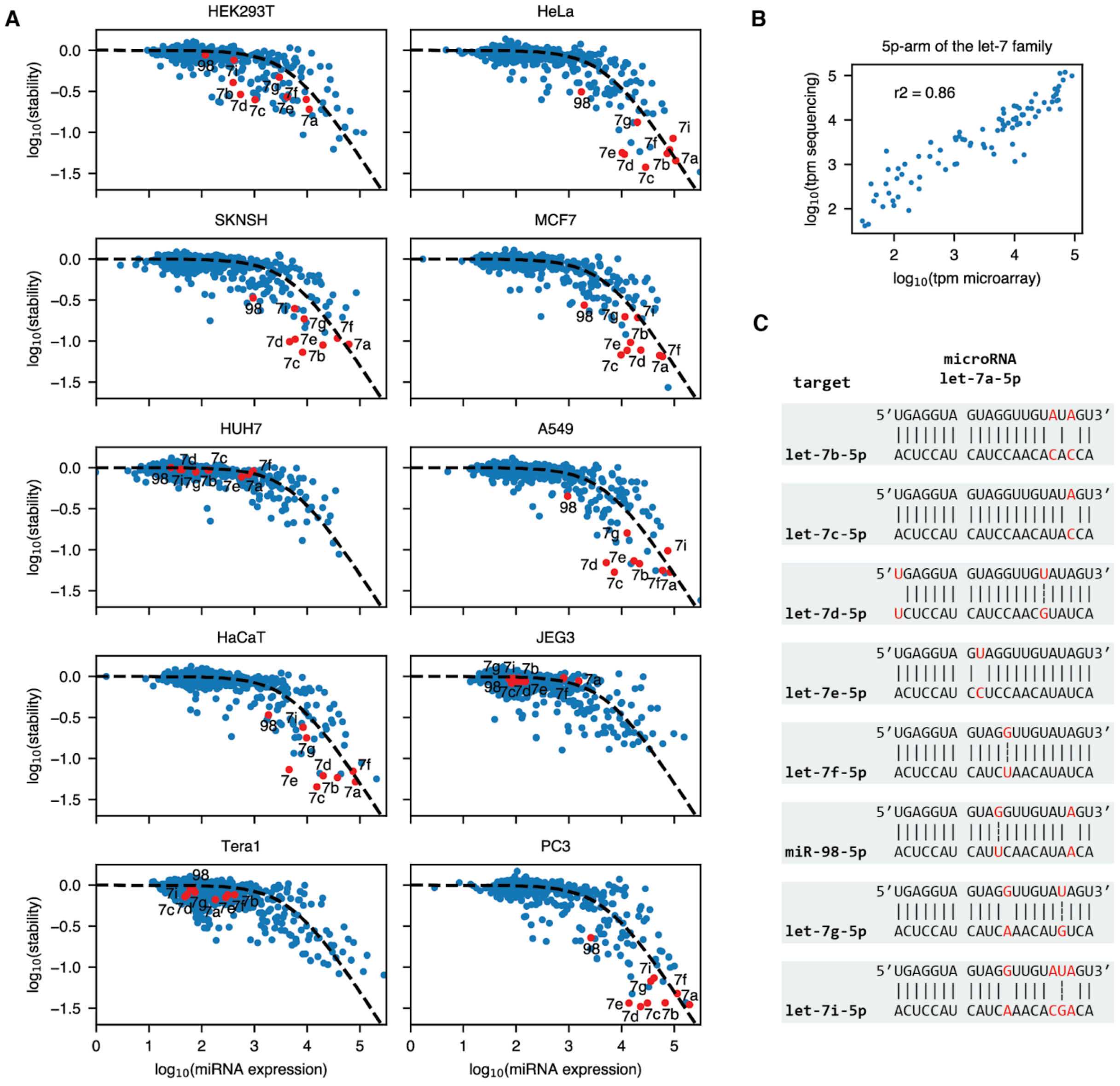


**Fig. S15. Crosstalk between let-7 family members causes outliers from the fit.** (**A**) In the stability versus expression plot, targets of the 5p arm of the let-7 family consistently appear as some of the strongest outliers with less stability than would be expected from the transfer function. The labeled miRNA targets are abbreviated from the full hsa-miR/let-x-5p name. (**B**) The let-7 family member expression is consistent between the microarray and sequencing measurements, indicating that the observed crosstalk is not a result of bias in the expression measurements themselves. (**C**) The mismatch pattern between an attacking hsa-let-7a-5p miRNA and target sites belonging to other let-7 family members generally only includes one or two mismatches (red, gap without dash) or wobble base pairs (red, dashed line), which explains qualitatively why such strong crosstalk is observed.

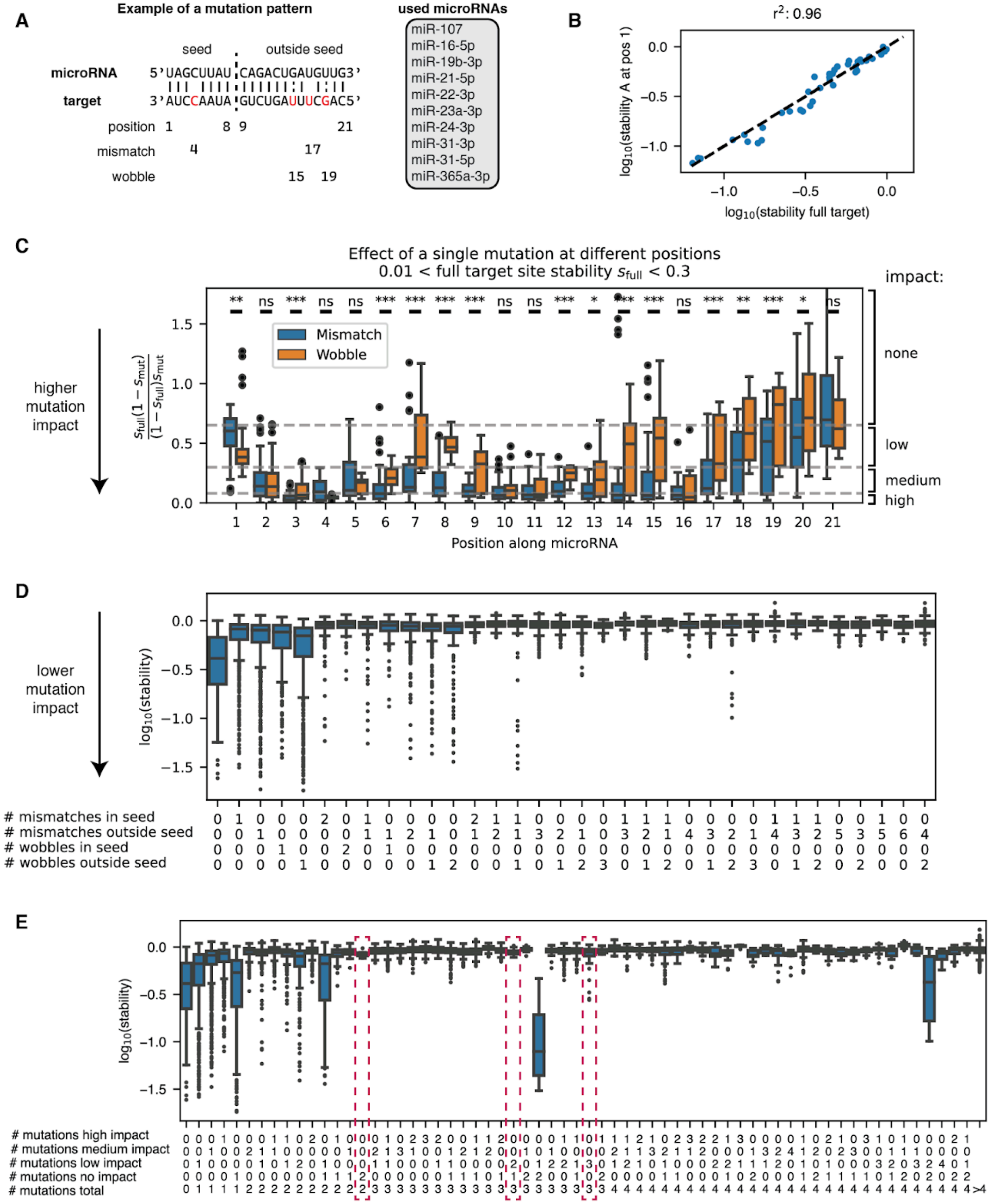


**Fig. S16. The impact of mutations can be classified based on their position in the miRNA. (A)** Example of a mutation pattern between a microRNA and a target. Note that the mutation position is denoted based on the position in the microRNA rather than in the target site although the mutations are introduced in the target site. (**B**) Stability for full target sites and those with an A mutation at position 1. A mutation to an A at the first seed-complementary nucleotide in the target does not reduce miRNA activity. (**C**) Relative loss of activity due to individual target site mutations (mismatches and wobbles). The dashed lines denote classification lines based on the median mutation impact. P-values for the difference between mismatches and wobbles were calculated using a two-sided Mann-Whitney U test. (**D**) Distribution of 3'UTR stabilities for single target sites containing multiple mutations grouped by the number and position of mutations and wobbles. (**E**) Distribution of 3'UTR stabilities for single target sites containing multiple mutations grouped by the number of mutations with a specific impact. Classification based on the mutation impact leads to a much cleaner classification than one based on the location inside or outside the seed. Dashed red boxes show examples of combined mutations with little individual impact that nevertheless strongly reduce miRNA activity on the target in combination.

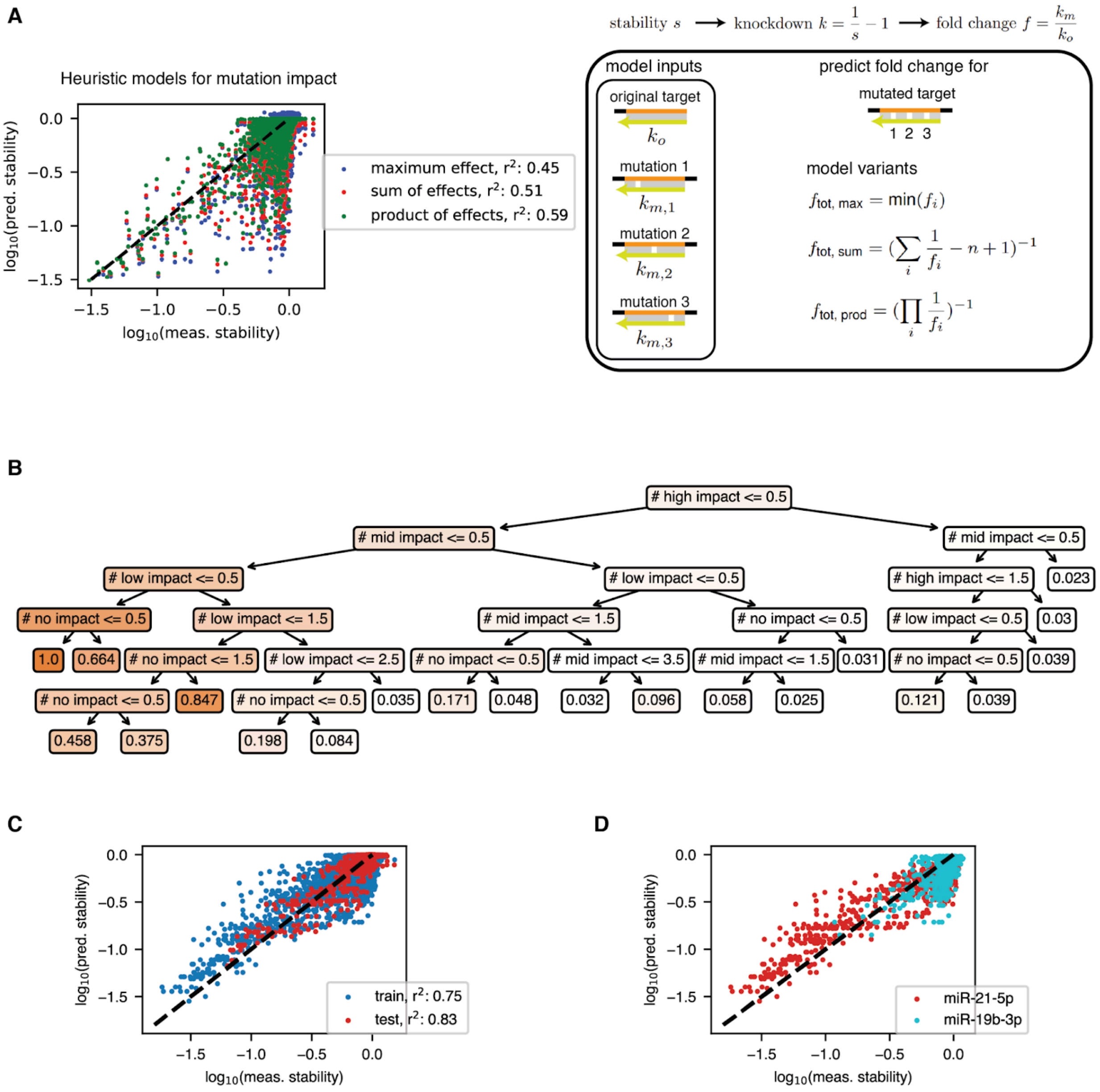


**Fig. S17. A regression tree model predicts the impact of multiple target site mutations on miRNA activity.** (**A**) The performance of three different heuristic models combining knowledge of the impact of individual mutations to calculate the overall mutation impact. None of the models work well because multiple individually inert mutations can combine to create a strong reduction in miRNA activity. (**B**) A regression tree that predicts relative knockdown values for a mutated target site from the presence and number of mutations classified as no, low, medium and high impact. (**C**) Performance of the regression tree on training and test data. The training was performed on data for all but one measured miRNA, the test data contains all mutation data for that specific miRNA (miR-31-5p). (**D**) Although the regression tree approximately captures the behavior of mutations, the behavior of mutations is strongly dependent on the individual miRNA sequence. While miR-21-5p is relatively resistant to mutations, miR-19b-3p quickly loses activity for mutated miRNAs.

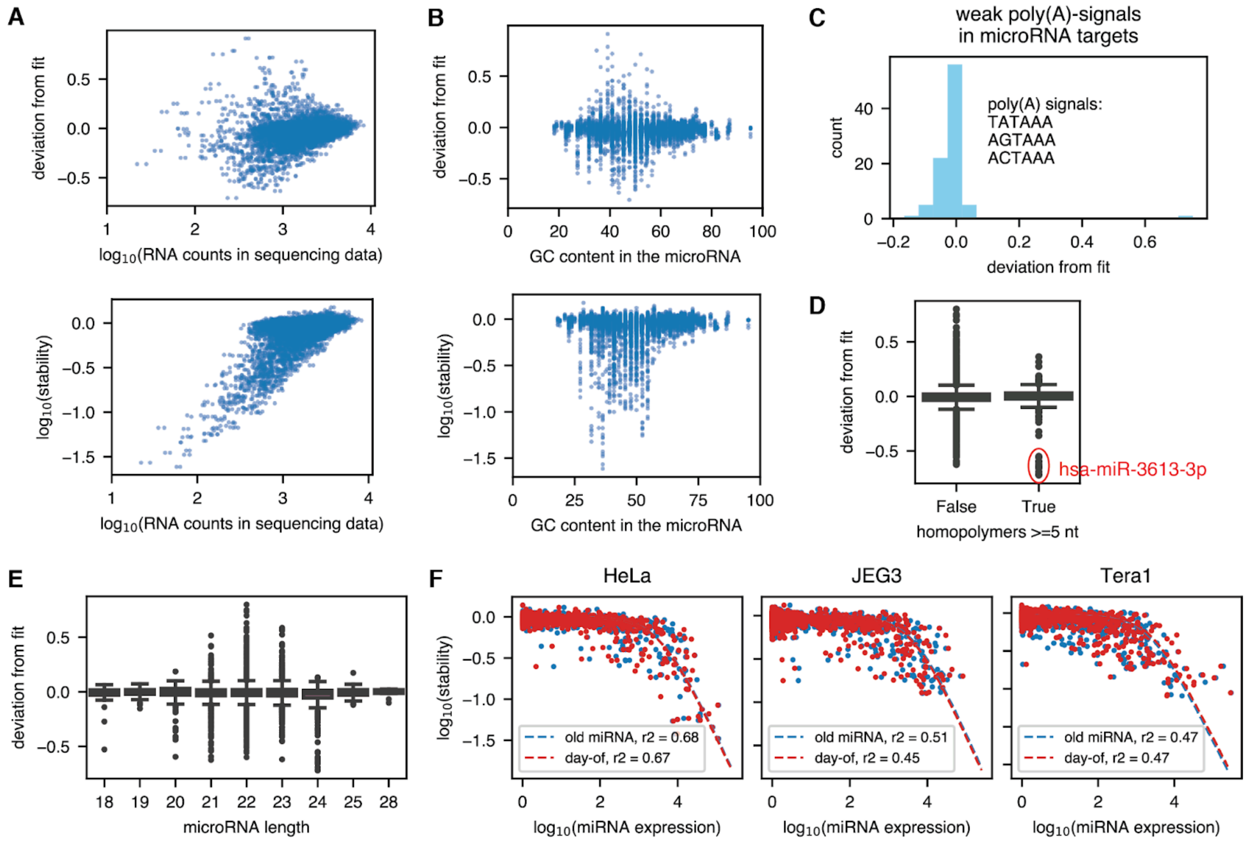


**Fig. S18. RNA counts, miRNA GC content, weak poly(A) signals, target site homopolymers, miRNA length, and inter-lab or experimental cell line variability have little effect on observed model deviations.** (**A**) Counts in the RNA sequencing data (all cell lines) versus the observed deviation from the fit and the inferred stability. Strong deviation from the fit is not explained by undercounting. (**B**) MicroRNA GC content versus the observed deviation from the fit and the inferred stability. There is no clear association between GC content and observed deviation from the fit. (**C**) Weak non-canonical poly(A) signals in the target site do not lead to a lower measured transcript stability (negative deviation). (**D**) Distribution of deviation values for microRNA targets depending on whether they contain contiguous homopolymers of a size of at least 5. Only hsa-miR-3613-3p stands out as an obvious outlier. It contains a long stretch of As: ACAAAAAAAAAAGCCCAACCCUUC. The resulting long stretch of Us in the 3'UTR probably destabilizes the transcript independent of miRNA regulation. (**E**) Deviation from the fit versus miRNA length and miRNA expression versus stability for the different miRNA lengths. In the design process, all miRNAs were standardized to a length of 21 nt by trimming from or adding uracils to the 3'end for the purpose of target site generation. We do not observe a pattern of reduced activity for shorter or longer miRNAs. (**F**) Comparison in the model fit for improved miRNA sequencing data collected either in a different laboratory (old miRNA) or from the same total RNA as for the stability assay (day-of).

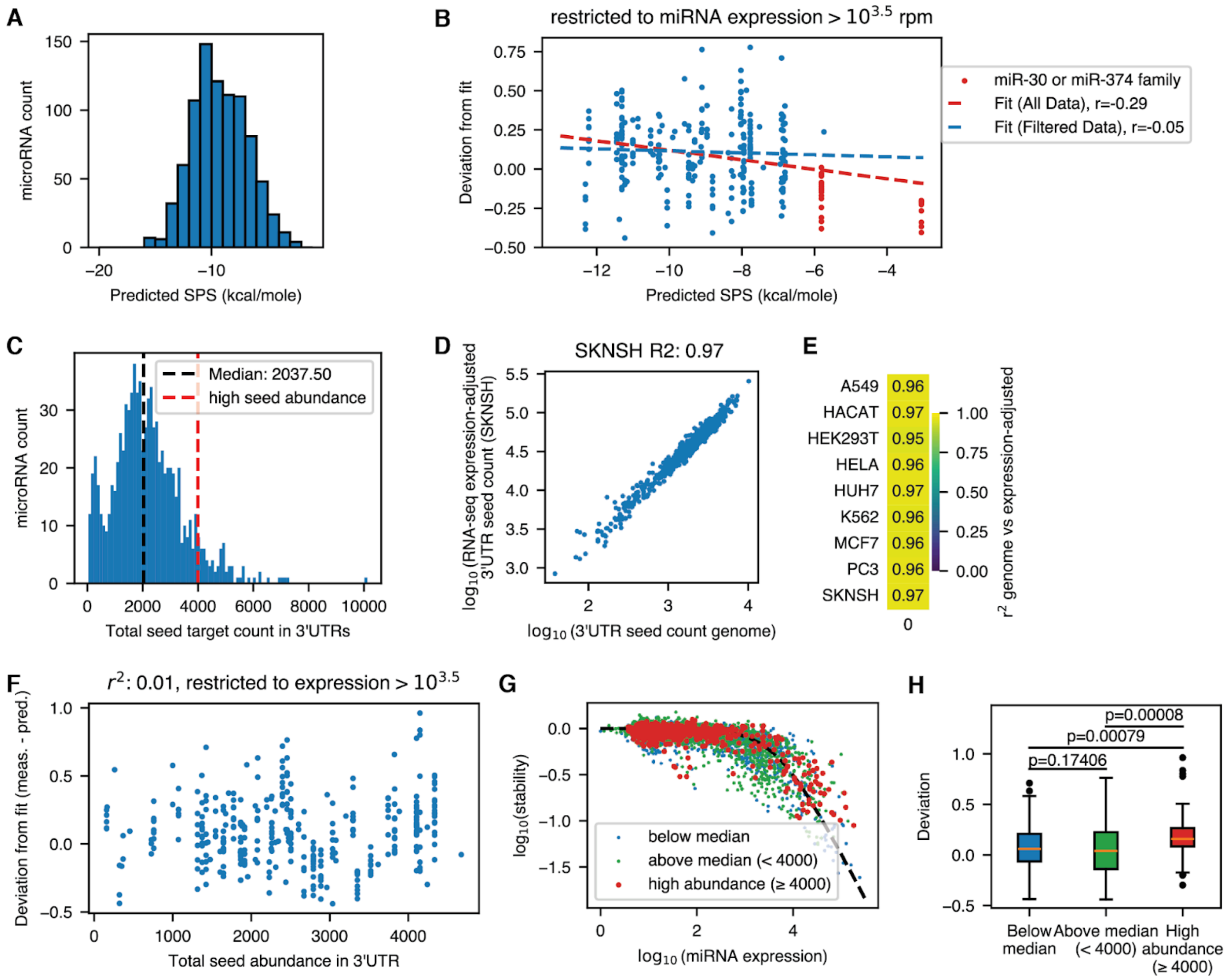


**Fig. S19. The effect of seed-pairing stability and target site abundance on microRNA targeting efficiency.** (**A**) Distribution of SPS values calculated by NUPACK. (**B**) Predicted SPS versus deviation from the fit (measured - predicted) for expressed miRNAs (>$10^{3.5}$ tpm) across all cell lines. (**C**) Distribution of seed target site counts in the 3' UTR of human genes. (**D**) Genomic 3' UTR seed target counts in the 3' UTR versus those adjusted for gene expression in SKNSH. (**E**) Pearson $r^2$ values for the association shown in D across cell lines. The high correlation justifies using genomic target counts instead of per-cell-line RNA-expression adjusted counts. (**F**) Total genomic seed abundance in the 3'UTR versus the deviation from the fit (measured - predicted) for expressed miRNAs (>$10^{3.5}$ tpm) across all cell lines. (**G**) Stability versus miRNA expression for three groups of miRNA by the abundance of 3'UTR seed targets in the genome across all cell lines. (**H**) Distribution of the deviation from the fit (measured - predicted) by seed target abundance category. p values were calculated using a two-sided Mann-Whitney U test.

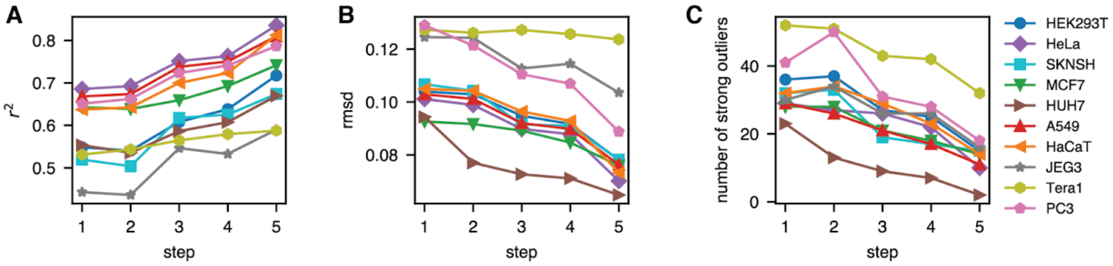


**Fig. S20. Changes in the transfer function fit by processing step and cell line.** (**A**) Pearson $r^2$ value. (**B**) Root mean square deviation (rmsd). (**C**) Number of strong outliers whose measured stability deviates by more than a factor of 2 from the predicted value. 1: unscaled microarray data, 2: scaled microarray data, 3: combination of microarray and sequencing data, 4: bias-aware merging, 5: removal of crosstalk.

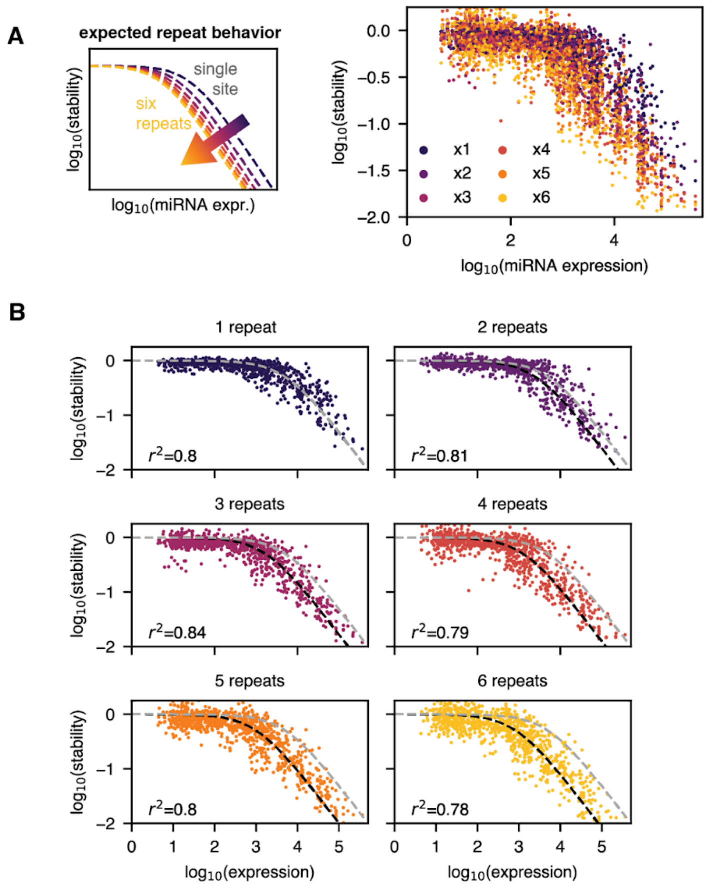


**Fig. S21. The behavior of target repeats follows the additive model.** (**A**) Stability versus miRNA expression for different repeat numbers across all measured cell lines. The plot on the left shows the expected behavior according to the additive model. (**B**) Stability versus miRNA expression for individual repeat numbers. The transfer function (dashed black line) uses the fitted parameters from Fig. 1. The dashed gray line shows the prediction for a single target site. The input miRNA expression is multiplied by the repeat number.

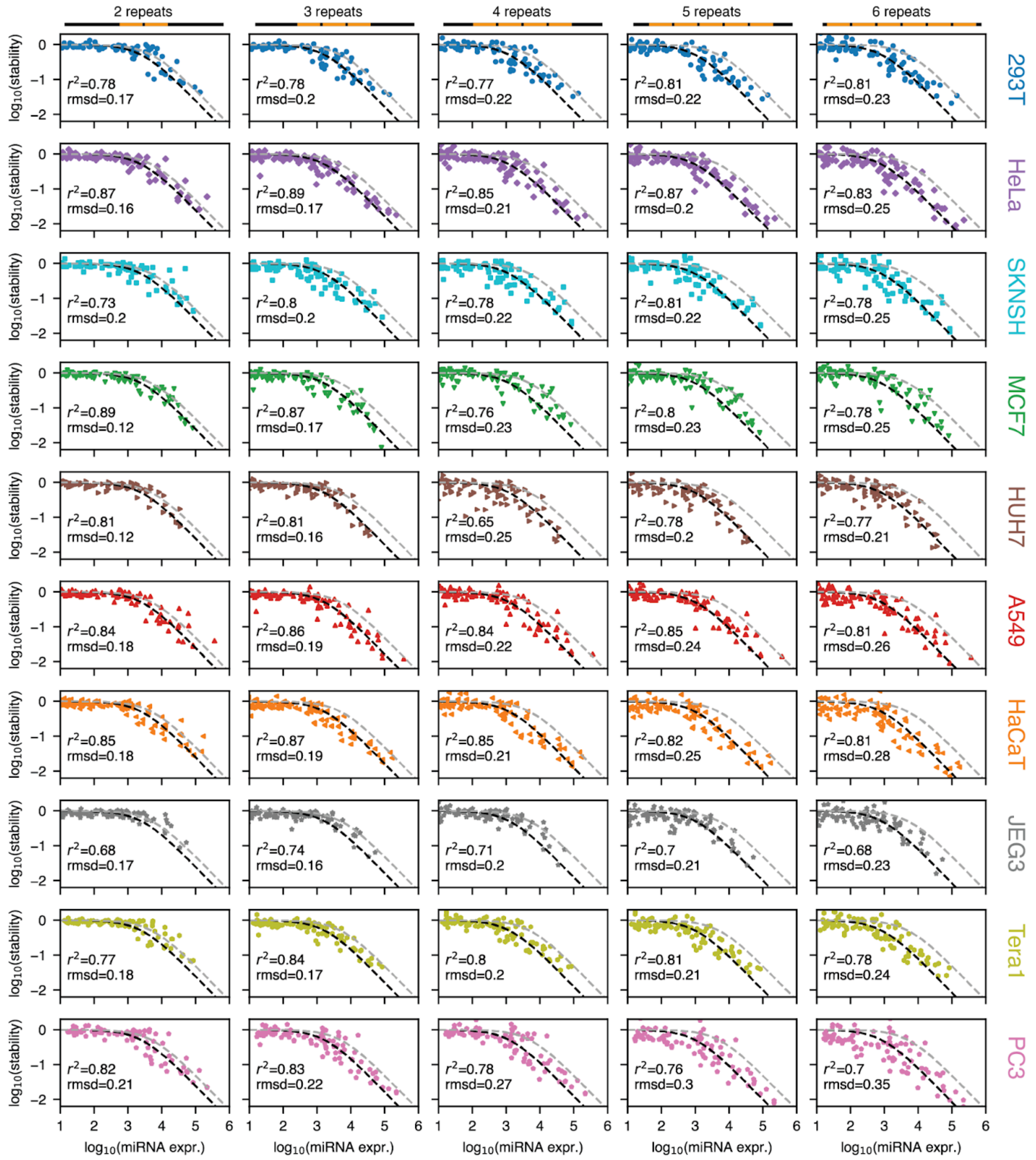


**Fig. S22. Predictions of the additive model for two to six repeats of a single target site.** The transfer function (dashed black line) uses the fitted parameters from Fig. 1. The dashed gray line shows the prediction for a single target site.

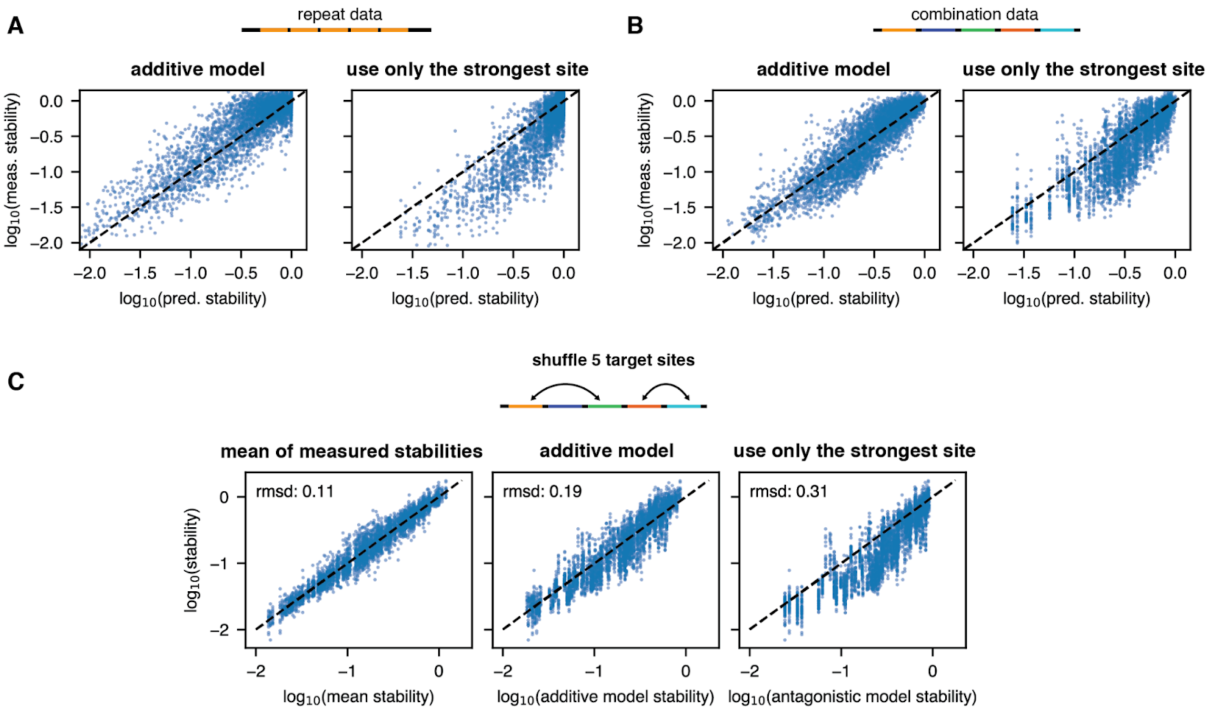


**Fig. S23. Comparison of the additive model and an antagonistic model using only the strongest site for repeat and combination data.** All plots use inferred expression values derived by inversion of the transfer function. We compare the additive model discussed in the main text and an antagonistic model that only uses the strongest target site (i.e., the one whose cognate miRNA has the highest expression) to predict stability. (**A**) Predictions of the two models and measurements for target repeats. (**B**) Predictions of the two models and measurements or combinations of different targets. (**C**) Model comparison for shuffled target sites. On the left, we show predictions based on the measured mean stabilities of the 15 shuffled variants per set of target sites.

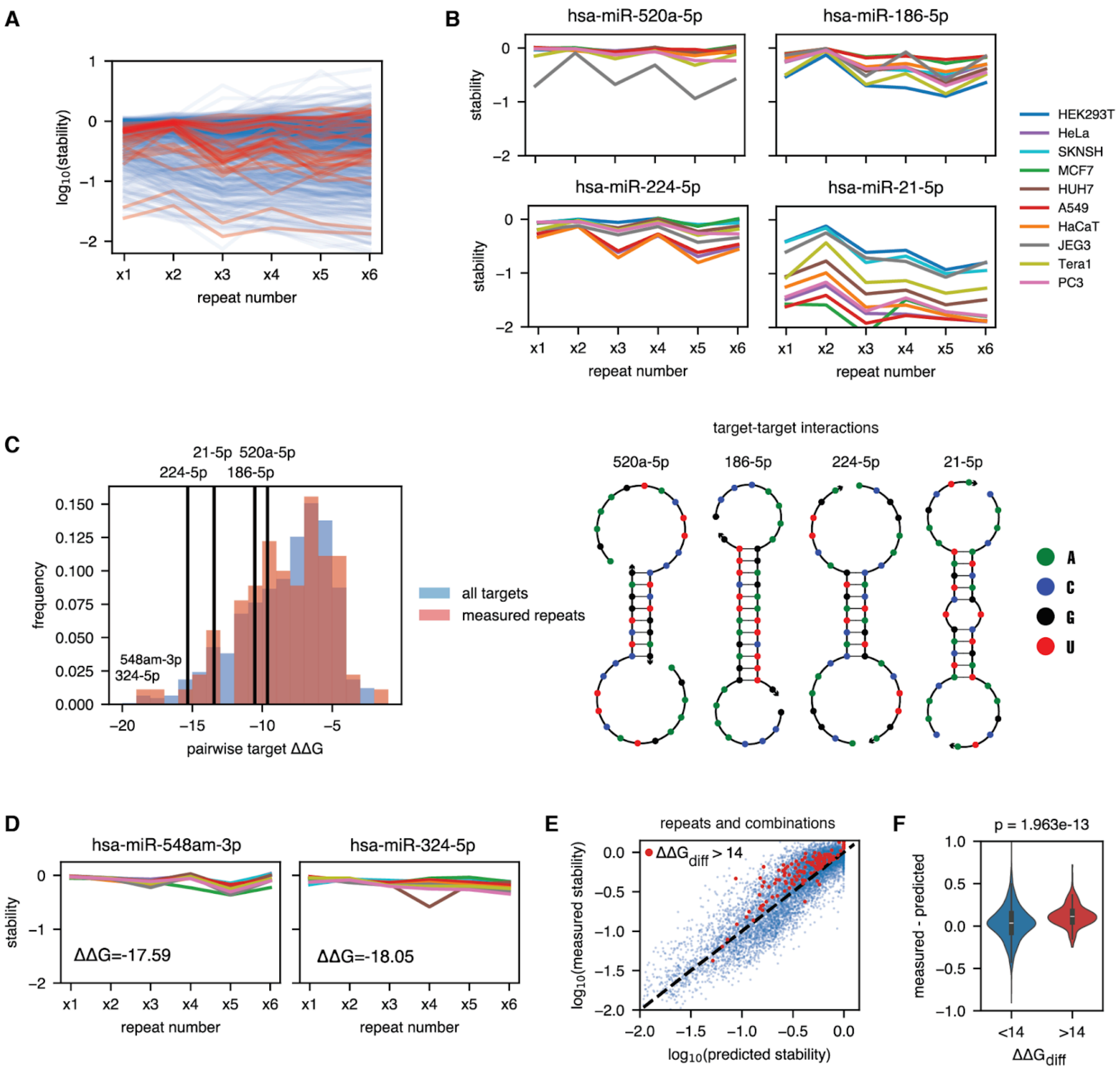


**Fig. S24. Interactions between target sites explain outliers in the repeat stability data.** (**A**). Stability of constructs containing one to six repeats of a single target site versus the number of repeats. Each solid line is for a single miRNA. Constructs with an unexpected pattern in which stabilities non-monotonically both increase and decrease at least twice with the number of repeats are highlighted in red. (**B**) Four example miRNA targets showing unexpected repeat stability patterns. There is a relative increase in stability for an even number of repeats. (**C**) Left: Distribution of binding energies for two identical targets for all miRNAs (blue, high confidence in miRBase or in MirGeneDB) and of all miRNA targets we chose for measuring target repeats (Fig. 2B, S22). The interaction energies of the outliers shown in (B) and of the two most strongly self-interacting miRNA targets are highlighted. Right: Predicted secondary structures for target-target interactions for miRNAs shown in (B). (**D**) Observed stability patterns for the two most strongly self-interacting miRNA targets. These miRNAs do not show unusual stability patterns in our measured stability data because they are not significantly expressed in any of the cell lines. (**E**) Predicted versus measured stability for all measured repeats and combinations. Measurements where the dominant miRNA target is expected to be obscured by strong secondary structure are highlighted in red. (**F**) Difference between measured and predicted stability for constructs without and with strong secondary structure.

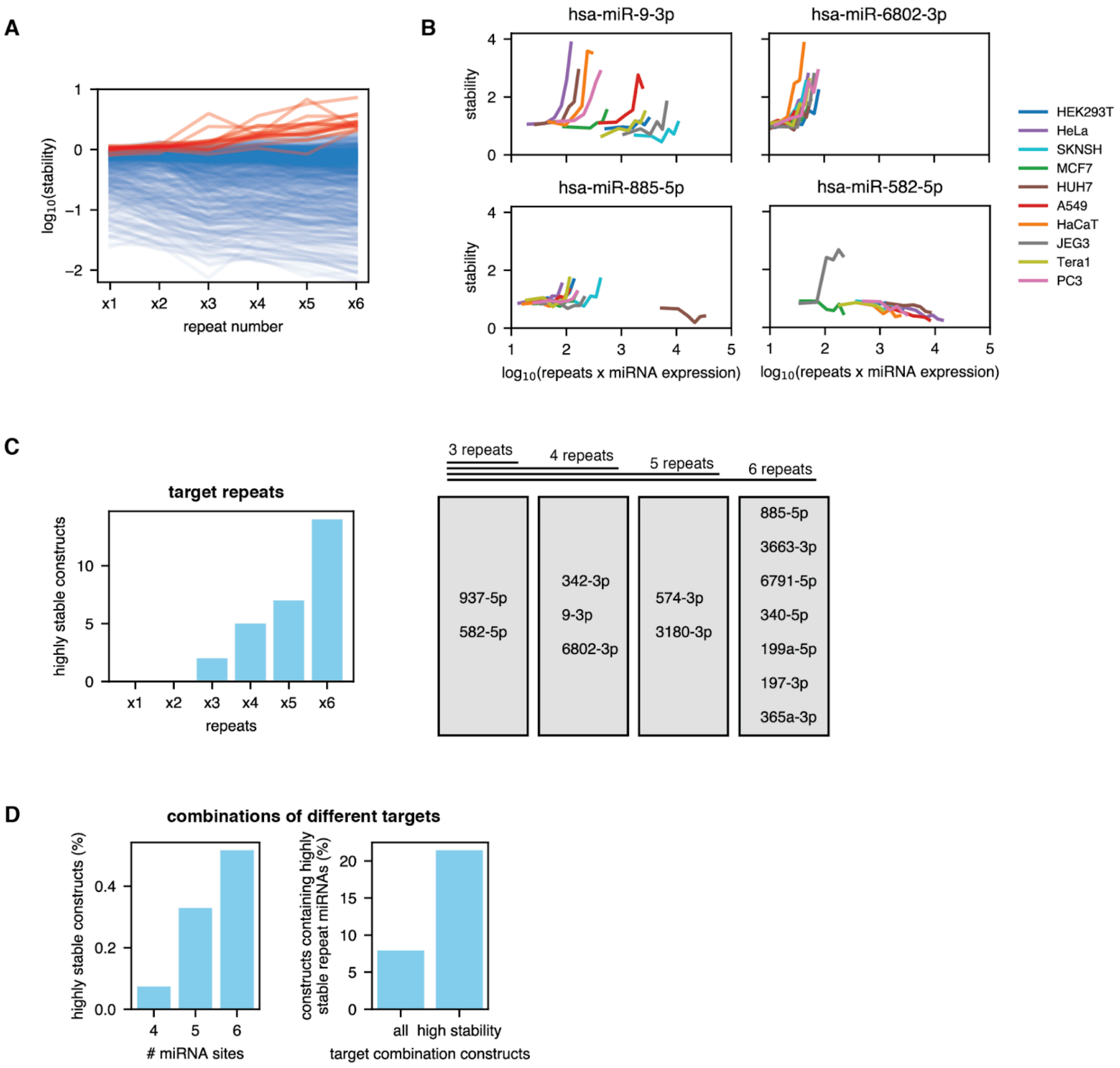


**Fig. S25. Some miRNA target repeats cause strong increases in stability.** (**A**) Stability of constructs containing one to six repeats of a single target site versus the miRNA expression. Each solid line is for a single miRNA. Constructs with an unexpected pattern in which there is a strong increase in stability with the repeat number are highlighted in red. (**B**) Four example miRNA targets that show a strong increase in stability in at least one cell line. Stability increases are more pronounced in cell lines where the cognate miRNA is not expressed. (**C**) The number of constructs with very high stability (>1.5) in at least one cell line increases with the number of repeats. All constructs that have high stability at a lower repeat number also show high stability at all higher repeat numbers. Stability-increasing miRNA targets therefore behave consistently across repeat numbers. (**D**) For combinations of different miRNA targets, the number of constructs with very high stability also increases with the number of target sites (left plot). The fraction is far lower than for repeats of a single target site. Among those combinations that do have high stability, a greater fraction contains miRNA target sites that increase stability in constructs with four or more repeats (right plot).

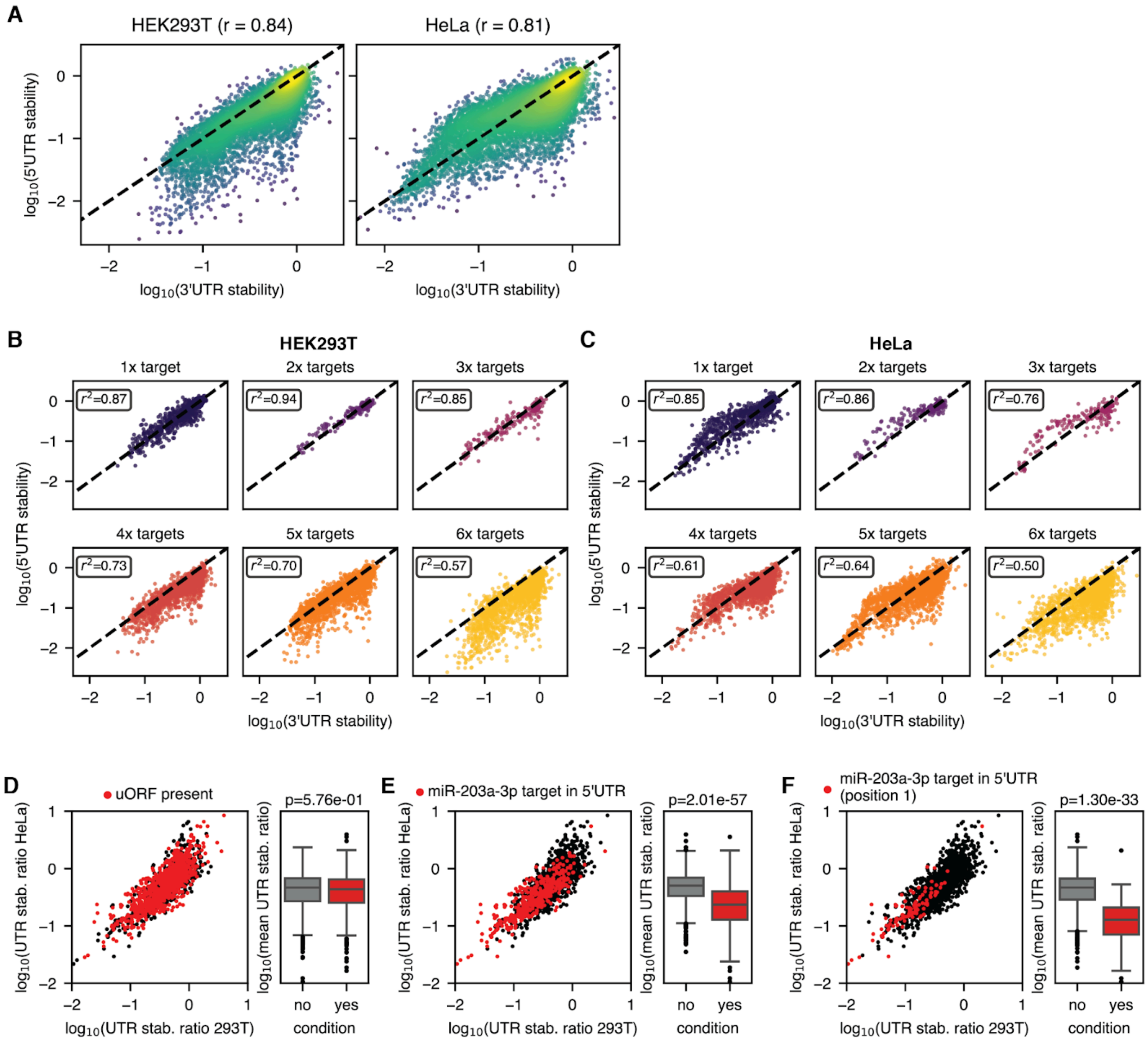


**Fig. S26. Comparison of stability for 5' and 3'UTR target sites.** Stabilities measured in the 5'UTR versus stabilities measured in the 3'UTR for (**A**) all library members in 293T and HeLa cells, (**B**) different total numbers of target sites for HEK293T, and (**C**) different total numbers of target sites for HeLa. The impact of (**D**) an upstream open reading frame (uORF), (**E**) at least one miR-203a-3p target site, (**F**) a miR-203a-3p target site on the position closest to the 5'UTR on the ratio of the stability in the 5'UTR and the 3'UTR.

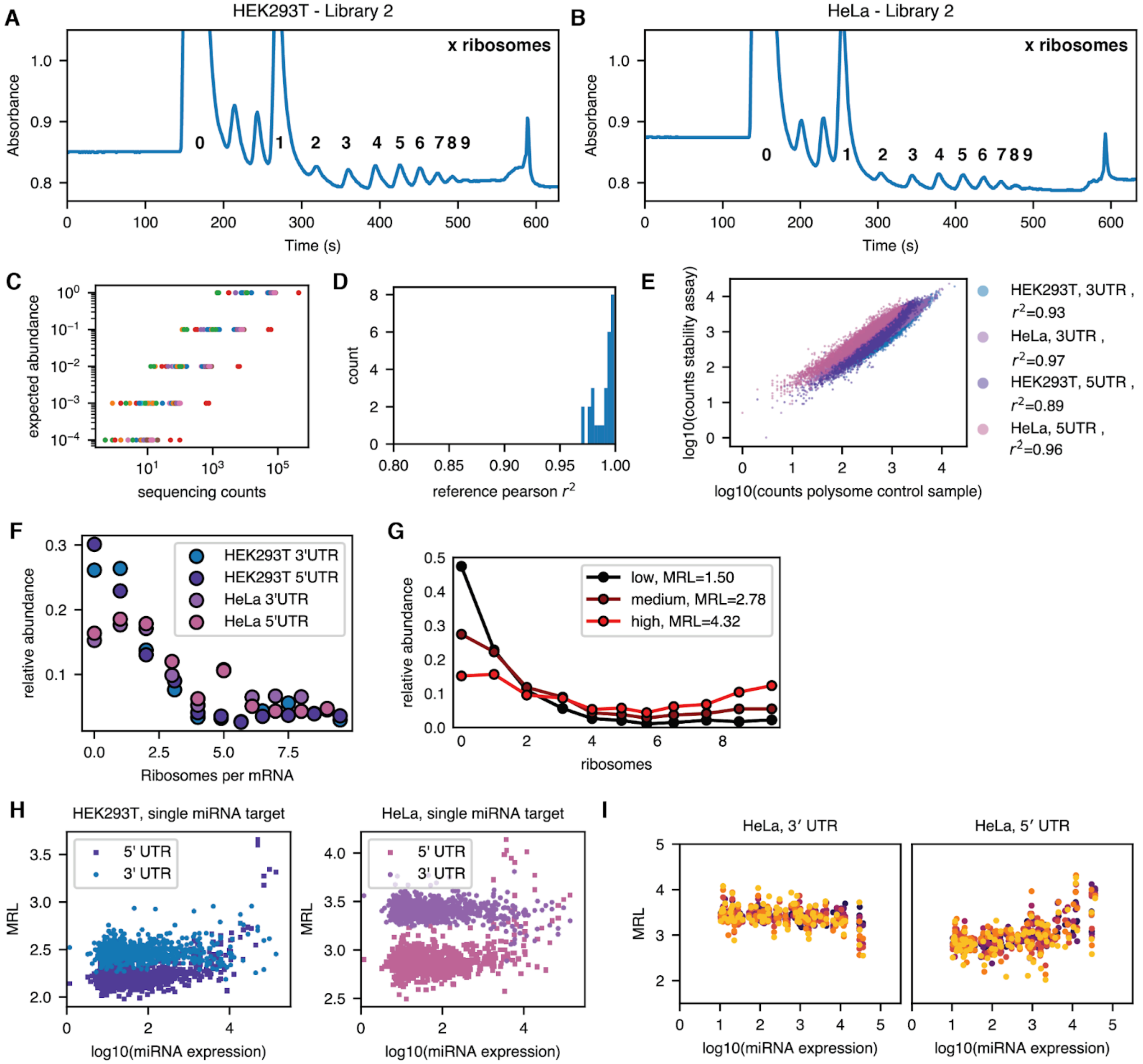


**Fig. S27. Polysome profiling of plasmid library mRNA.** Absorbance trace and ribosome peak assignment for the (**A**) HEK293T and (**B**) HeLa libraries. (**C**) Expected abundance and sequencing counts for spike-ins used to normalize the counts across different polysome fractions. (**D**) Pearson correlations of the values in C. (**E**) Counts in the control samples of the polysome assay versus counts in the ratiometric stability assay. (**F**) Distribution of relative abundances across different polysome fractions for the four tested libraries. (**G**) Examples of low, medium, and high MRL samples and their ribosome distribution. (**H**) MRL versus microRNA expression for all single target site designs. (**I**) Impact of different numbers of repeats of microRNA target sites versus the expression of their associated microRNA in HeLa.

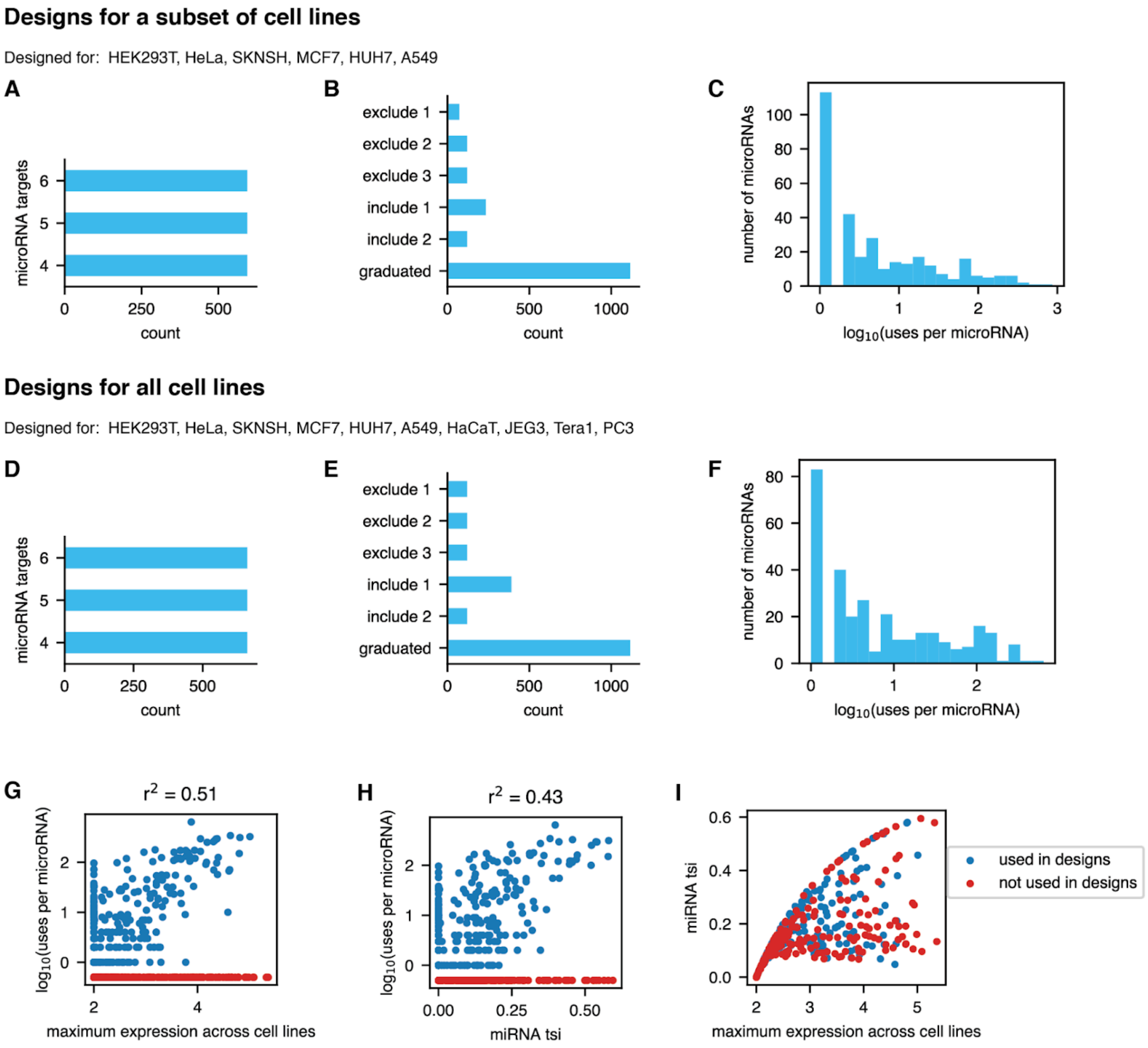


**Fig. S28. Statistics on the generated designs.** We created designs for either six of the ten cell lines (A)-(C) or all measured cell lines (D)-(I). (**A**)/(**D**). The number of designs with 4, 5, or 6 miRNA target sites. (**B**)/(**E**). The number of designs for different binary design targets including or excluding specific cell lines or graduated stability patterns. (**C**)/(**F**) Distribution of the number of times a specific miRNA target occurs across all designs. (**G**) The maximum expression level of a miRNA across cell lines and its usage frequency in our designs. More highly expressed miRNAs are used more often but many highly expressed miRNAs are nevertheless not used in any design. The correlation value is only calculated for used miRNAs. (**H**) Association of miRNA usage in our designs with the tissue-specificity index (tsi) (*62*) of the miRNA across our measured cell lines. More tissue-specific miRNAs are used more often but many tissue-specific miRNAs are not used in any of our designs. (**I**) Maximum expression versus tissue-specificity for miRNAs that were either used or not used for our designs.

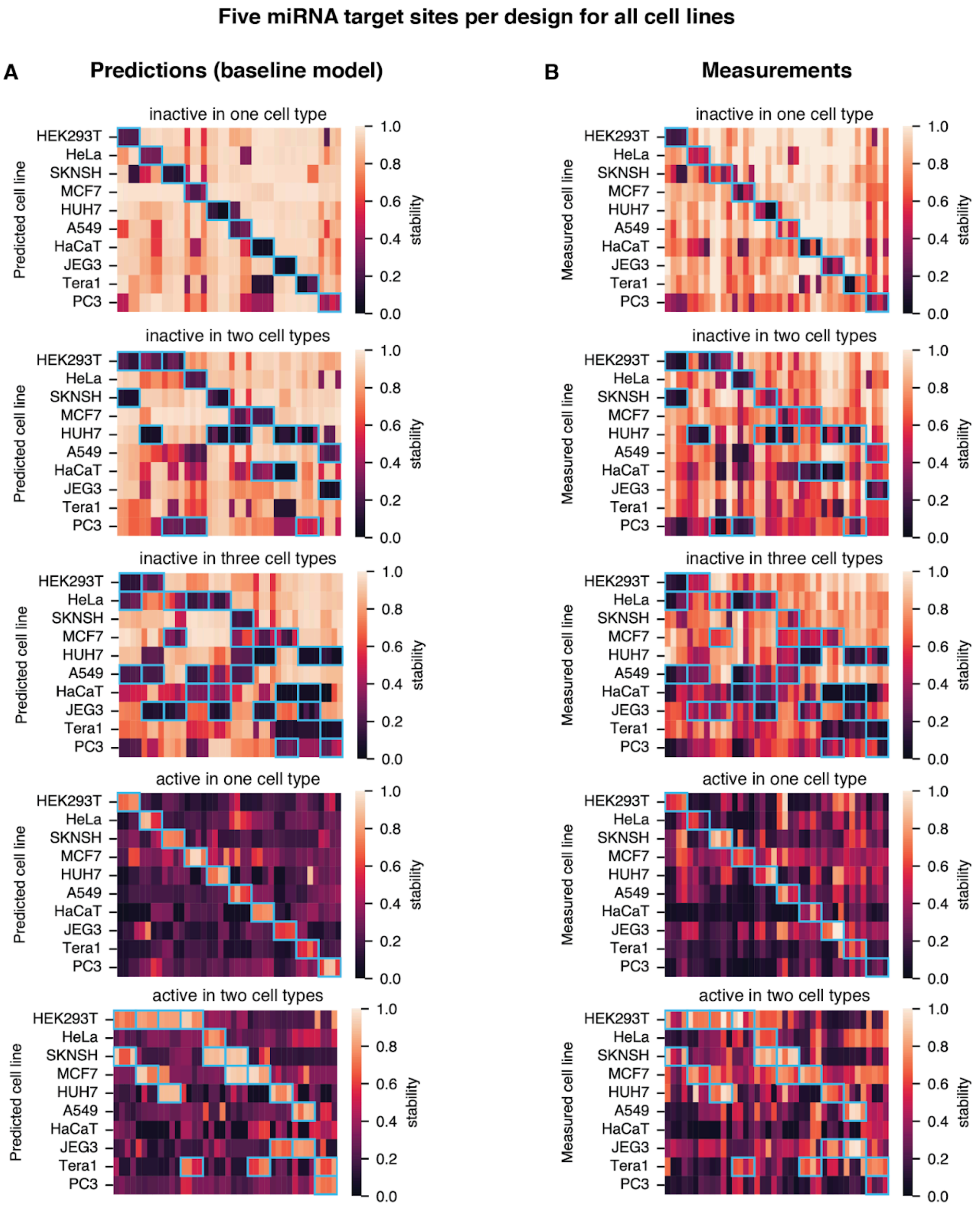


**Fig. S29. Predictions and measurements for all mse-based binary designs for all cell lines with five microRNA target sites.** We generated four designs per target pattern (e.g., inactivity in a single cell line) for each of the five design types. Each column shows one design. (**A**) Stabilities predicted by the baseline model and (**B**) measured stabilities across the cell lines. The blue boxes indicate the cell lines in which designs are meant to be active or inactive.

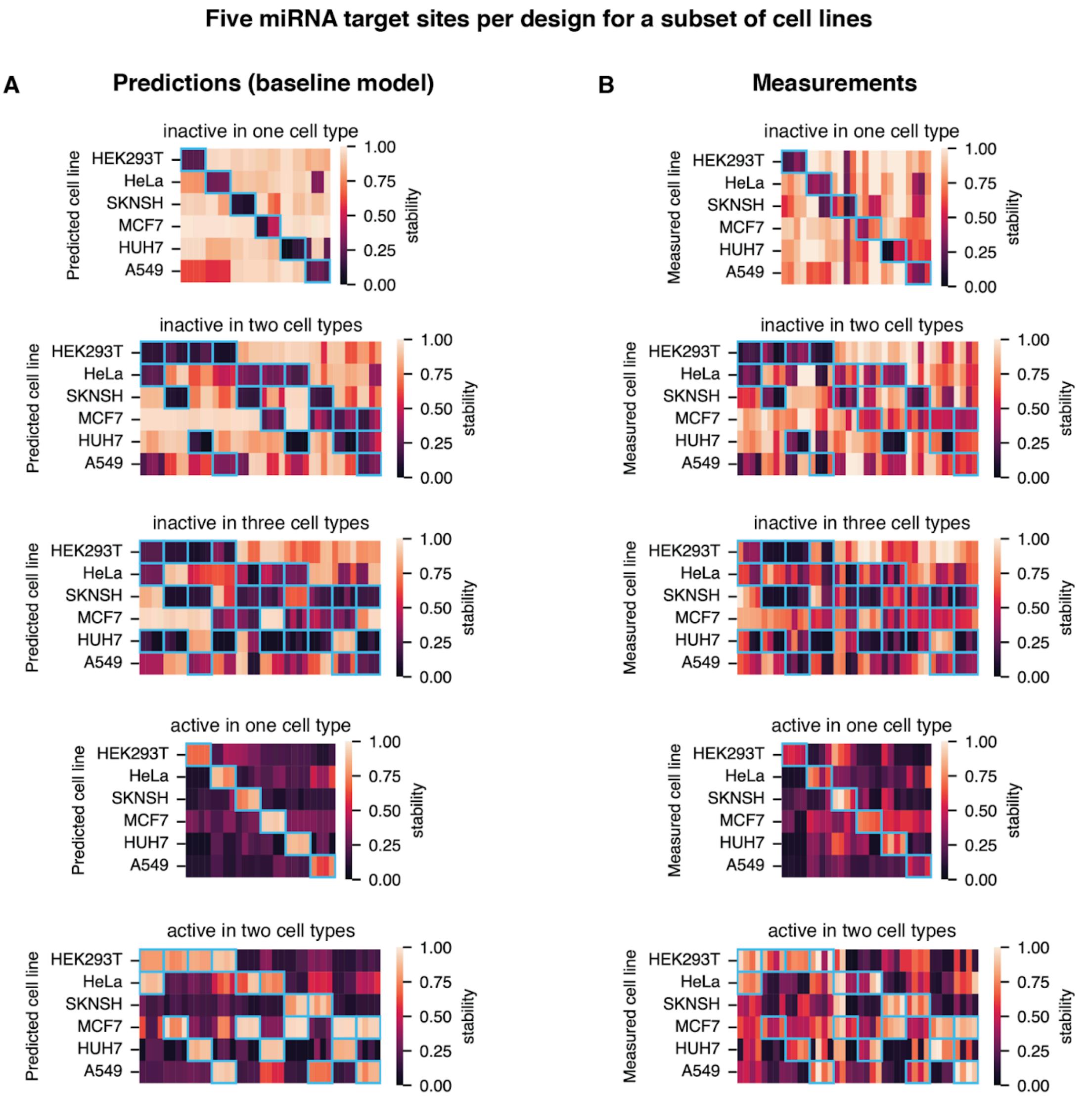


**Fig. S30. Predictions and measurements for all mse-based binary designs for a subset of cell lines with five microRNA target sites.** We generated four designs per target pattern (e.g., inactivity in a single cell line) for each of the five design types. Each column shows one design. (**A**) Stabilities predicted by the baseline model and (**B**) measured stabilities across the cell lines. The blue boxes indicate the cell lines in which designs are meant to be active or inactive.

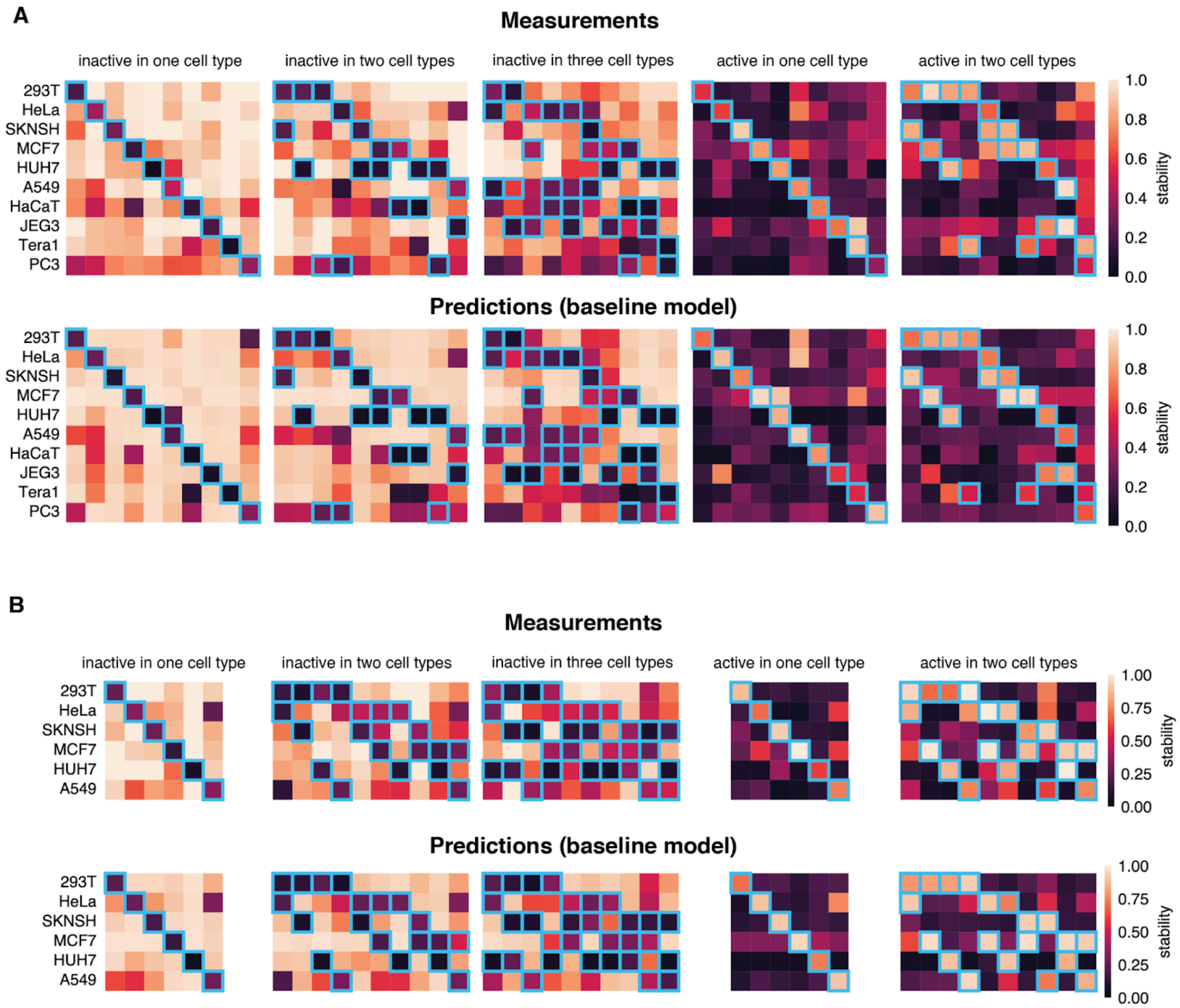


**Fig. S31. Measurements and predictions of the baseline model for the best-performing binary designs.** We show one design per target cell line and design type with the smallest measured weighted mean-squared error between the target stabilities and the measurement. Designs were created for (**A**) all cell lines or (**B**) a subset of 6 cell lines. Design failures are generally predicted in advance even by the baseline model. The blue boxes indicate the cell lines in which designs are meant to be active or inactive.

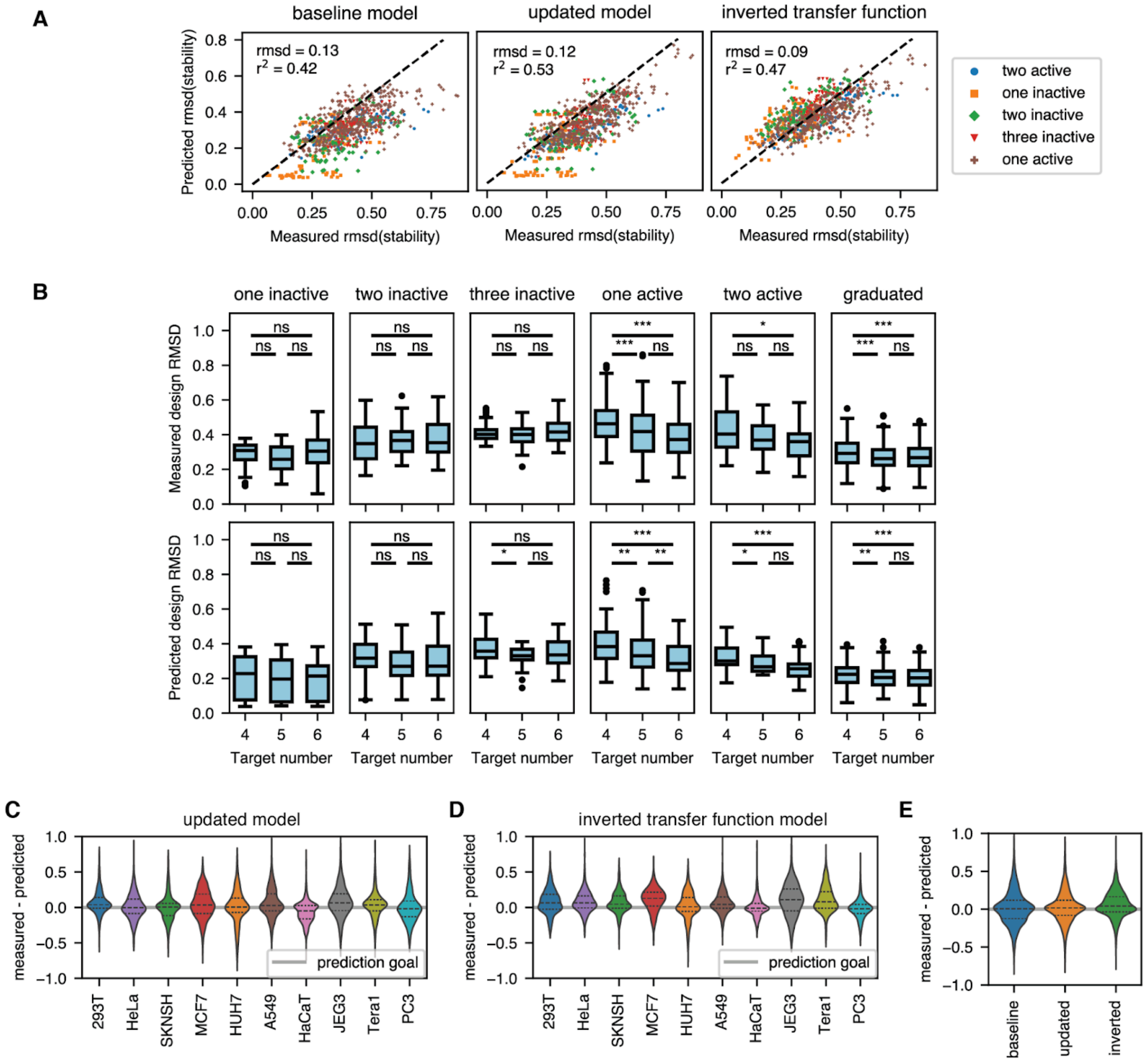


**Fig. S32. Performance of different prediction models.** (**A**) Measured and predicted design rmsd values for the different binary designs. (**B**) Measured and predicted design rmsd (baseline model) for different numbers of target sites. The significance was calculated using a two-sided Mann-Whitney U test. *= p<0.05, **=p<0.01, ***=p<0.001, ns=not significant (**C**) Deviations of measurements from the predictions of the updated model. (**D**) Deviations of measurements from the predictions of the inverted transfer function model. (**E**) Deviations of measurements from predictions for the three models averaged across cell lines.

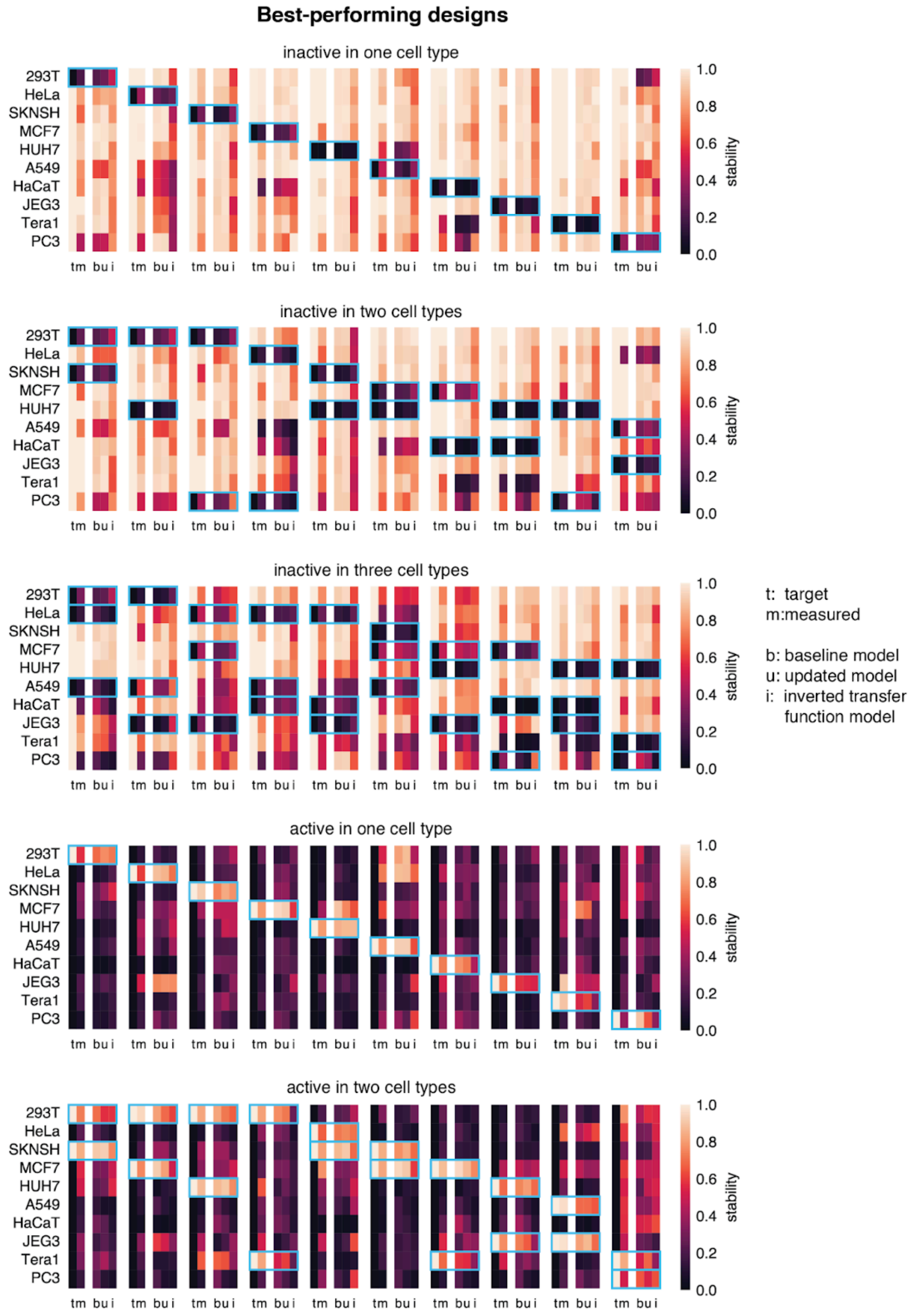


**Fig. S33. Measurements and predictions by three different models for the best-performing binary designs targeting all cell lines.** We show one design per target cell line and design type. We show the target and measured stability as well as predictions by the three different prediction models. The blue boxes indicate the cell lines in which designs are meant to be active or inactive. t: target stability, m: measured stability; b: baseline, u: updated, and i: inverted transfer function model.

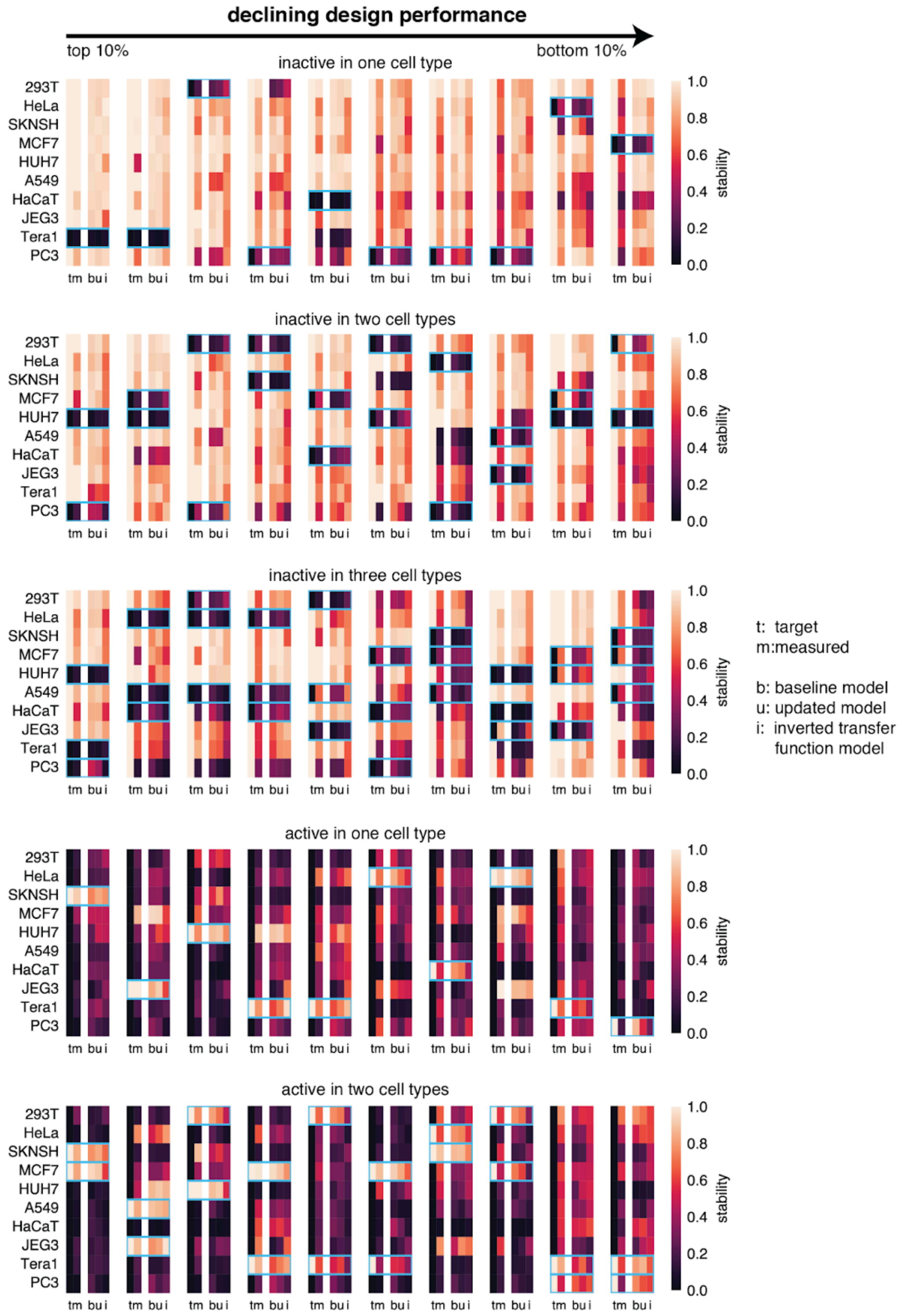


**Fig. S34. Measurements and predictions by three different models for binary designs with a range of performances.** The figure shows one design from each decile of design performance for each design type. We show the target and measured stability as well as predictions by the three prediction different models. The blue boxes indicate the cell lines in which designs are meant to be active or inactive. t: target stability, m: measured stability; b: baseline, u: updated, and i: inverted transfer function model.

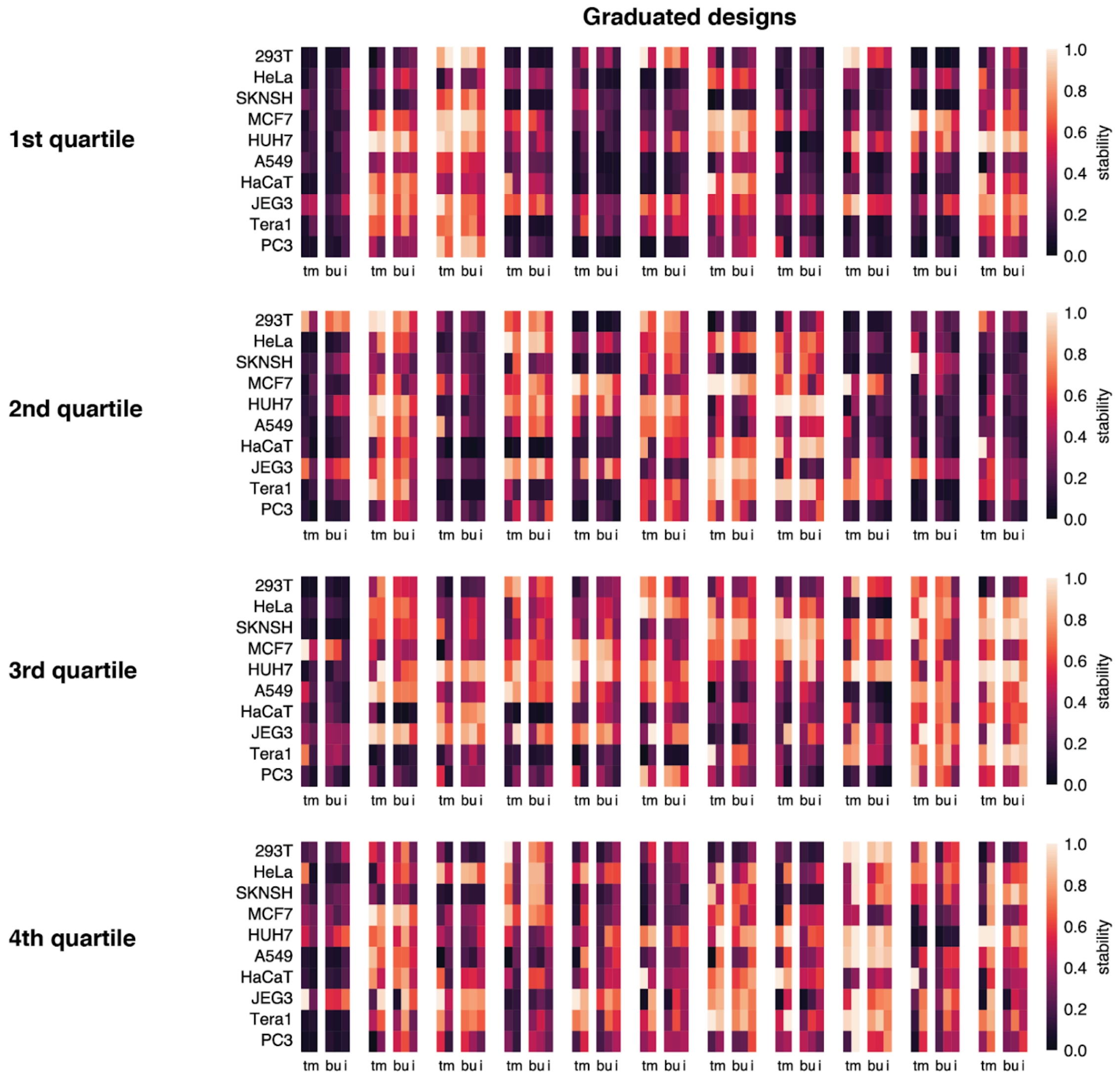


**Fig. S35. Measurements and predictions by three different models for graduated designs across design performance quartiles.** (**A**). We show ten designs for each quartile of design performance. We show the target and measured stability as well as predictions by the three different prediction models. t: target stability, m: measured stability; b: baseline, u: updated, and i: inverted transfer function model.

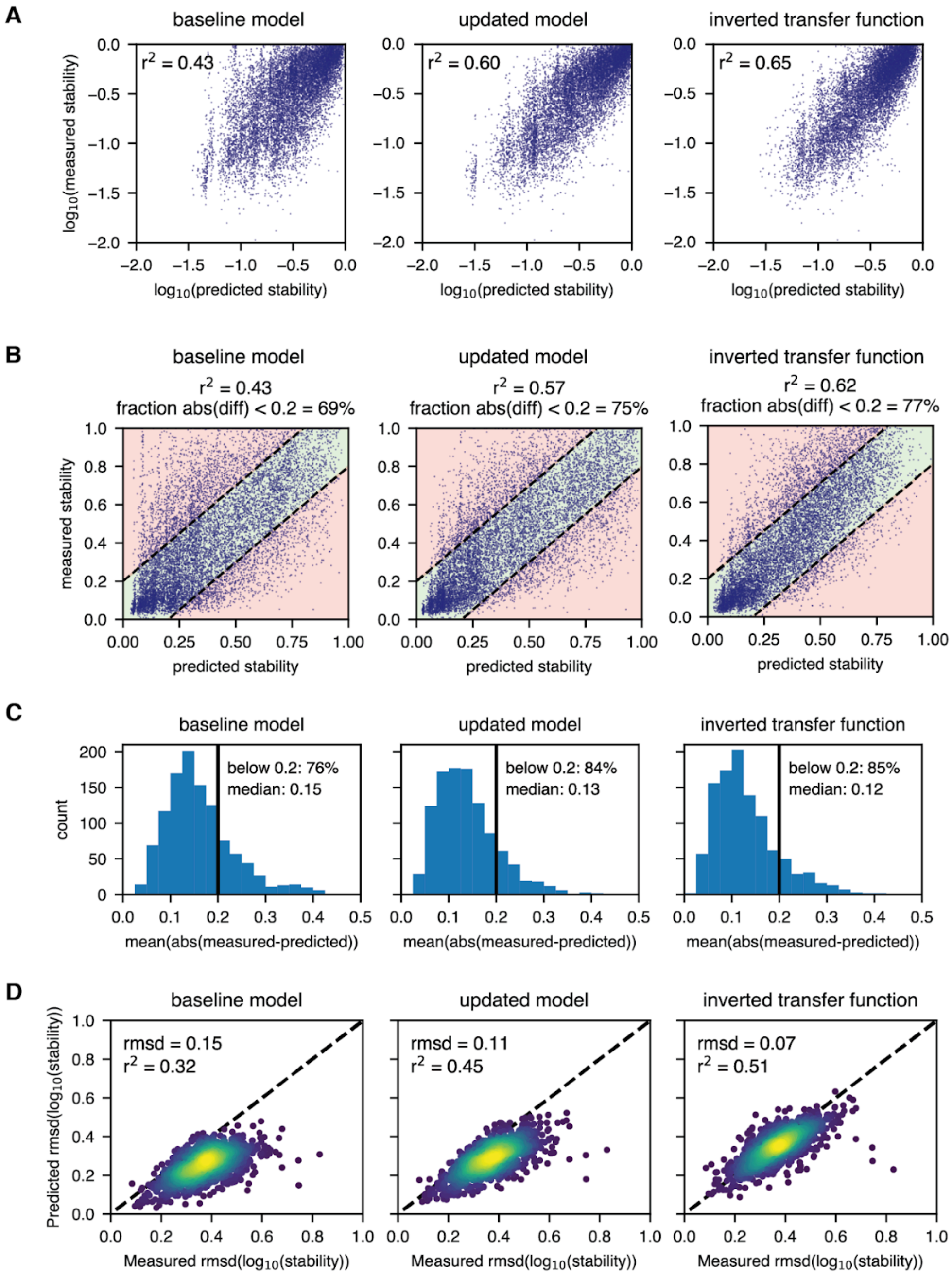


**Fig. S36. Prediction performance by different models for graduated designs targeting all cell lines.** (**A**) Predicted and measured logarithmic stabilities. (**B**) Predicted and measured linear stabilities. The area where the difference between prediction and measurement is less than 0.2 is shaded green, the rest is shaded red.(**C**) Absolute prediction error per design averaged across cell lines. (**D**) Predicted and measured rmsd values to the target $\log_{10}$ stabilities for all graduated designs.

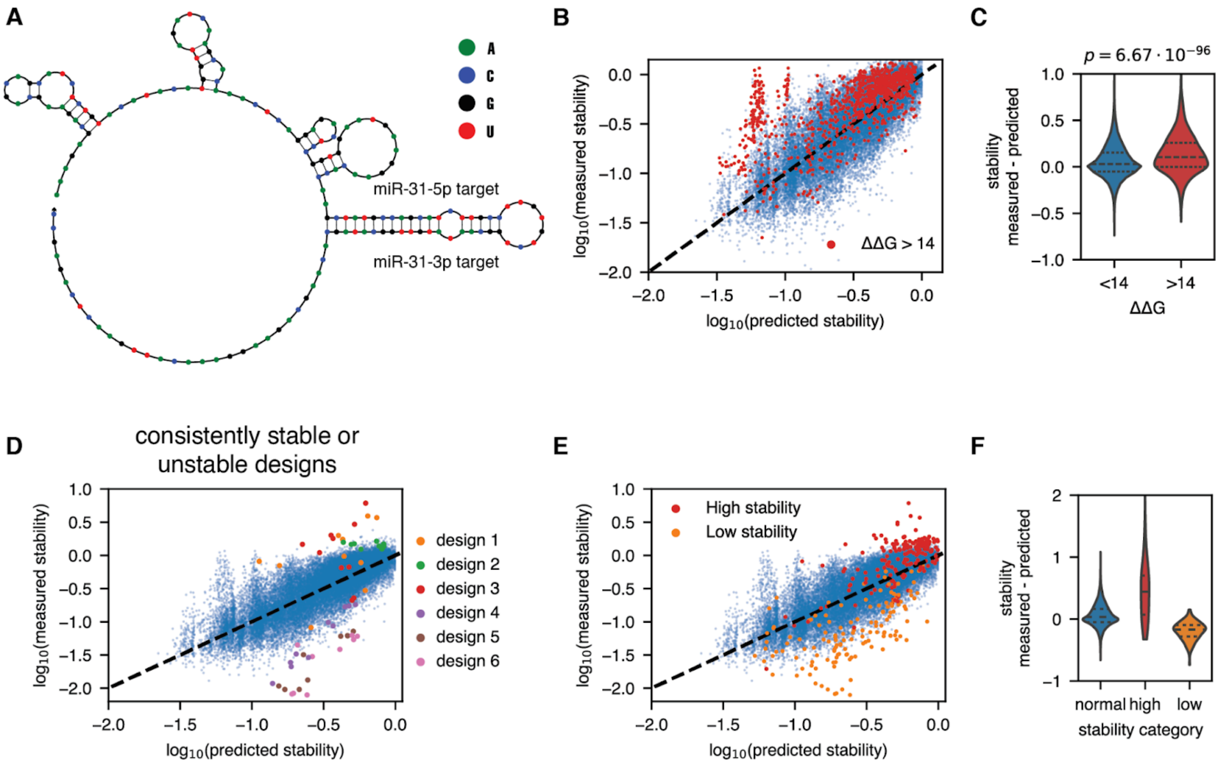


**Fig. S37. Secondary structure and global stability explain some prediction failures.** (**A**) Example of a design with strong secondary structure due to the use of the 5p and 3p arm of the same miRNA. (**B**) Predicted (inverted transfer function model) and measured stabilities for all designs targeting all cell lines. Designs with strong secondary structure in a dominant miRNA target site are highlighted. (**C**) Difference between measured and predicted stabilities for low and high $\Delta\Delta G$ designs. The p-value was calculated by a Mann-Whitney U test. (**D**) Three particularly stable and three particularly unstable designs in the prediction plot for the inverted transfer function model. The designs tend to be excessively stable or unstable across all measured cell lines. (**E**) All high stability (stability larger than 1.5 in any cell line) and low stability (ratio between predicted and measured stability larger than 6.3 in any cell line) designs. (**F**) Difference between measured and predicted stabilities for normal, high stability, and low stability designs.

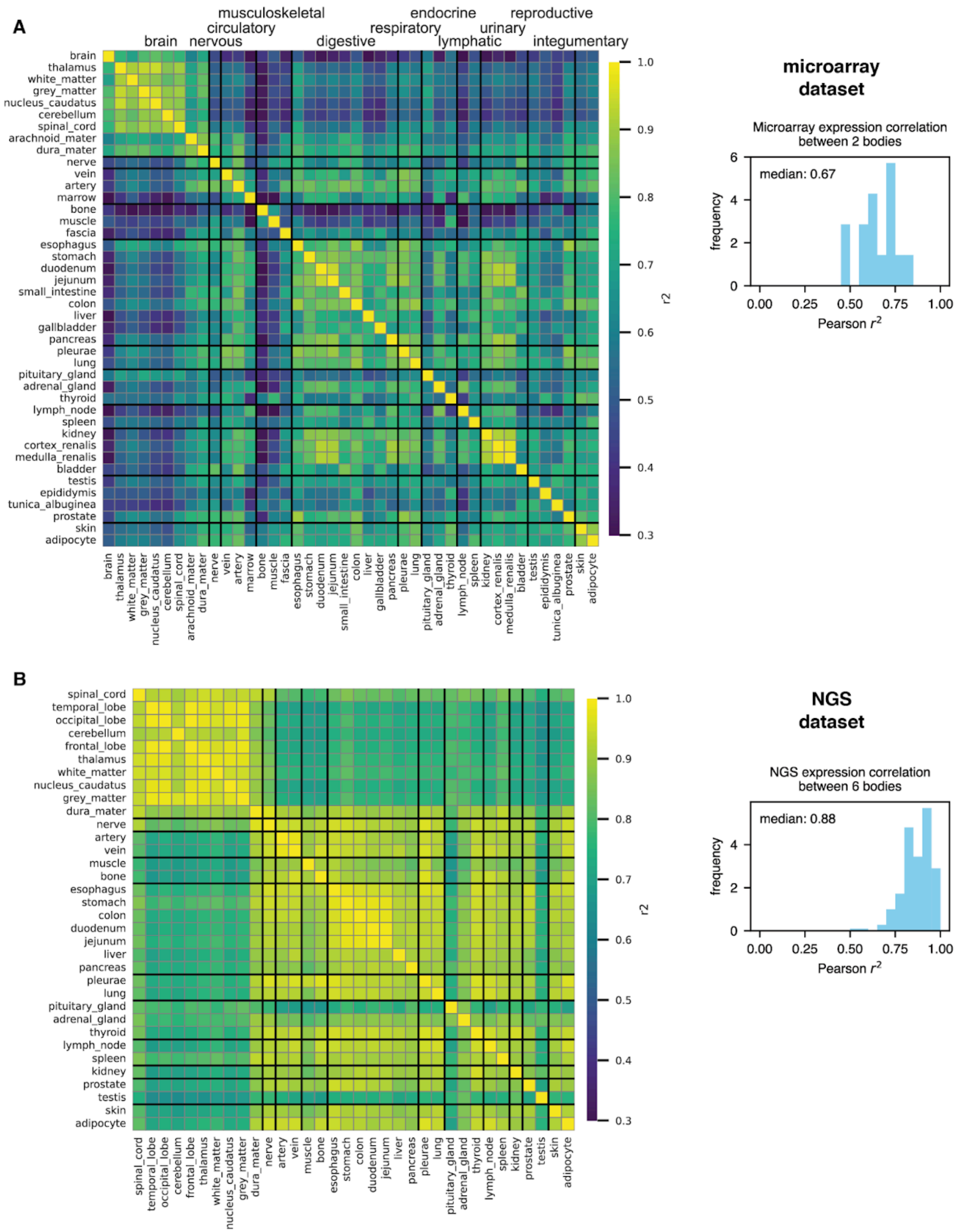


**Fig. S38. Comparison of human tissue microRNA expression datasets.** (**A**) Pearson correlation values between different tissues in the microarray dataset. The microarray tissue dataset was generated by merging expression values for 2 different subjects via a geometric mean. The correlation for the same tissue between the expression values measured for the two subjects before merging is shown on the right. (**B**) Pearson correlation values between different tissues in the NGS dataset. The dataset was generated by merging expression values for 6 subjects via a geometric mean. The correlation for the same tissue between the expression values measured for the six subjects before merging is shown on the right.

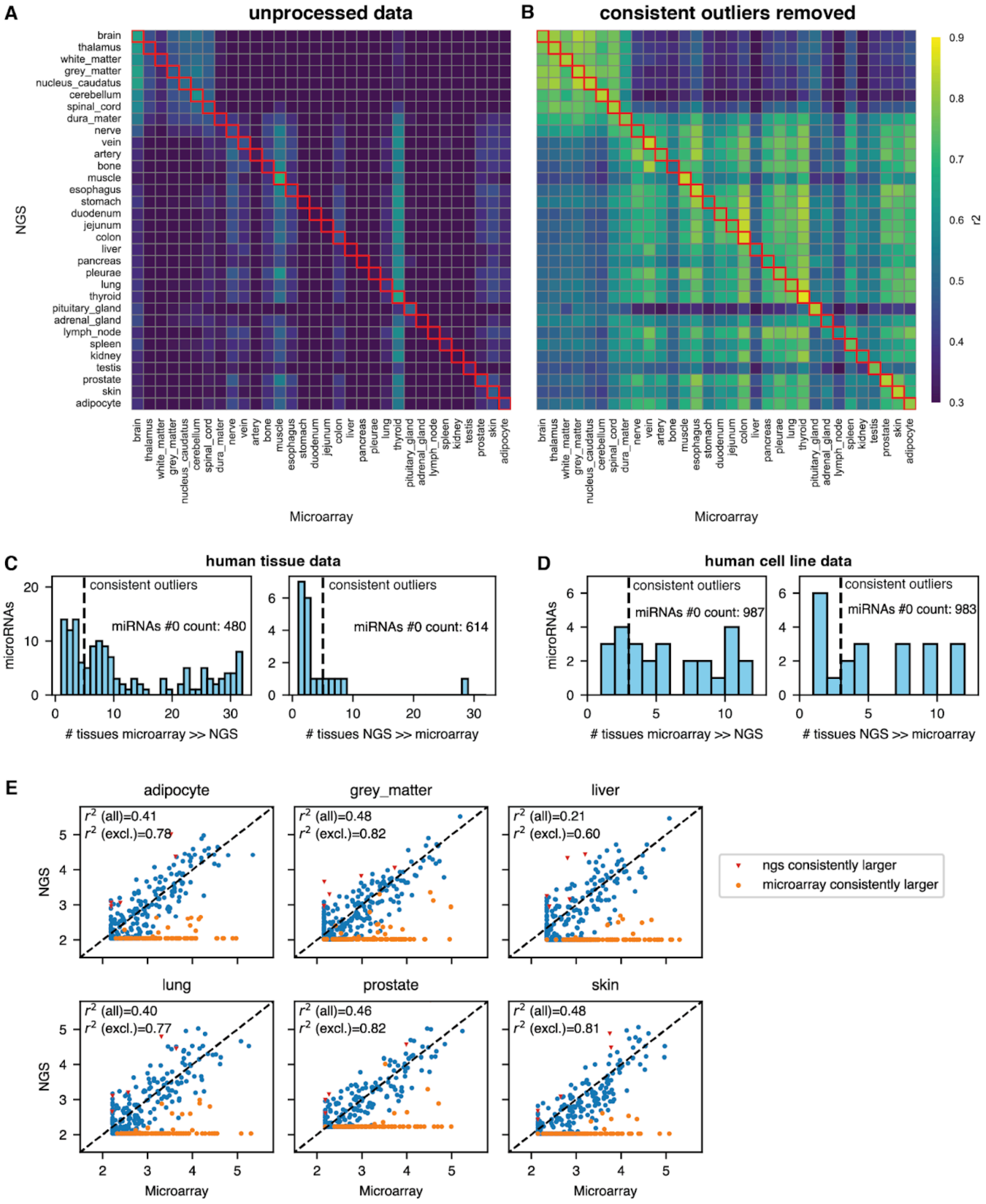


**Fig. S39. Consistent outliers reduce the correlation between microarray and NGS-based human tissue datasets.** (**A**). Correlation between microarray and NGS tissue datasets for all microRNAs that are in either high confidence in miRbase or listed in MirGeneDB. (**B**) Correlation between tissue datasets after removal of consistent outliers. (**C**) Outliers in each tissue are defined as microRNAs with an expression ratio of 10 or more between the two datasets. Consistent outliers are outliers in five or more tissues. The microarray data contains far more outliers with a consistently higher expression than what is seen in the NGS dataset. (**D**) Outliers as in (C) but for expression data for our measured cell lines. For the cell line data, the outlier patterns are more symmetric between the two data types. (**E**) Expression levels for microRNAs in the two datasets for six chosen tissues. MicroRNAs that are consistently higher in one of the two datasets are highlighted. The correlation values are given either for all microRNAs or with consistent outliers excluded.

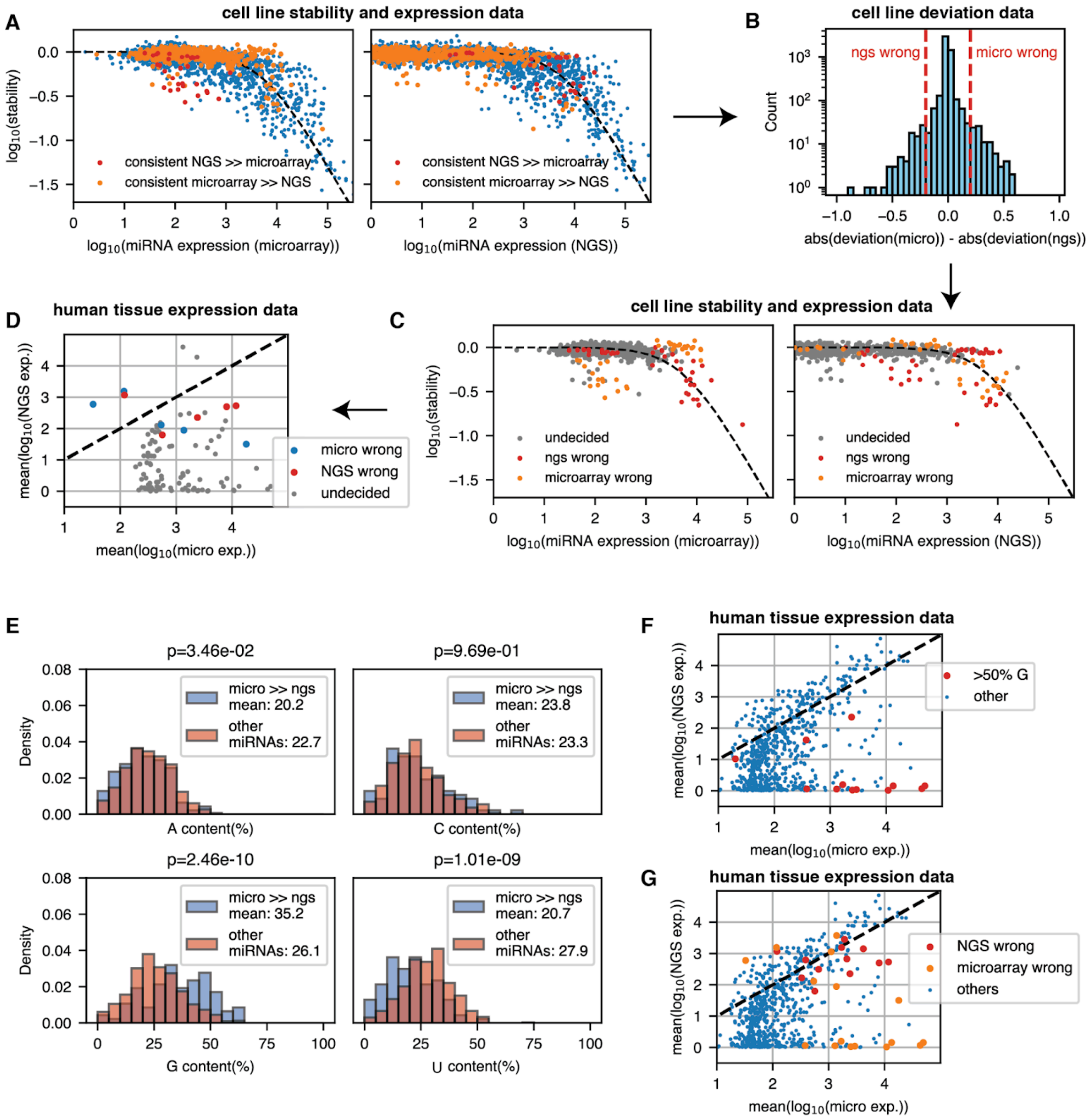


**Fig. S40. Merging procedure for human tissue datasets.** (A)-(D): Decision procedure for consistent tissue outlier miRNAs identification by comparison with cell line stability data. (**A**) Consistent outlier miRNAs identified in the tissue NGS and microarray data as measured in our cell line expression and stability data. As can be seen in the top left plot, some outliers in which NGS data is consistently much larger in the tissue data are also underpredicted by microarray data for the cell lines. (**B**) Distribution of the difference in absolute deviation between the measured and predicted $\log_{10}$ stabilities in cell lines for NGS and microarray data for all measured miRNAs. A much larger deviation for one or the other in a given cell line is interpreted as a sign of bias. (**C**) Cell line expression and stability of miRNA targets for miRNAs that were called as correct in one of the tissue datasets. The decision was made based on the deviation in (B) (Methods). (**D**) Mean of the log10 expression across all tissues for microRNAs that were either called as correct in one of the two datasets or left undecided. Most microRNAs, especially those with high microRNA expression and no NGS expression, were left undecided because they are not expressed in any of the cell lines according to both cell line expression datasets. (**E**) Composition bias for microRNAs that are much more highly expressed in the microarray data. Most notably, these microRNAs have a higher G and lower U content, in line with an earlier study by Backes *et al.* (*82*). P-values were calculated using a two-sided Mann-Whitney U test. (**F**) The most biased miRNAs often have a very large G content of over 50%. Backes et al. (*82*) observed the same bias and confirmed that these microRNAs are also measured as low expression by RT-qPCR, which could indicate that miRNAs with very high G content might be systematically wrong in microarray data. (**G**) Mean expression for miRNAs in the two datasets. MicroRNAs that were found to be likely incorrect in one of the two datasets are highlighted.

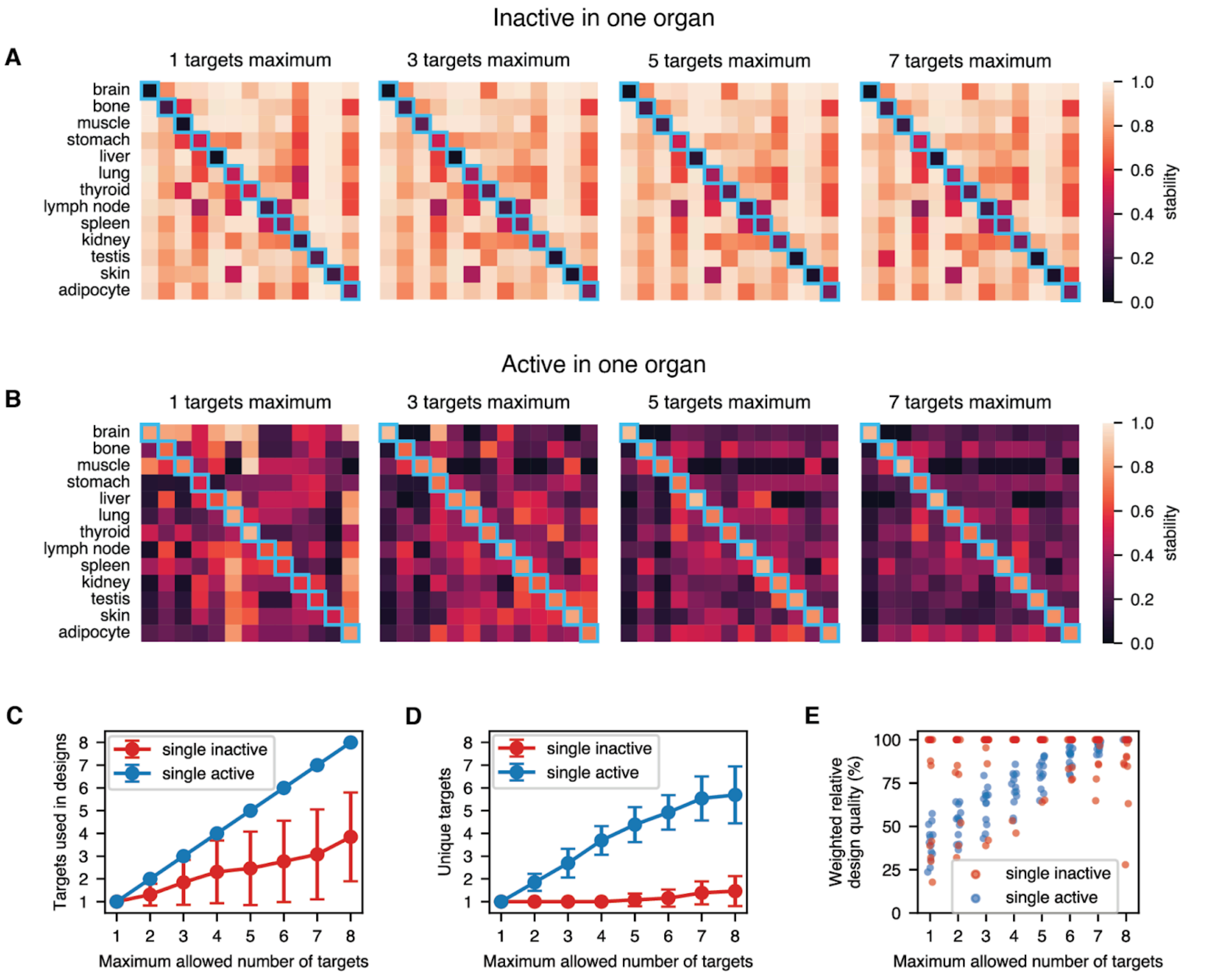


**Fig. S41. The impact of the maximum allowed target site number on the predicted performance of designs targeting human tissues.** We generated designs with a maximum of between 1 and 8 target sites. The design algorithm was also given the option of using fewer than the maximum number of target sites. The designs either (**A**) eliminate expression in or (**B**) constrain expression to a single organ. In the main text, we show designs with up to six target sites. The behavior of the algorithm differs drastically between the two design objectives. (**C**) Number of targets used by the design algorithm. For inactivity in a single organ, the optimum number of targets is often fewer than are allowed. For activity in a single organ, all available targets are used for every design without exception. (**D**) Unique targets per design. Inactivity in a single organ is overwhelmingly achieved by a single type of miRNA target site. Activity in a single organ is achieved by combining many different target site types. (**E**) Weighted design quality (inverse mse) by the number of allowed target sites. Every dot represents a design targeting a different organ. The design quality is shown relative to the highest achieved quality design generated for a specific target organ across all allowed numbers of target sites. For inactivity in a single organ, a design performance close to the maximum is already achieved by a single target site in most cases. For activity in a single organ, the performance keeps improving with the allowed number of target sites. Note that there are organs for which inactivity in a single organ is also improved by up to six target sites. In those cases, this usually means repeating the same target site six times.

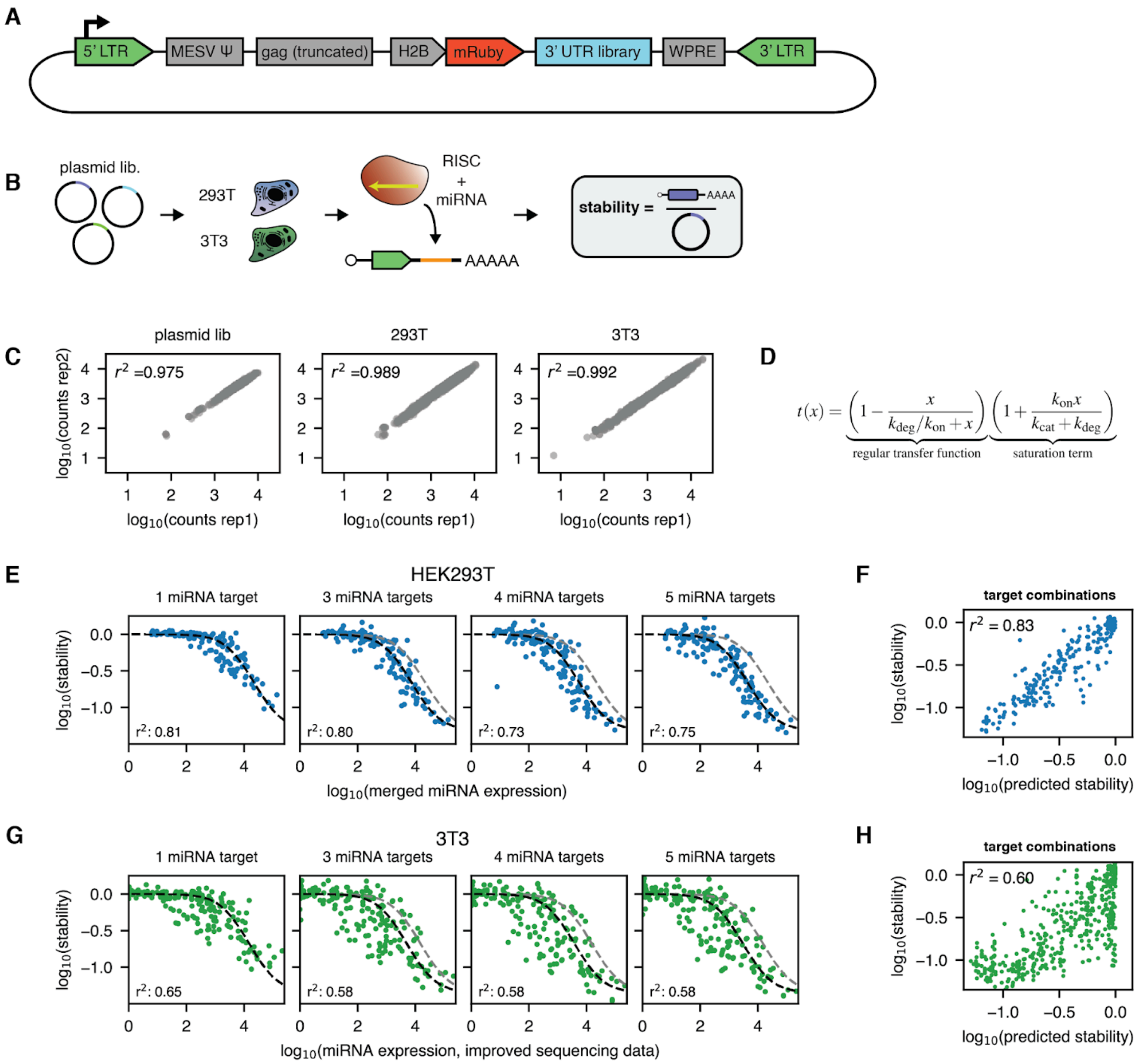


**Fig. S42. Design of the MSCV reporter construct and its measurement in cell lines.** (**A**) Design of the reporter construct. (**B**) The experiments in HEK293T and 3T3 cells were performed using the construct in (A) as a plasmid for a ratiometric stability assay. (**C**) Correlation of counts for replicates for the plasmid library and the stability assays in HEK293T and 3T3. (**D**) Transfer function with a saturation term used to fit expression - stability curves. (**E**) Stability versus miRNA expression data (bias corrected, merged microarray and improved sequencing data) in HEK293T for different numbers of target repeats. The gray dashed lines show predictions for a single target site. Constructs were subset to those with identical miRNA target sites as for human miRNAs for the first 19nt. (**F**) Measured versus predicted stability in HEK293T for combinations of target sites using conditions as in (E). (**G**) Stability versus miRNA expression data (improved sequencing data only) in 3T3 for different numbers of target repeats. The gray dashed lines show predictions for a single target site. (**H**) Measured versus predicted stability in 3T3 for combinations of target sites using conditions as in (G).

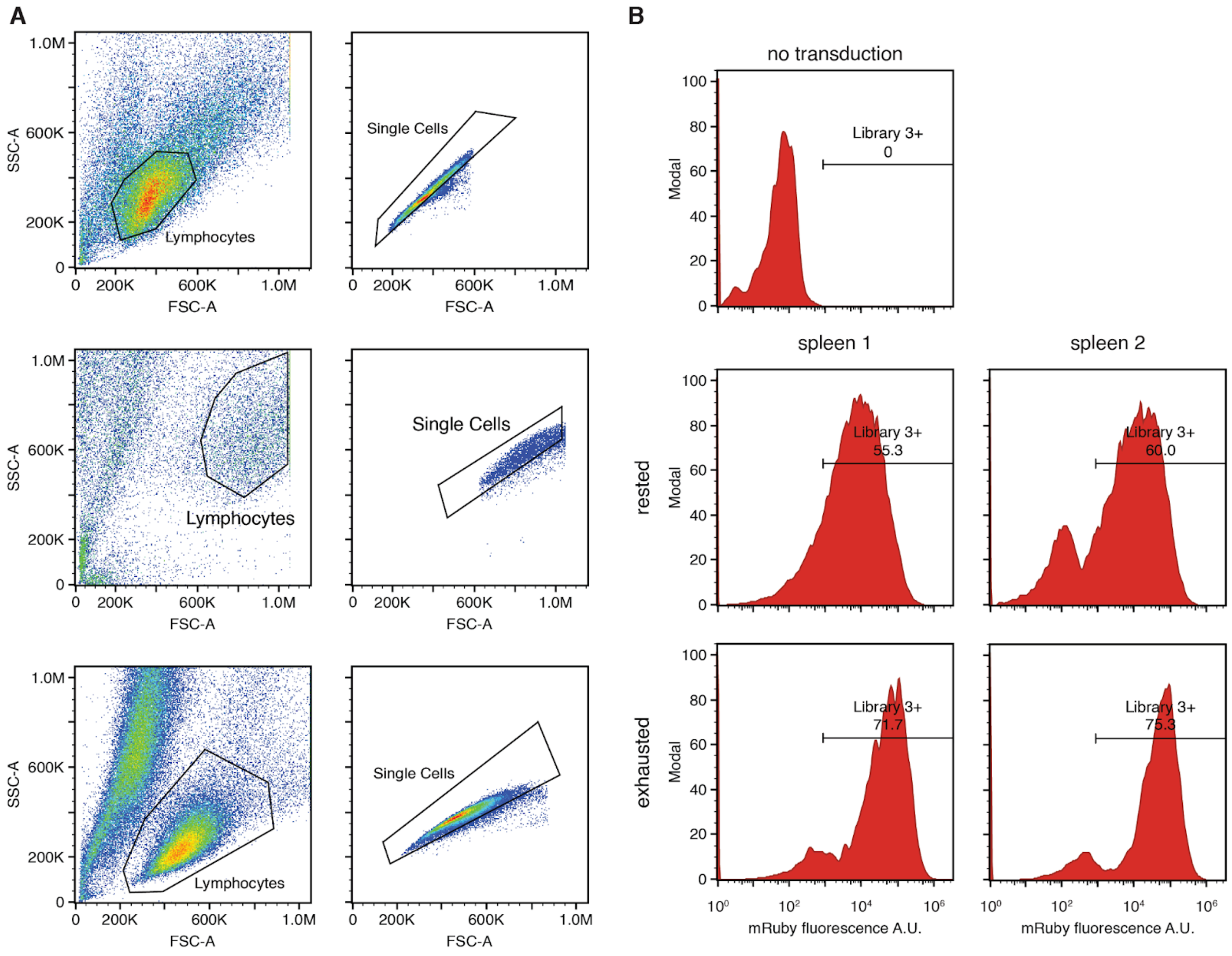


**Fig. S43. Flow cytometry of rested and exhausted CD8 T cell libraries.** (**A**) Live lymphocyte (left) and single cell (right) gating for the first CD8+ T cell experiment rested (top row) and exhausted conditions (middle row), and the second CD8+ T cell experiment all conditions (bottom row). (**B**) Histograms showing transduction efficiency of Library 3 virus in the first CD8+ T cell experiment. All histograms show only events within the Single Cells gate in (**A**, top and middle rows). The values associated with each histogram gate represent the percentage of cells successfully transduced with Library 3 as marked by expression of the mRuby fluorescent protein. Transduction results are displayed for two biological replicates (spleen 1 left; spleen 2 right).

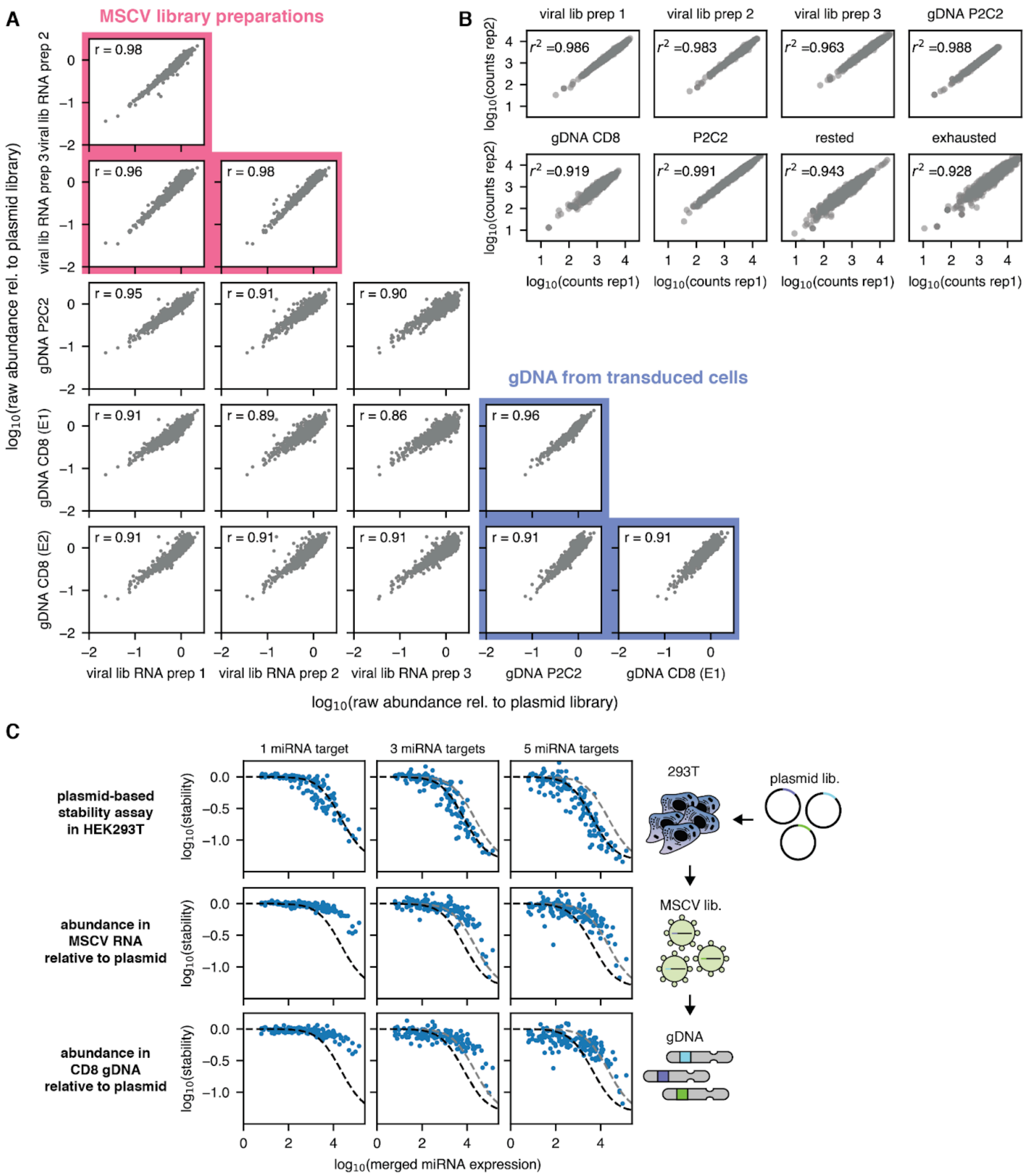


**Fig. S44. HEK293T miRNA regulation determines MSCV library construct abundance.** (**A**) Count correlations for sequencing libraries from different MSCV viral preps and integrations in P2C2 and CD8 gDNA. E1 and E2 refers to the first set of experiments with two conditions and the second set of experiments with four conditions, respectively. (**B**) Count correlations between two replicates for the sequencing libraries of experiment 1. (**C**) Stability values (calculated relative to the abundance in the plasmid library) versus merged microRNA expression in HEK293T for different numbers of target sites. The topmost plot is for mRNA for a plasmid transfection of only the library in HEK293T, the middle plot is for MSCV virus RNA purified from HEK293T, and the bottom plot is for constructs integrated into CD8 gDNA after MSCV transduction.

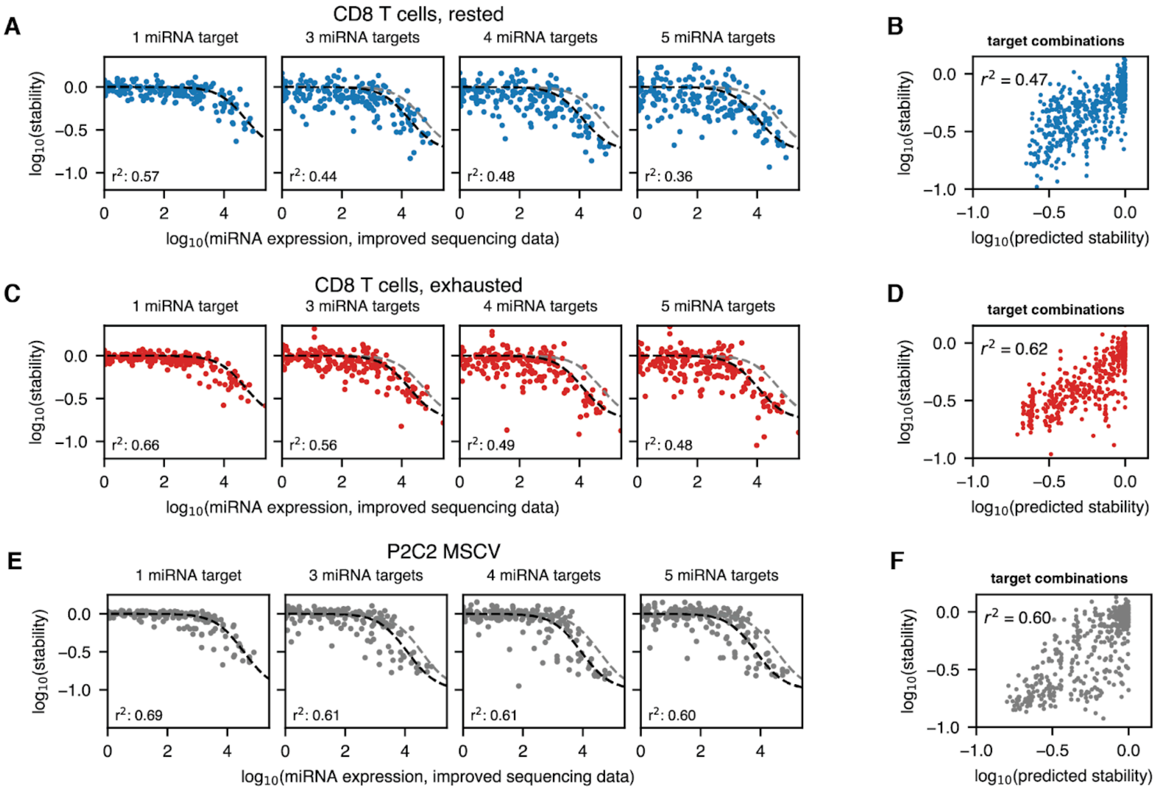


**Fig. S45. Transfer function fits for rested and exhausted T cells and P2C2 cells.** (A), (C), (E): Stability versus miRNA expression data (improved sequencing data) in (**A**) rested CD8 T cells (**C**) exhausted CD8 T cells and (**E**) P2C2 cells for different numbers of target repeats in MSCV transduction experiments. The gray dashed lines show predictions for a single target site. (B), (D), (F): Measured versus predicted stabilities in (**B**) rested CD8 T cells (**D**) exhausted CD8 T cells and (**F**) P2C2  for combinations of target sites.

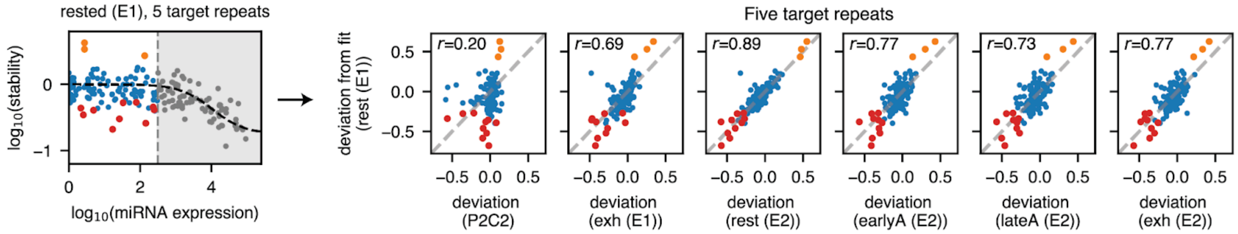


```
mmu-miR-365-3p     UAAGGAUUUUUAGGGGCAUUA
mmu-miR-129-1-3p   UACUUUUUGGGGTAAGGGCUU
mmu-miR-129-2-3p   UGCUUUUUGGGGTAAGGGCUU
```

**Fig. S46. Outliers from the transfer function in CD8 T cells are highly consistent across conditions and experiments.** We identified outliers from the transfer function (orange: highly stable, red: highly unstable) for which the corresponding miRNAs were not expressed in a first experiment (two biological replicates) with rested CD8 T cells. The deviation from the fit for these same target sites was highly consistent across T cell conditions (rested, early activated, late activated, and exhausted) and experimental runs (E1, E2). The target sites of the three highly stable outliers contain highly similar UUUUU(A)GGGG motifs.

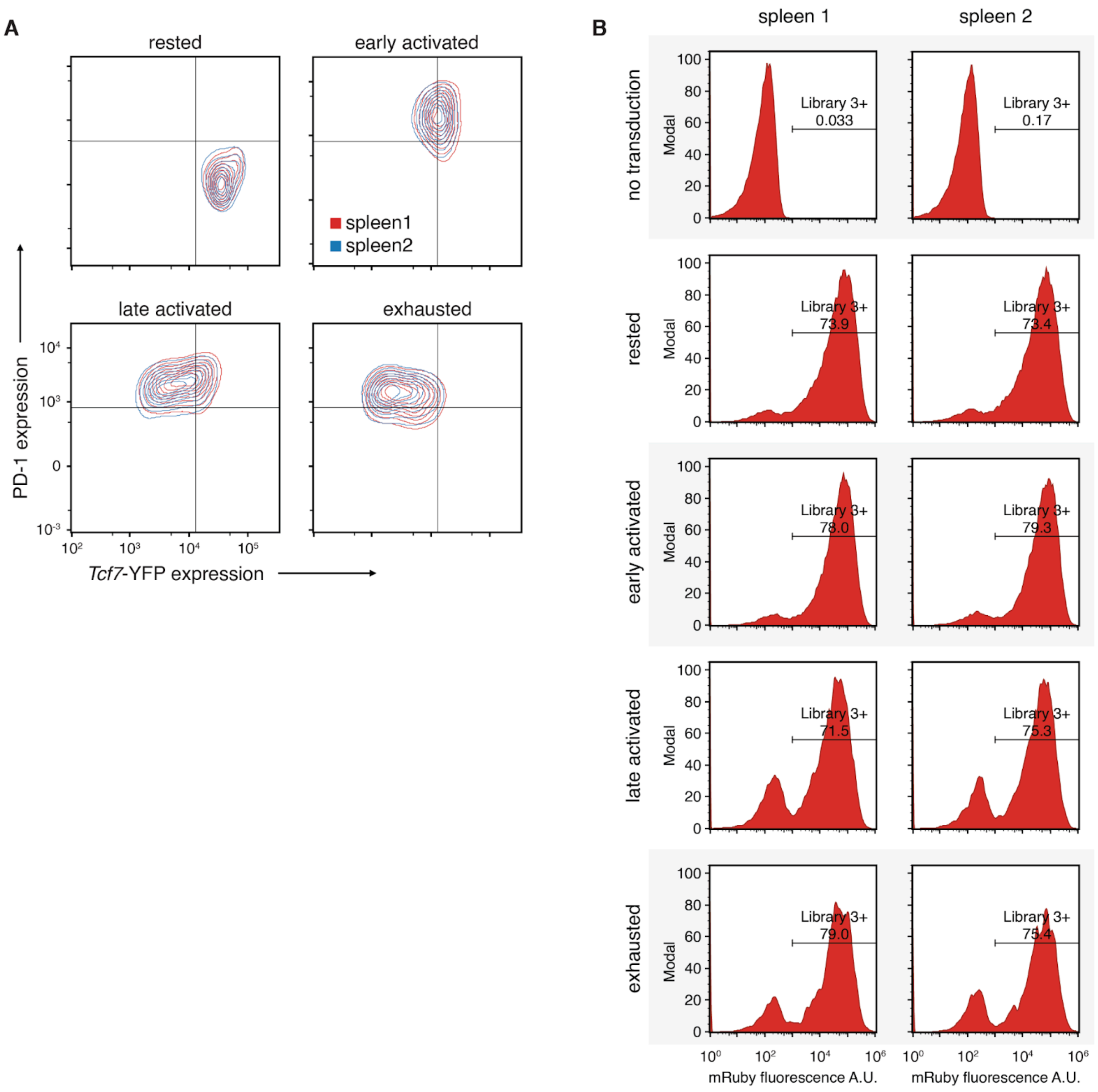


**Fig. S47. Flow cytometry of rested, early activated, late activated, and exhausted CD8 T cell libraries.** (**A**) Flow cytometry measurements of PD-1 vs. *Tcf7*-YFP expression for four different T cell culture conditions. (**B**) Histograms showing transduction efficiency of Library 3 virus in the second CD8+ T cell experiment. All histograms show only events within the Single Cells gate in (**Fig. S47A**, bottom row). The values associated with each histogram gate represent the percentage of cells successfully transduced with Library 3 as marked by expression of the mRuby fluorescent protein. Transduction results are displayed for two biological replicates (spleen 1 left; spleen 2 right).

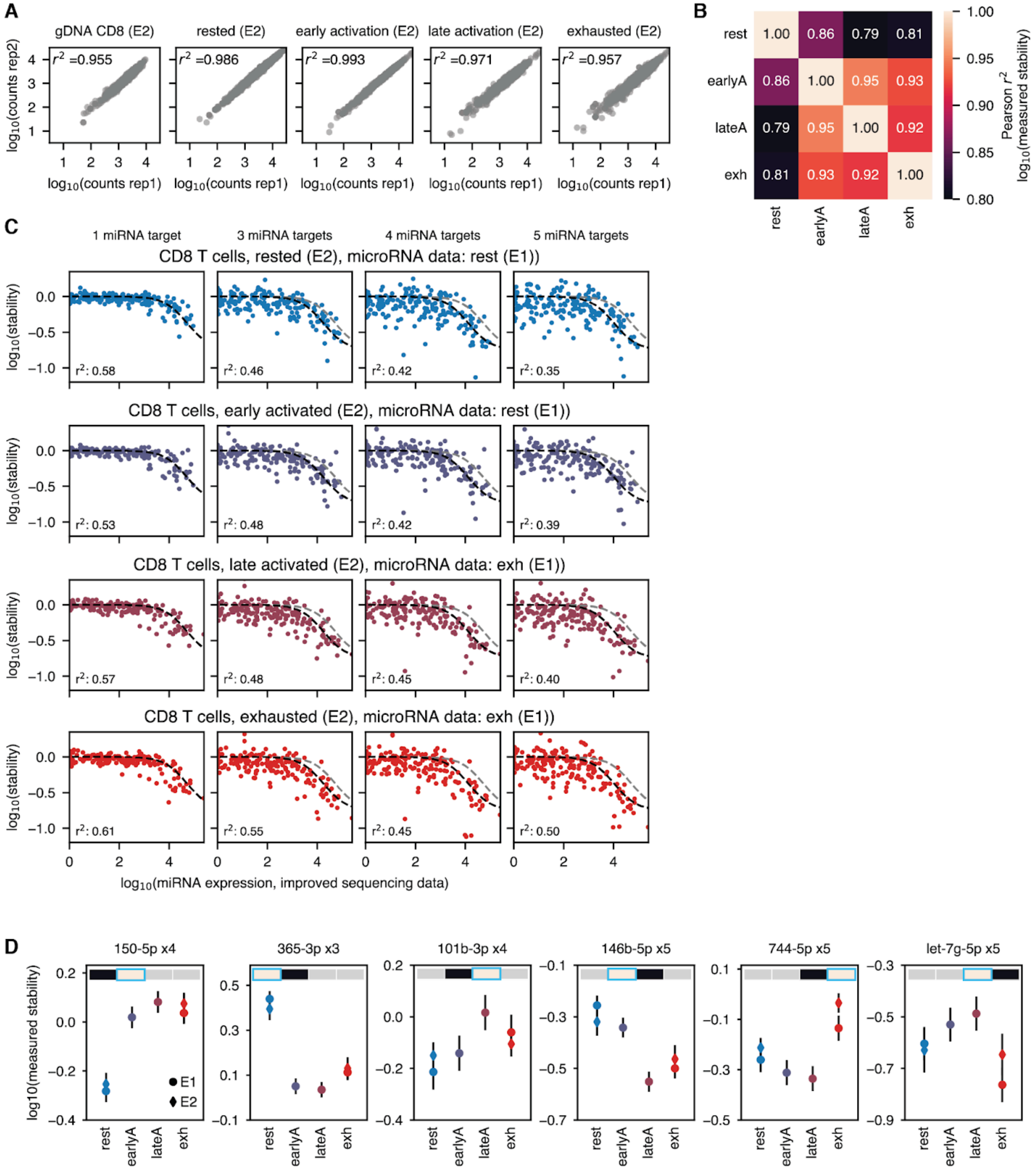


**Fig. S48. Stability data for rested, early activated, late activated, and exhausted CD8 T cells.** (**A**) Correlation of counts for replicates for stability assays across different CD8 T cell conditions. (**B**) Pearson correlation of stability values in different T cell states across the entire library. (**C**) Stability versus miRNA expression data (improved sequencing data) in different T cell states for varying numbers of target repeats in MSCV transduction experiments. The gray dashed lines show predictions for a single target site. The used microRNA expression data was taken from experimental campaign 1. (**D**) Target repeats that most clearly distinguish adjacent T cell states (e.g., low stability and rested and high stability in early activated CD8 T cells for the leftmost plot.

### Supplementary Text

#### The transfer function for predicting the stability an mRNA containing a single microRNA target site

Here, we derive the transfer function for predicting the stability of an mRNA with concentration $m$ containing a single target site for a microRNA with concentration $x$. We assume that $x$ is not appreciably depleted by target binding. In the absence of a microRNA target site, $m$ is degraded at a rate $k_{\text{deg}}$ and produced at a rate $p$:

$$\dot{m} = p - k_{\text{deg}} m$$

with a resulting steady steady concentration of

$$m_0 = \frac{p}{k_{\text{deg}}}.$$

Let $m_{\text{free}}$ be the concentration of free mRNA and $m_{\text{bound}}$ be the concentration of mRNA bound by the microRNA. Assuming irreversible binding this results in

$$\frac{\mathrm{d}}{\mathrm{d}t} m_{\text{free}} = p - k_{\text{deg}} m_{\text{free}} - k_{\text{on}} x m_{\text{free}} \quad (1)$$

$$\frac{\mathrm{d}}{\mathrm{d}t} m_{\text{bound}} = -k_{\text{deg}} m_{\text{bound}} + k_{\text{on}} x m_{\text{free}} - k_{\text{cat}} m_{\text{bound}} \quad (2)$$

for a catalytic rate $k_{\text{cat}}$ of the RISC. In the steady state and with $c_1 = k_{\text{deg}}/k_{\text{on}}$, equation 1 can be rewritten as

$$0 = m_0 - m_{\text{free}} - \frac{x}{c_1} m_{\text{free}} \quad (3)$$

$$\Leftrightarrow \quad \frac{m_0}{m_{\text{free}}} = 1 + \frac{x}{c_1} \quad (4)$$

$$\Leftrightarrow \quad m_{\text{free}} = m_0 \frac{1}{1 + x/c_1}. \quad (5)$$

Under the simplifying assumption that degradation by the RISC is very fast compared to its binding ($m \approx m_{\text{free}}$), the stability of a microRNA-regulated mRNA relative to its baseline stability is given by

$$t(x) = \frac{m}{m_0} = 1 - \frac{x}{c_1 + x}, \quad (6)$$

which is the transfer function used in Fig. 1 to Fig. 4 of the main text.

When a finite catalytic rate $k_{\text{cat}}$ of the RISC is explicitly taken into account, the bound mRNA fraction is given by

$$m_{\text{bound}} = \frac{k_{\text{on}}}{k_{\text{cat}} + k_{\text{deg}}} x m_{\text{free}},$$

which together with equation 5 yields an expression for the total mRNA concentration:

$$\frac{m_{\text{total}}}{m_0} = \frac{m_{\text{bound}} + m_{\text{free}}}{m_0} = \left(1 - \frac{x}{c_1 + x}\right)\left(1 + \frac{x}{c_2}\right) \quad (7)$$

$$c_2 = \frac{k_{\text{cat}} + k_{\text{deg}}}{k_{\text{on}}}. \quad (8)$$

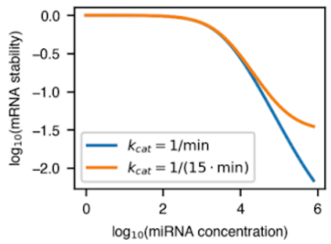


The behavior of the transfer function for small and large catalytic rates. We assume a baseline stability of 8 hours in line with our time course stability assay.

For large microRNA concentrations, the ratio between the mRNA concentrations without and with microRNA regulation in equation 7 thus tends to

$$\frac{m_0}{m_{\text{total}}} \xrightarrow{x \gg c_1, c_2} \frac{c_2}{c_1} = \frac{k_{\text{cat}} + k_{\text{deg}}}{k_{\text{deg}}} = 1 + \frac{t_{1/2,0}}{t_{1/2,\text{cat}}}.$$

This is the maximum loss of stability by binding of a single miRNA. For our first reporter
2148 construct, we found that this term is not necessary even for repeats of six targets sites for

highly expressed microRNAs. Martinez et al.(10.1101/gad.1187904) estimated the *in vitro* half-life of human RISC-bound RNA $t_{1/2,\mathrm{cat}}$ to be approximately one minute, while our $t_{1/2,0}$ is 8 hours. This would imply a maximum stability ratio of 480, which is substantially smaller than the smallest values measured in our libraries. Note that we here measure stability: If an mRNA with a cleaved poly(A)-tail can nevertheless still produce a small amount of protein, a maximum effect of miRNA in the 3'UTR is compatible with our results.
For our second reporter construct, we found this term useful for improving model fit. It is not obvious why these particular reporter construct would differ in this respect. While derived above for a finite catalytic rate of the RISC after binding, essentially any process that generates a subset of the reporter construct resistant to microRNA degradation would have the same effect. For example, if the second reporter is much slower to be exported from the nucleus, this could cause the observed difference.

#### The dependence of the transfer function on total microRNA concentrations and the baseline stability

The transfer function

$$t(x) = \frac{m}{m_0} = 1 - \frac{x}{k_{\mathrm{deg}}/k_{\mathrm{on}} + x}$$

only has a single parameter, namely the ratio between the baseline degradation rate and the on-rate of the RISC. We initially fit $c_1 = k_{\mathrm{deg}}/k_{\mathrm{on}}$ ($\approx 10^{3.7}$ for human cell lines and our reporter construct) as a global parameter for all cell lines. Different total microRNA concentration can be taken into account by a scaling factor $s$ for the total microRNA concentration $x$, i.e., $x_{\mathrm{scaled}} = s \cdot x$ for normalized concentrations $x$.
However, $c_1$ also depends on the baseline degradation rate $k_{\mathrm{deg}} \propto 1/m_0$. When comparing context sequences with different baseline stabilities in one cell line, or a single context sequence with cell type-specific stability, $c_1$ also needs to be adjusted for the change in $k_{\mathrm{deg}}$. A factor $f$ change in stability $\tilde{m}_0 = f \cdot m_0$ thus leads to

$$\begin{aligned}\frac{\tilde{m}}{\tilde{m}_0} = \tilde{t}(x) = 1 - \frac{x}{\tilde{k}_{\mathrm{deg}}/k_{\mathrm{on}} + x} &= 1 - \frac{x}{\frac{1}{f}k_{\mathrm{deg}}/k_{\mathrm{on}} + x} \\ = 1 - \frac{f \cdot x}{k_{\mathrm{deg}}/k_{\mathrm{on}} + f \cdot x} &= 1 - \frac{\tilde{x}}{k_{\mathrm{deg}}/k_{\mathrm{on}} + \tilde{x}}.\end{aligned}$$

An increase in baseline stability by a factor of $f$ therefore has the same effect as an increase in the microRNA concentration by a factor of $s$.

In Figure 1D of the main text, we plot the relative stability predicted by the transfer function for microRNA target sites in different contexts. In our measurement data, stabilities are normalized relative to the stability of our main context. The transfer function parameter $c_1$ has also been fitted for the main context stability. For each context sequence $s$ we also the measure baseline stability $m_{s0}$. If $t_{\mathrm{main}}(x)$ is the transfer function for the main context, the transfer function $t_s(x)$ for microRNA targets in a new context $s$ is thus given by

$$\begin{aligned}\frac{m_s}{m_{s0}} = t_c(x) = 1 - \frac{x}{k_{\mathrm{deg,\,s}}/k_{\mathrm{on}} + x} &= 1 - \frac{x}{\frac{m_{\mathrm{main,\,0}}}{m_{s0}} k_{\mathrm{deg,\,main}}/k_{\mathrm{on}} + x} \\ = 1 - \frac{\frac{m_{s0}}{m_{\mathrm{main,\,0}}} \cdot x}{k_{\mathrm{deg,\,main}}/k_{\mathrm{on}} + \frac{m_{s0}}{m_{\mathrm{main,\,0}}} \cdot x} &= t(x_s),\end{aligned}$$

with $x_s = m_{s0}/m_{\mathrm{main,\,0}}$. To account for different baseline contest stabilities, we therefore both divide the measured stability for each microRNA in a new context and rescale the microRNA concentration by the measured context baseline stability $m_{s0}$. Note that the baseline stability for a single context can differ between cell lines.

#### The additive model for microRNA regulation of multiple target sites.

We start with the same equation as above but for multiple independent miRNAs $x_i$ for the target sites $t_1, t_2, \ldots, t_n$:

$$\frac{\mathrm{d}}{\mathrm{d}t} m_{\mathrm{free}} = p - k_{\mathrm{deg}} m_{\mathrm{free}} - \sum_{i=0}^{n} k_{\mathrm{on}} x_i m_{\mathrm{free}}$$

Assuming instant degradation by the RISC and using the same procedure as above, this yields the following transfer function

$$t(x_1, x_2, ..., x_n) = \frac{m}{m_0} = 1 - \frac{\sum_{i=0}^{n} x_i}{k_{\text{deg}}/k_{\text{on}} + \sum_{i=0}^{n} x_i},$$

which is exactly the same transfer function as for a single site except that the concentrations of attacking microRNAs are summed before the transfer function is applied.
